# The *Trypanosoma cruzi* kinetoplast DNA minicircle sequences transfer biomarker of the multidrug treatment of Chagas disease

**DOI:** 10.1101/2021.12.16.473091

**Authors:** Alessandro O Sousa, Clever Gomes, Adriana A Sá, Rubens J Nascimento, Liana L Pires, Ana M Castro, Francisco Moreno, Antonio RL Teixeira

## Abstract

**Background:** The *Trypanosoma cruzi* infection renders the transfer of the mitochondrion kinetoplast DNA minicircle sequences into the host’s genome. The Aves are refractory to the infection, but chicks hatched from the *T. cruzi* inoculated eggs integrate the DNA minicircle sequences into the germ line cells. Rabbits, mice and chickens with the minicircle sequences mutations develop the Chagas cardiomyopathy and the DNA transfer underpins the heart disease.

**Methodology:** The PCR with the specific primer sets revealed the Protist nuclear DNA and the kinetoplast DNA in the agarose gels bands probed with the radiolabel specific sequences from tissues of the *T.* c*ruzi-*infected rabbits and of the mice. A target- primer TAIL-PCR amplification employing primer sets from the chickens, rabbits and mice, in combination with primer sets from the the *T. cruzi* kinetoplast minicircle sequences was used. This approach led us to disclose the integration sites of the kinetoplast DNA biomarker, then, used to monitor the effect of multidrug treatment of the *T. cruzi* infected mice.

**Principal findings:** The Southern hybridization, clone and sequence of the amplification products revealed the DNA minicircle sequences integrations sites in the LINE transposable elements. An array of inhibitors of eukaryote cells division was used to arrest the DNA transfer. It was shown that nine out of 12 inhibitors prevented the kinetoplast DNA integration into the macrophage genome. The multidrug treatment of the acutely *T. cruzi-*infected mice with Benznidazole, Azidothymidine and Ofloxacin lessened circa 2.5-fold the rate of the minicircle sequences integrations in the mouse genome and inhibited the rejection of the target heart cells.

**Conclusion and significance:** The *T. cruzi* mitochondrion kinetoplast minicircle sequences transfer driven pathogenesis of Chagas disease is an ancient Cross-Kingdom DNA phenomenon of evolution and, therefore, paradigm research with effective purposing inhibitors is needed.

**Authors summary:** Chagas disease is considered the main cause of human heart failure in the Western Hemisphere. The treatment of the clinically manifested Chagas heart disease is considered unsatisfactory. Perhaps the most important problem in the field of Chagas disease is determination of the pathogenesis of the target heart cells lysis. We showed the transfer of the *T. cruzi* kDNA minicircle sequences into the genome of rabbits and mice, and to Bird refractory to the infections. The inoculation of a few *T. cruzi* in the fertile chicken eggs renders the kDNA sequences integration in the stem cells. Interestingly, the chicks that hatched retain the kDNA and develop the Chagas-like cardiomyopathy indistinguishable to that in the rabbits and mice. This result prompted the multidrug treatment of the Chagas heart disease with inhibitors of the eukaryotic cells division. We showed that nine out of 12 inhibitors prevented the transfer of the kDNA mutations into the macrophage genome, and that the treatment of the acutely *T. cruzi*-infected mice with Benznidazole + Ofloxacin + Azidothymidine lowered circa 2.5-fold the rate of the mutations in the chromosomes. These findings translated to the pathology showing inhibition of the heart lesions in the treated *T. cruzi-*infected mice. We suggest purposing new inhibitors should be tested to overturning the Chagas heart disease.

## Introduction

The *Trypanosoma cruzi* (Eukaria, Excavata, Euglenozoa, Zoomastigophora) agent of American trypanosomiasis is a Protist flagellate showing a disc-shaped kinetoplast in the symbiotic mitochondrion. An ancient chain of events initiated at 670 million years ago (mya) prompted the joining of prokaryote undulipodia to a protist eukaryotic precursor, possibly, to the ancestral *Trypanosoma gray* recognized in fish [1, 2, 3]. At the Devonian era, 360 mya, wing-insect fed upon existing vascularized plant and initiated a divergent evolution. On the one hand, it gave rise to the tsetse fly (*Glossina palpalis*) salivary transmission of the *Trypanosoma brucei* agent of the African trypanosomiasis, firstly transmitted to the crocodile [1]. On the other, the insectivorous Marsupialia (Chordata, Didelphimorphia), existing since the Permian 245 mya, is the ancient host of the *Trypanosoma cruzi* stercorary flagellate thriving in the hindgut of bloodsucking triatomines (Hemiptera, Reduviid) since the Cretaceous, circa 100 mya. The omnivorous marsupials, anteaters, armadillos, and the mammalians belonging to several classes (edentates, lagomorphs, rodents, carnivores, primates, and chiropterans) are hosts for *T. cruzi.* Birds are refractory to *T. cruzi* infections [4]. The infections were transmitted to over 1,000 species of wild mammal’s host to the *T. cruzi* [1]. Approximately 50 thousand years ago the *Homo sapiens* arrived in the America Continents where the “kissing bugs” sympatric span an extensive geographical area between 42°N in the United States to 43°S in Argentina [2, 3]. In the presence of domesticated mammals and of “kissing bugs” in the household the anthroponotic infections took place.

The American trypanosomiasis was discovered by Doctor Carlos Chagas (Chagas, 1909) in the Brazilian Minas Gerais State hinterland: during the provision of health assistance to a feverish child, he found some flagellates in the child’s blood. In the following weeks, while she recovered spontaneously from the fever, Doctor Chagas awareness of the annoying “kissing bugs” in that region led him to find some flagellates in the triatomines hind gut. Upon injection of the contaminated bug excreta into the mouse, Doctor Chagas and his associates showed the *T. cruzi* trypomastigote in the blood, and the niches of replicative amastigote forms in the host’s tissue cells [5].

The *T. cruzi* infections in humans can be acquired through the insect-bite site in the skin or through the mucosal surfaces contaminated with the triatomine excreta, but the entry of the parasite is aimless [6]. During the last century the human population concentrated in the rural areas of Latin America mainly were victimized and, insofar, the Chagas disease neglected, possibly, because it was believed to affect the poor under the exposure to triatomine bites [7, 8]. The growing evidence shows that the *T. cruzi* infections transmitted by several routes [7–9] streamline the current epidemiologic trends of the people migration, with the convert demography towards the county borough, and spread to the global, cosmopolitan Northern hemisphere [8–20]. The *T. cruzi* infections transmitted by blood transfusion [21–23], by oral route [24], and by accident in hospitals and research laboratories [25, 26]. Moreover, the congenital, sexually transmitted *T. cruzi* infection from males and females to naive mates could play an ongoing pandemic role [6, 27–38, 39].

The early phase *T. cruzi* infections in humans are usually asymptomatic and not perceived by the individual with the flagellates in the blood, and thus he or she, and the family, do not seek for health care (2, 6, 39). However, some oligosymptomatic acute cases refer to fever, headache, malaise, muscle pain, and prostration, and occasional fatal cases with severe acute *T. cruzi* infection show myocarditis and/or encephalitis. In the last three decades an average four fatal acute cases of Chagas disease have been published [40]. The diagnosis of the early phase Chagas disease is achieved, usually, by the direct microscopic demonstration of the *T. cruzi* in the patient’s blood and by the hemoculture [5, 6].

The late phase, long-lasting *T. cruzi* infections run in the absence of clinical symptoms and signs. However, circa one third of the chronically infected humans develop clinical manifestations of disease averaging 40 ± 10 years after onset of infection [7]. The Chagas disease, thus affecting the heart (94.5%) and the digestive system megacolon and megaesophagus (5.5%), and combined multifaceted neuroendocrine syndromes associate to parasympathetic neuronal cells lysis in the chronically infected case [7].

Chagas disease with an associated family history is the most prevalent cause of heart failure in the Western Hemisphere [33, 35, 36], leaving behind orphanhood and an enormous economic burden [6]. The intermediate chronic phase of the infection diagnosed by indirect methods, such as the search for specific anti-*T. cruzi* antibodies [2, 6, 23, 21, 42]. However, the specificity and sensitivity of these serologic methods to the diagnostics of the chronic infections is a matter of concern during the disclaim of blood donators. With this respect, the combination of serologic with nucleic acids assays are recommended for the diagnosis of Chagas disease, affidavit [6, 36, 37, 43].

The prevalence of Chagas disease, in the absence of systematic methods to obtaining epidemiologic data for the meta-analysis, has been assumed by the extrapolation of the information from independent field studies carried out in various counties of the Latin America [2, 23–42]. The systematic, country-wide survey that was conducted in Brazil, during the years 1975-1980, employed the indirect immunofluorescence method for the search of the *T. cruzi* antibody eluted from a drop of blood serum sample collected in filter paper. The elution and storage of the blood samples serologic assay hidden flaws cannot be deemed by using a second immunoassay, because unaccountable cases of the congenitally transmitted *T. cruzi* infections bear immune tolerance in the absence of the specific antibody [43]. Nonetheless, the systematic survey revealed a positive tests ratio of 0.58%, which translated a total of 6 million positive people. At that time the Brazilian population was about one-third of that much in the Latin America countries and, thus, constantly present *T. cruzi* infections was assumed as thrice, that is, to say circa 18 million Chagas people [44]. The nucleic acids assays that reveal the *T. cruzi-*infected immune tolerance family members, showing the sexually transmitted infections [33–36] are not available in most diagnostic facilities and hospitals. In the absence of systematized protocols to obtain data for the meta-analysis the true dimension of the world health problem remains unrecognized, and, therefore, Chagas disease is neglected for over a century.

Interestingly, the Protist *T. cruzi* enclosed aneuploidy in the growing parasitic stages bear extensive genetic variations with biological implications [45]. The protozoan genome average size 50 Mb nuclear DNA (nDNA) includes circa 8000 genes [46–48]. The *T. cruzi* mitochondrion shows a disc-shaped kinetoplast adjacent to the flagellum basal body, which concentrates 25% of the cell total DNA. The kinetoplast contains a few dozen of maxicircles and circa 20 thousand minicircles in a catenated network [49]. The maxicircles are almost identical to the mitochondrial kinetoplast DNA (kDNA) of mammals with the ribosomal RNAs coding sequences [50–52]. The thousand copies of minicircles average size 1.4 kb, each comprising four conserved 238 base pairs (bp) regions interspersed by four variable sequences size 122 bp intercalated sequence blocks – CSB1, CSB2, and CSB3 [51]. The demonstration of the *T. cruzi* kDNA minicircle sequences transfer into the vertebrate host’s genome, showing the *locus-*specific insertion mutation, generates genome growth and evolution, and natural selection with practical biological and medical consequences over time [53].

The pathogenesis of the Chagas disease multifaceted clinical manifestations has been associated to the lifelong parasitic infection and the transfer of the mitochondrion kDNA sequences into the host’s genome [33, 39, 41, 53–55]. The long-lasting *T. cruzi* infection could be an essential condition to a continuous flow of kDNA, and the cumulative mutations over time may induce the genetically driven autoimmune mechanisms leading to the destruction of the target tissues of its host [56, 57]. The *T. cruzi* kDNA minicircle sequences are consistently transferred to the genome of mammals and birds [6]. The kDNA insertional mutations into the host’s genome retrotransposons, 72.8% of which integrated in hundreds of copies to several chromosomes could drive the autoimmune lesions in Chagas disease [56, 57].

The understanding of the pathogenesis of Chagas disease and of the roles played by animal models towards the *T. cruzi* infections translated into the disease clinical manifestations is essential to accomplish the drug treatment objectives [58, 59, 60]. In this regard, the rodent (*Mus musculus*) model highly susceptible to the *T. cruzi* infections are mostly used to monitor the drug killing of the trypanosomes and assessments of a satisfactory treatment of the disease [62, 63]. The rabbit (Chordata, Lagomorpha) and the dog (Carnivora, *Cannis Lupus familiaris*) are relatively resistant to an early infection, but they die of chronic chagas heart disease usually over one year after the acquisition of the *T. cruzi* infection [63–65]. The primate is not considered an experimental research model for the ethical and ecological reasonings [66, 67]. However, the trans kingdom chicken model system refractory to the *T. cruzi* infection is essential to the genetic study towards unravelling the Protist kDNA transfer phenomenon [41, 56, and 57].

Over the last 60 years the lead heterocyclic compounds Benznidazole [N-Benzyl-2-(2-nitro-1H-imidazol-1-yl) acetamide] and Nifurtimox [3-Methyl-4-(5’-nitrofurylidene-amino)-tetrahydro-4H-1,4-thiazine-1,1-dioxide] are recommended for the treatment of the *T. cruzi* infections [58, 59]. The nitro heterocyclics killing of eukaryotic cells implicate the reductase enzymes activity, forming nitro anion radical (R-NO²^-^) and free O²^-^, OH^-^ and H^2^O^2-^ [70, 71]. The nitro-anion binding to macromolecules kills the trypanosomes and any other nucleated cell type. The administration of the nitro heterocyclic may induce severe hypersensitive reactions, dermatitis, polyneuritis, joint and muscle pain, neutropenia, bone marrow depression, paresthesia, and convulsion. The drug toxicity precludes the completion of the full treatment, and the proportion of failure is high [60–64]. In addition, the nitro heterocyclics are considered unsatisfactory because the long-lasting treatment (60 to 90 days) do not clear the parasite from the host’s body [62–69]. Moreover, the nitro heterocyclic free-anion radicals bind to DNA and the resulting DNA adducts associate severe genotoxicity [70–74].

The treatment of Chagas disease requires a cornerstone program for the scientific achievement of its prevention. In the last decades, several multicentric research teams introduced new compounds aiming at the treatment of the *T. cruzi* infections [75–81]. Meanwhile, the nitro heterocyclics Benznidazole and Nifurtimox lead the curtailment of the trypanosomes in the human blood [68, 69]. Herein, we advance the hypothesis the eukaryotic cells metabolic pathways of cell growth and differentiation should be targeted with an array of specific inhibitors. The suggested multidrug treatment could curtail the parasitic load, and to overturning the *T. cruzi* mitochondrion kDNA insertional mutations into the transposable elements of the host’ genome, for preventing the installment of the multifaceted life-threatening clinical-pathological manifestations of the Chagas disease. With this respect, some features of the metabolic pathways of eukaryote cells growth and differentiation check points are reviewed, briefly.

The protein kinase dependent cyclins (Cdk) that phosphorylates serine-threonine protein inducers of the cell growth (G1), synthesis (S), (G2) and mitosis (M) cycle [81–85]. A family of D cyclins responsive to external factors activate the nuclear signaling pathways and transcription of the primary cell cycle response genes [86]. A class of cyclins that binds the Cdk at the S1 phase and are required for the inception of the cell cycle S phase, and other class that binds Cdk required to inducing the cell division. The expression of the cyclins (D1, D2 and D3) activates CDK4 and 6 phosphorylates the proteins in command of the passage of the cell growth (G1) to synthesis (S) phase [86–89]. The mitogen-activated protein kinases (MAPK) playing a role in regulation and expression in eukaryotic cells comprise various cyclin-dependent, serine-threonine protein kinases (MAPKs), glycogen synthase (GSK3) and CDK-like (CDKL) kinases, undergoing biological function cascade of the cell growth and differentiation [90–93]. The serine-threonine kinases phosphorylation activated (MAPKK) enclosing 14 family members identified in the mammalians’ cells [95]. In addition, those cells extracellular signal regulated kinases (ERK1 and 2), c-Jun N-terminal kinase (JNK), and their additional isoforms carrying out protein’s phosphorylation [93–96]. The PI3K classes of lipidic kinases phosphorylates inositol 3’-OH ring. The class I PI3K heterodimers enclose a catalytic (CAT) subunit forming phospholipids [97].

The PI3K receptors with protein kinase activity and the G protein receptors are inducers of autophosphorylation. Its subclasses IA and IB activated, respectively, by protein tyrosine-kinase or by G-protein receptors undergo autophosphorylation to forming phospholipids [98]. Thus, the allosteric activation of second messenger phosphatidylinositol-3,4,5-triphosphate (PI3,4,5-P3), recruiting other serine-threonine signaling proteins such as 3’-phosphoinositide-dependent kinase 1 (PDK1) and Akt-protein-kinase B (PKB) that regulates cell growth and survival. Akt-PKB inactivates pro-caspase and inhibits Forkhead transcription factors to the cell apoptosis, such as Faz-L. Akt/PKB may activate the NO synthesis [99–103]. The protein IKKα that inhibits IKβ and promote the NFKβ that induces cyclin D1 secondary response genes [104]. In addition, the Ras protein in the cell membrane activates PI3K that phosphorylates phosphatidyl inositol 3-phosphate that links to protein kinase-B (PKB) and AKT through specific domains. In addition, the serine-threonine protein kinase C (PKC) isoforms activated by Ca++ released from the endoplasmic reticulum by diacylglycerol (DAG) and inositol 1,4,5-triphosphate (IP3), contributes to the cell growth pathways. Twelve PKC isoforms act upon functional ionic channels, enzymes, and cytoskeletal proteins. The isoforms regulate the enzyme substrates to different signaling pathways to the control of the cell differentiation [105].

The hypothesis to overturning the Chagas disease clinical-pathological manifestations is set forward herein, whereby a multidrug treatment with a lead nitroheterocycle compound, which curtails the parasitic load, should be administered altogether with the drug inhibitors of the main metabolic pathways of the eukaryotic cells growth and differentiation, in order to disclaim some of those *T. cruzi* mitochondrion kDNA minicircle insertional mutations that call out the clinic manifestations of Chagas disease over time. The investigation showed that the treatment of *T. cruzi* infected mice with the lead Benznidazole plus the Ofloxacin and Azidothymidine, respectively, polymerases and reverse transcriptase inhibitors, lessened the lateral transfer of the kDNA into the mouse’s genome and cut short the myocarditis of Chagas disease.

## Results

### Genetic markers of the *Trypanosoma cruzi* infection

Upon entry of *T. cruzi* into the body of the mammalian, the infective trypomastigotes are taken up by mononuclear phagocytes [52, 106–110]. The few intruding flagellate may be destroyed in the phagocytes, but the infection with many of the parasite replicate as amastigotes before returning to the trypomastigote forms that then burst out of the host cell to invade any tissue or cell type. The parasite resistance form’s persistence within a vertebrate is spent hidden from the host immune system inside non-phagocytic cells. Indeed, *T. cruzi* amastigotes can persist for decades in this setting without causing the host significant damage [111]. We observed that the treatment of chronically infected rabbits with a trypanosome killing agent curtailed tissue parasitism, but it did not stop the progressively destructive myocarditis and peripheral nerve system ganglionitis, hallmarks of Chagas disease [112, 113]. What could be sustaining the active destruction of the heart cells? To answer this question, we postulated that some rate of genetic transfer from the parasite to the host genome and that the resulting mutation could explain the autoimmune-driven lesions. In this regard, we decided to search for parasite DNA in the genome of Chagas patients with the *T*. cruzi infections for over 30 years and had recently manifested the clinical symptoms of chronic Chagas heart disease. [61, 113].

We analyzed genomic DNA samples from the blood of 13 Chagas heart disease patients that had shown antibody specific to the *T. cruzi.* These entire samples harbored parasite DNA as judged by PCR amplification with primer for both *T. cruzi* nDNA and kDNA minicircle, showed in the S1 and S2 Tables. Southern hybridizations were performed on genomic DNA from these patients. In a typical example, the DNA sample from Chagas patient 245 revealed 400 bp and 100 bp bands with the kDNA minicircle probe only, whereas digested and catenated minicircles revealed high molecular weight bands of 1.4 kb and larger. (Figure 1). The genetic markers of the *T. cruzi* infection shown in the Figure 1 A and B were used to investigate our postulate that the transfer of the *T. cruzi* kDNA minicircle sequence into the vertebrate’s genome could introduce critical biological host-pathogen interactions, and significant modifications translated to the clinical manifestations of Chagas disease.

**Figure 1.**
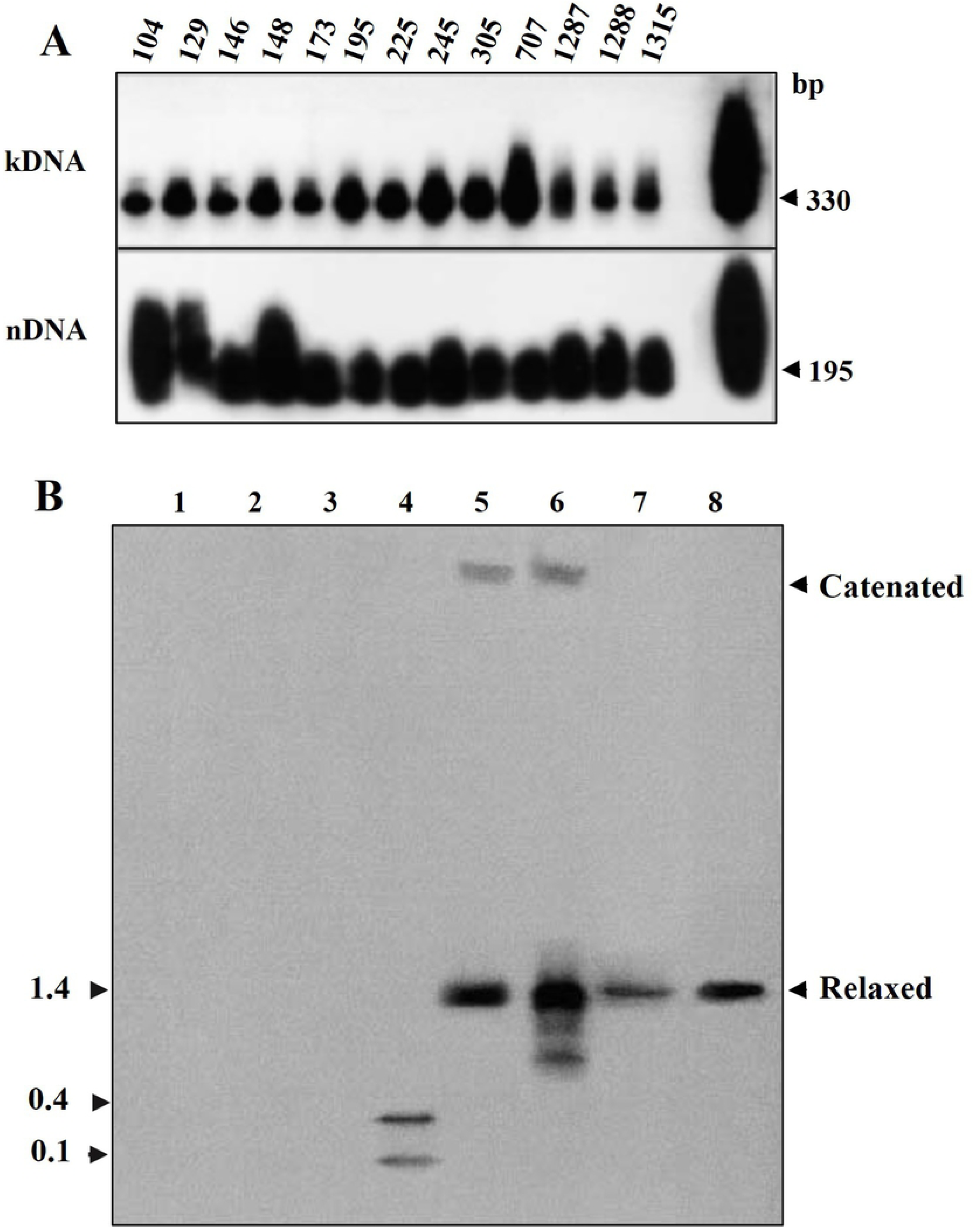
Genetic markers of the *Trypanosoma cruzi* infections and genomic kDNA integration in chronic Chagas patients. The *T. cruzi* kDNA 330 bp and the nDNA 195 bp bands obtained by the PCR amplifications with specific primer sets, separation through a 1 % agarose gel, the DNA stained with ethidium bromide, blotted, and probed with the specific kinetoplast DNA (kCR) and nuclear DNA (nCR) oligonucleotides (See Methods). **A)** Evidence of the persisting *T. cruzi* infections. PCR amplification products obtained from template DNA of 13 Chagas heart disease patients and specific sets of kDNA (S35/S36; upper panel) and nDNA (Tcz1/Tcz2; lower panel) primers, which hybridized with the internal specific probes kCR and nCR, respectively, on blots of 1% agarose gels. The size markers indicate the minimal unit of amplification in each case (See methods). **B)** Southern hybridization of integrated kDNA revealed by a minicircle (kCR) probe. A 0.7% agarose gel was used to analyze undigested DNA or *Eco*RI and *Bam*HI digested DNA from the blood of Chagas disease patient 245. After size separation through agarose a 0.7% gel, the DNA was stained with ethidium bromide, blotted, and probed with the kCR oligonucleotide. DNA from a non-infected human donor and purified kDNA were included: 1) Control; 2) control, digested; 3) Chagas patient; 4) Chagas patient, digested; 5) Control + kDNA; 6) kDNA; 7) kDNA, digested; 8) Control + kDNA, digested. DNA from a noninfected human donor and purified kDNA were included as controls. The predicted sizes of catenated and relaxed or linear minicircles are indicated.

The genome of the *T. cruzi* contains the nuclear DNA and the mitochondrial kinetoplast DNA. The PCR amplification with specific primer sets S35/S36 and Tcz1/Tcz2 products, separate in 0.8% agarose gels, blot and Southern hybridization with the internal probes showed, respectively the 330 bp kDNA and the 195 bp nDNA bands. The search for the kDNA and the nDNA markers of the *T. cruzi* infections in the blood of chronic Chagas disease patients showed the amplifications products of the nDNA and of the kDNA and Southern hybridization with the [α^32^P]-dATP specific nDNA (nCR) and kDNA (kCR) internal probes (see Methods) yielded identical 195 bp nDNA and 330 pb kDNA bands. The primers and probes employed to the PCR-amplification assays that revealed the genetic markers of the *T. cruzi* infections, showed in the S1 and S2 Tables. To proceed the investigations, we bode to clarify whether the naked *T. cruzi* kDNA minicircle could integrate into the phagocyte genome.

### Searching for the naked *T. cruzi* kDNA sequences in eukaryote cell’s genome

The macrophage uptake and processing of intruder macromolecules is a defense mechanism, but there is a lack of information on the fate of the naked kDNA sequences taken in by the phagocyte. The assessment of the role of macrophages in the engulfment and digestion of nucleic acids sequences from different sources: CAT gene, cloned kDNA, crude *T. cruzi* kDNA, and macrophage kDNA sequences, showed in Figure 2. The lipofection-treated sequences uptake by the macrophages harvested at the set points shows the CAT sequence cleared out at day the 14^th^, whereas the *T. cruzi* kDNA sequences wasted out at the day 21^st^.

**Figure 2.**
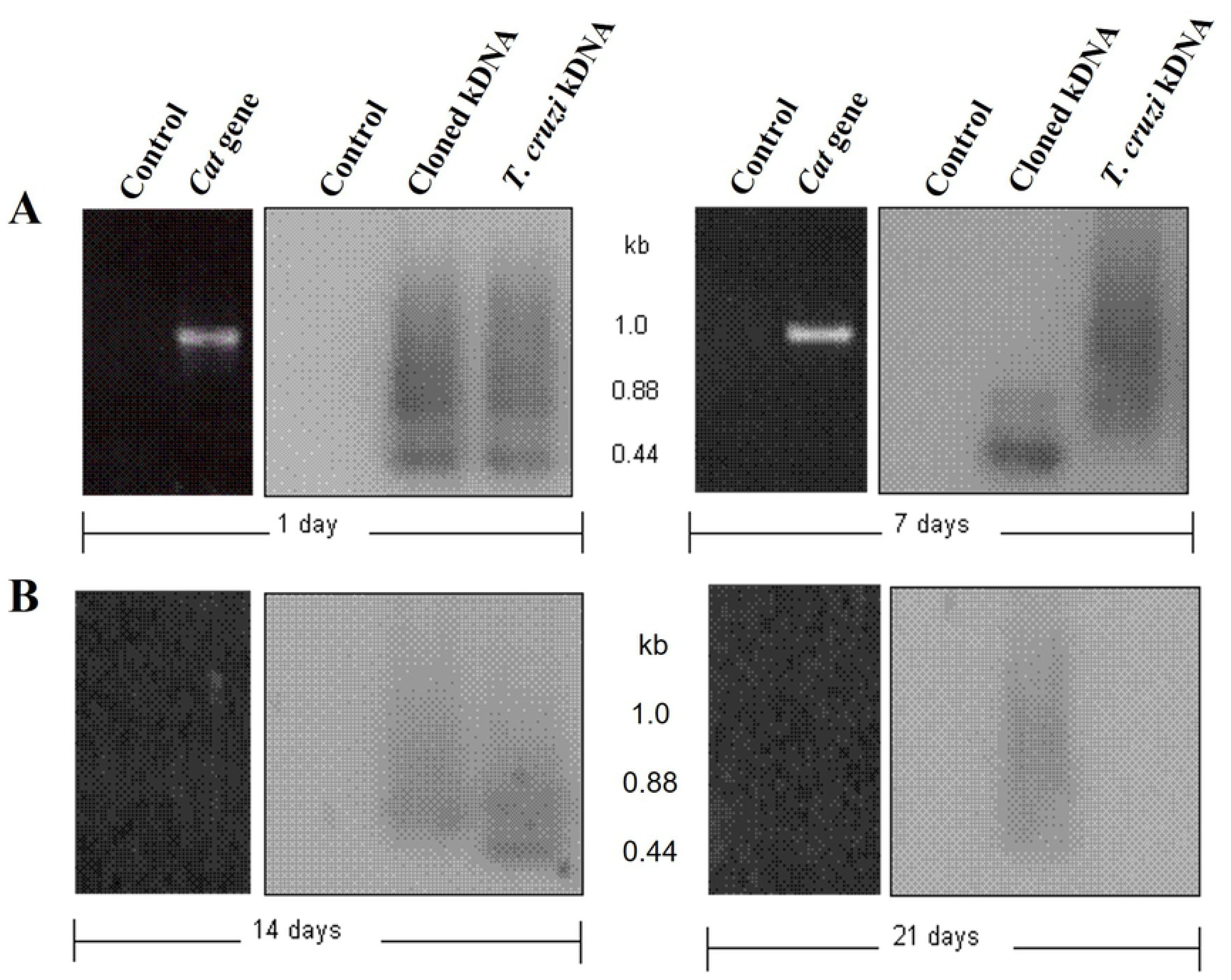
The U937 macrophages waste out the nude kDNA minicircle sequences soon after the ingestion. The DNA sequences uptake by the macrophages in FBS-free DMEM. A) At days 1 and 7 the lipofection-treated *Cat-*gene sequence, the cloned kDNA and the crude kDNA sequences separated in agarose gel 8%, blotted, and probed. B) The *Cat-*DNA cleared out at the day 14^th^, and the cloned kDNA quenched at the day 21^st^.

### The *Trypanosoma cruzi* infection needed to integrate the kDNA minicircle sequences into the host’s cell genome

The results of the above experiments suggested that the integration of the *T. cruzi* minicircle kDNA sequences into the macrophage genome called for the host’s cell-pathogen interactions. Initially, we sought to reproduce the phenomenon *ex-vivo* in the U 937 human macrophage at set points of the infection with the *T. cruzi*-macrophage ratio set at 5 to 1. The PCR amplification of the *T. cruzi-*infected macrophage DNA obtained at the day-7 post-infection showed the 1.2, 1.8, and 2.2 kb bands, in addition to the 330 bp kDNA band. Contrastingly, at the 31^st^ day post-infection the *T. cruzi-*infected macrophage DNA showed the upper 1.8, and 2.2 kb bands, in the absence of the 330 bp kDNA band. This finding documented the kDNA 1.8, and 2.2 kb sequences transfer into the macrophage genome, and the absence of the 330 bp band suggested the low ratio *T. cruzi-*macrophage infection was eradicated. The results of the transfer of the T. *cruzi* kDNA minicircle sequences into the macrophage genome showed in Figure 3. Moreover, the macrophage retained the kDNA upper bands during the monthly serial passages in culture flasks now for over several years [114]. The time-lapse modification of band profiles suggested that the kDNA minicircle sequences integrated into the host cell genome over time could represent a critical element in the host-pathogen interactions. With this respect, we search for drug inhibitors of the phenomenon of the *T. cruzi* kDNA transfer into the host cell’s genome.

**Figure 3.**
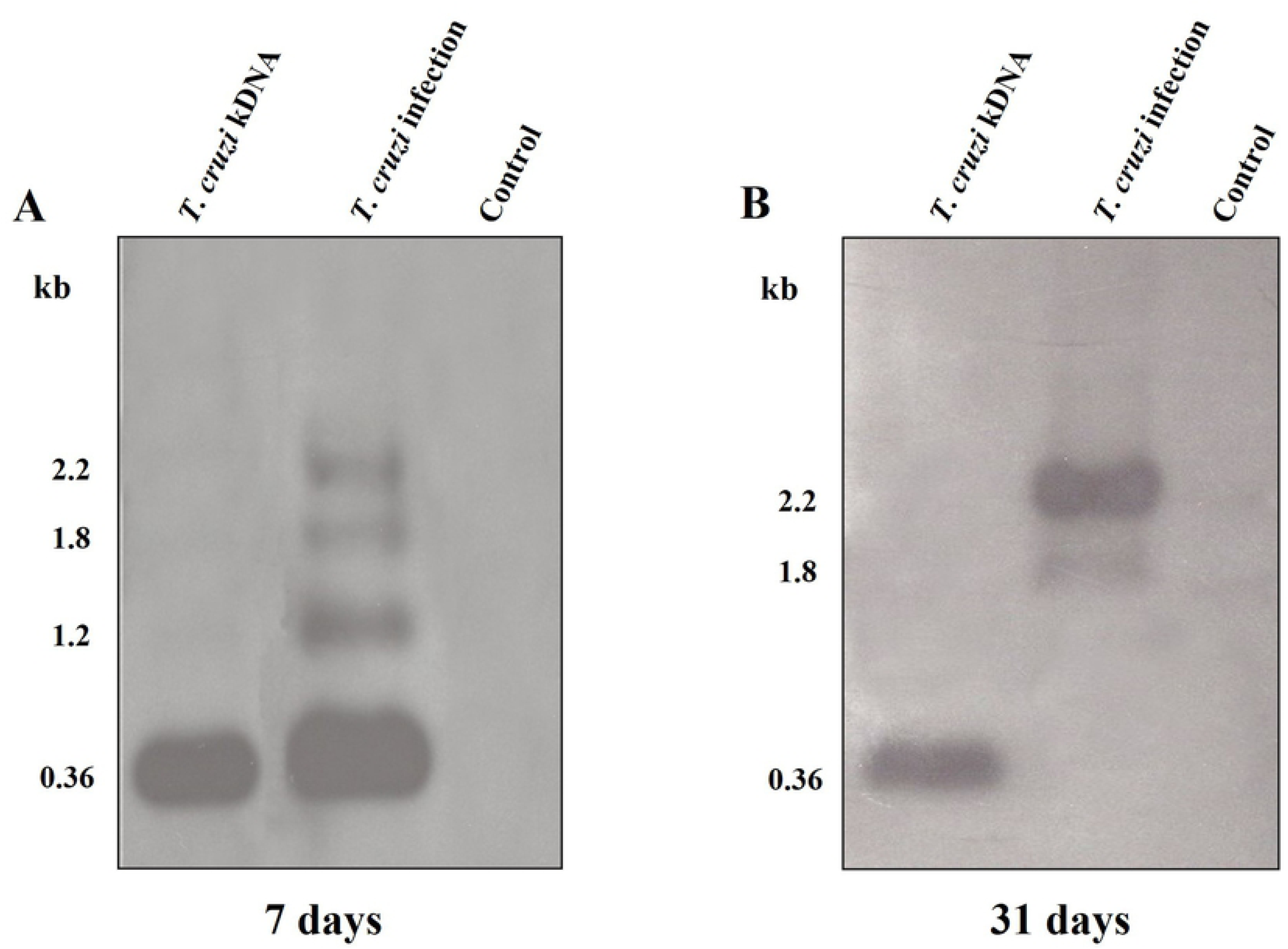
Integration of the *Trypanosoma cruzi* mitochondrion kDNA minicircle sequences into the U937 macrophage genome. The assessment of the macrophage DNA at days 7 and 31 post-infection: - Panel **A**) The PCR amplification of the *T. cruzi* DNA shows the 330 bp kDNA band at lane 1, whereas the *T. cruzi-*infected macrophage DNA of the day-7 post-infection shows 1.2, 1.8, and 2.2 kb kDNA bands at lane 2. The DNA from control, uninfected macrophages (lane 3) showed none. **Panel B**) The 31-day *T. cruzi-*infected macrophage DNA sample retains the top 1.8 and 2.2 kb kDNA bands, only. The control, uninfected macrophage showed none (lane 3).

### Searching for the drug inhibitors of the *T. cruzi* kDNA transfer

The chemical structures and short names of the drug inhibitors of specific metabolic check-point pathways of cell growth and differentiation, showed in the S1 Figure. The select array of inhibitors of eukaryote cells growth aiming at the multidrug treatment of the *T. cruzi* infections with specific objectives: *i*) toxic, lethal effect against the *T. cruzi* forms; *ii*) minimal toxicity to the host cell; *iii*) Overturning the kDNA transfer. The Table 1 depicts the metabolic pathways inhibitors and the optimal concentration determined by the *in vitro* dose response.

**Table 1.** Checkpoint inhibitors of cell growth and differentiation

| Inhibitor | Activity | Posology (µg/100µL)* | References |
| --- | --- | --- | --- |
| **Cell growth arrest** |  |  |  |
| Azidothymidine | Intercalating thymidine to the DNA inhibits reverse transcriptase G2/M phase, caspase-induced cell division arrest, and apoptosis | 37 | [115, 116] |
| Bromodeoxy-uridine | Intercalating thymidine at phase S cell cycle arrests reverse transcription and cell growth | 30 | [117, 118] |
| Colchicine | Suppression tubulin assembly arrests microtubule formation, cell division, and apoptosis | 50 | [119-121] |
| Genistein | ATP inhibition through protein tyrosine kinase (P TK)-and cell growth arrest | 10 | [122, 123] |
| Microcystis-LR | Inhibits PP1 and PP2 at serine/threonine phosphatases and cell arrest | 5 | [124-126] |
| Mitomycin C | Inhibits PP1 and PP2 at serine/threonine phosphatases and cell arrest | 10 | [127, 128] |
| **Polymerase inhibitors** |  |  |  |
| Camptothecin | Inhibits PP1 and PP2 at serine/threonine phosphatases and cell growth inhibition | 50 | [129, 130] |
| Cycloheximide | Inhibits p38 activated protein kinase (MAKPK) and cell entry at G2/M phase | 5 | [131, 132] |
| Etoposide | Down regulate polymerase II at S-G2 phases and prevents cell division | 50 | [133, 134] |
| Ofloxacin | 4-Fluorquinolone inhibits polymerase at G2/M phase, and apoptosis | 30 | [135, 136] |
| Praziquantel | Isoquinolone inhibits serine/threonine kinase (PI3-k) polymerases I and II | 46 | [137-139] |
| Staurosporine | Protein kinases inhibitor (A, C, CAM and PI3K) prevents cell growth at G1 check point | 9 | [140,141] |

\*Each drug suspension ( $\mu\text{g}/100 \mu\text{L}$ ) in 0.15M NaCl used in the cytotoxic *in vitro* assays.

The search for the optimal dose of the cell growth inhibitor secured on the *T. cruzi* epimastigotes (10 x 10^6^/mL) incubated with the concentration of each inhibitor for 4 h at 27 °C (see methods). The cytotoxic effect determined in aliquots of 1 ml of the specific inhibitor-treated *T. cruzi* forms suspension in 5 mL of fresh LIT medium and growth in a shaker incubator at 27 °C for 14 days. The motile epimastigotes showing typical whitish silhouette in a counting chamber was recorded and plotted, and the cytotoxic effect of each inhibitor to show in the Figure 4. The findings revealed that Bromodeoxyuridine, Azidothymidine, Praziquantel, and Ofloxacin, respectively, killed 96%, 94%, 92%, and 85% of the *T. cruzi* epimastigotes. The parasite killing was confirmed in the trypan-blue dye staining of the immobile flagellate. The inhibitor concentration showing lethal effect to the *T. cruzi* epimastigotes and undetectable toxicity to the muscle cells was employed in the *ex vivo* investigation.

**Figure 4.**
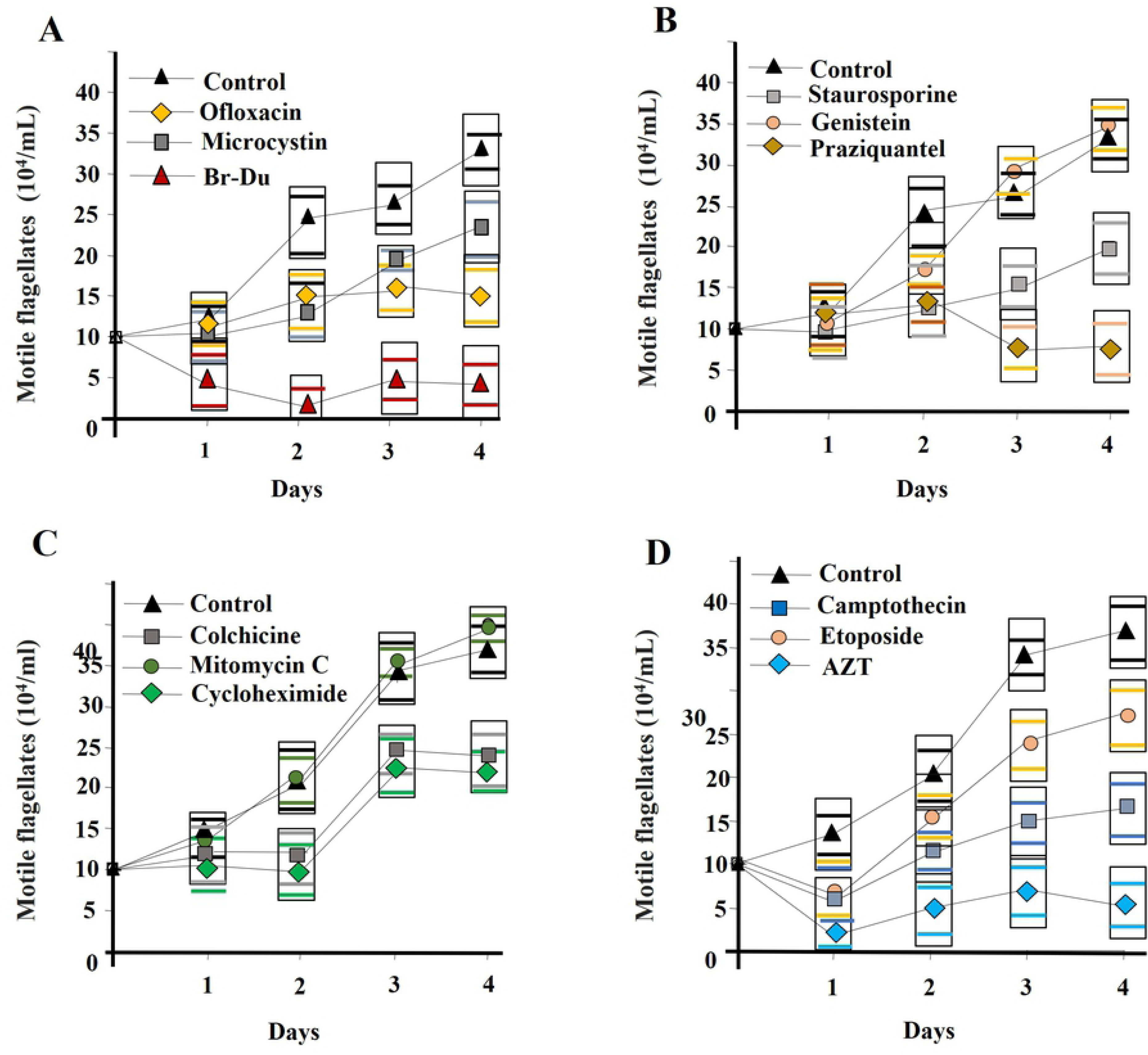
The effect of specific inhibitors of cell growth and differentiation on the *Trypanosoma cruzi* epimastigote forms. Growth curves depicted the white, motile parasites. Note that Ofloxacin (A), Praziquantel (B), and AZT (D) showed high lethal effect upon the flagellates. The remaining of the drugs showed much less pathogen killing activity (A to D).

The experiments further searched for the inhibitor toxic effect against the L6 muscle cells’ host of the *T. cruzi* infection. The suspension of 10 x 10^5^*/*mL muscle cells in DMEM was incubated with the inhibitor optimal final concentration shown in Table 1. The mock control *T. cruzi-*infected muscles cells suspension maintained in the absence of the drug inhibitors. After 4 hours of incubation at 37 °C, 3 mL of the cells’ aliquots seeded in tissue-Tek (Fischer Scientific) culture plate with complete DMEM. After two-week incubation at 65% in a humidity chamber into a CO2 incubator at 37 °C, the muscle cells monolayers washed with PBS, H-E staining, and exams. The counts of the *T. cruzi* amastigotes and of the host cells under the microscope yielded the information showed in the Figure 5. The toxic effect of the inhibitors on the host cells revealed the blurred fade-away silhouette of some amastigotes, whereas the muscle cells showed the loss of the membrane boundaries and vacuolation of the cytoplasm. None of these features were seen in the eukaryotic cells in the absence of the drug inhibitors. The graphic representation of the toxic effect of the drug inhibitors on the muscle cells showed that Microcystin, Bromodeoxyuridine, and Mitomycin inhibited the growth of both the *T. cruzi* and the L6 muscle cells. Ofloxacin and Azidothymidine did not show the host cell detectable toxic effect at the optimal concentration chosen for the *in vivo* treatment of the *T. cruzi-*infected L6 muscle cells.

**Figure 5.**
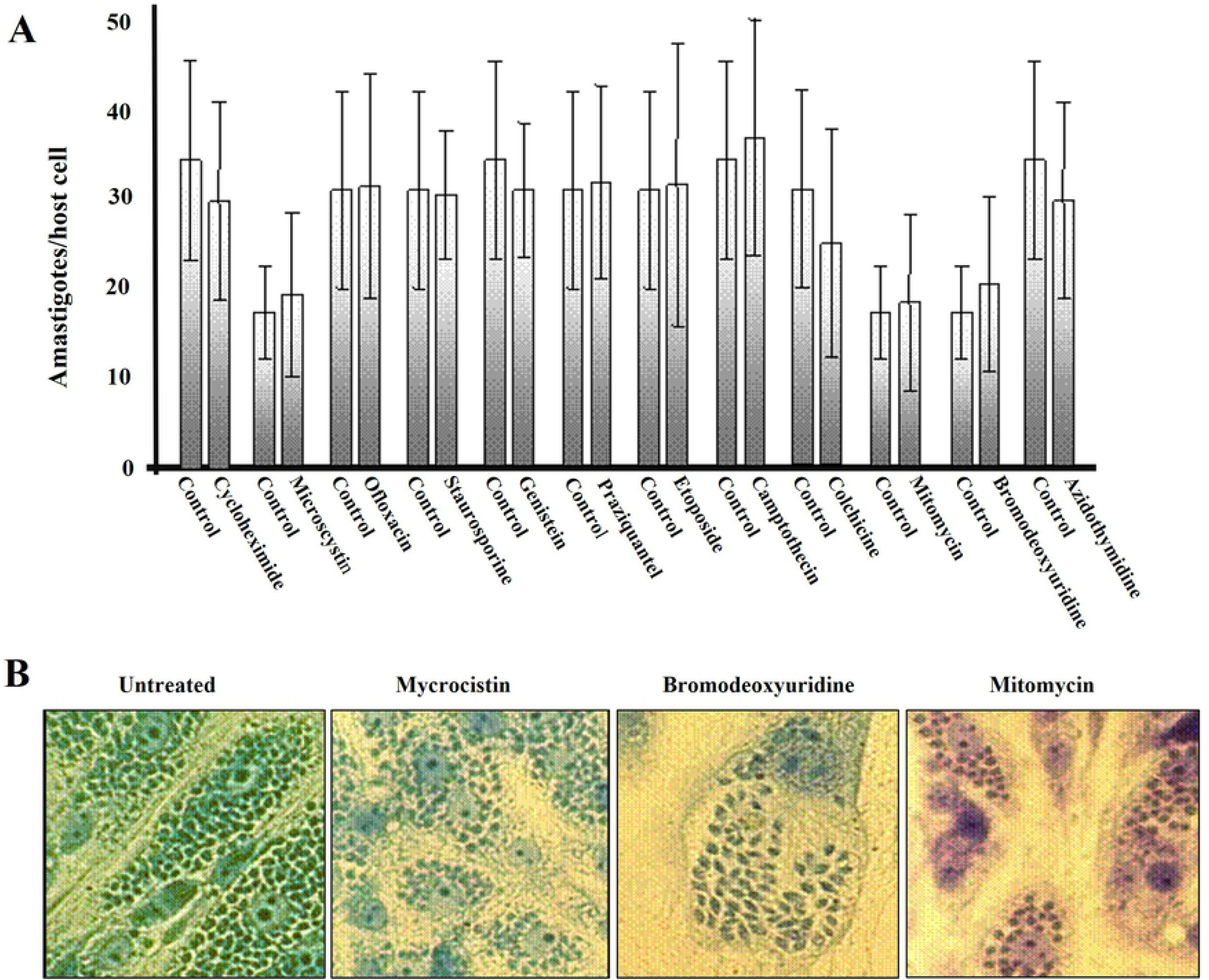
The cytotoxic effect of drug inhibitors of metabolic pathways on the growth of *Trypanosoma cruzi* and of the L6 murine muscle cell. The cytopathic effect recorded at two-week growing cells, staining and microscopy. **A)** Histogram showing the growth of *T. cruzi--*amastigotes into the L6 muscle cells. Note that Microcystin, Mitomycin, and Bromodeoxyuridine showed significant inhibition (p < 0.05) of the host cell and of the parasite growth. **B)** Microscopic detection of the toxic effect of the Microcystin, Mitomycin and Bromodeoxyuridine on the L6 muscle cells. 1) Normal growth of the untreated host muscle cells-swarm the dividing *T. cruzi* amastigotes, in the absence of a drug inhibitor. 2) Severe cytopathic effect of Microcystin-LR translates into loss of the muscle cell boundary, foamy cytoplasm, and amastigote lysis. 3) The intense toxicity of the Bromodeoxyuridine. 4) The Mitomycin-C moderate toxic foamy effect and loss of the membrane boundary.

Having shown the dose of each inhibitor that did not kill the murine muscle cell we searched for the effect of the drug inhibition of the kDNA transfer to the genome of the *T. cruzi-*infected human macrophage. The Southern blot analysis of the *T. cruzi*-infected macrophage DNA-PCR amplification products obtained with specific primer sets and radio label kCR probe showed that the phosphorylases, polymerase II, and late phase cell division inhibitors: - Camptothecin, Cycloheximide, Etoposide, Genistein, Ofloxacin, Praziquantel, and Staurosporine -, prevented the high size 1.2, 1.8, and 2.2 kDNA minicircle sequence bands. In addition, Colchicine that suppress tubulin assembly and microtubule formation, and Azidothymidine intercalating thymidine to DNA arrest reverse transcriptase and telomerase at G2/M cell growth phase, overturned the transfer of those high size 1.2, 1.8 and 2.2 kb kDNA sequence bands. Contrarily, the Microcystin, Camptothecin, and Bromodeoxyuridine, respectively, phosphatases, polymerase I, and forming DNA adduct at phase S cell cycle arrest reverse transcription and cell growth, did not overturn the transfer of the kDNA 1.2, 1.8 and 2.2 kb bands into de host cell. Interestingly, the Microcystis, a serine/threonine phosphatase PP1 and PP2A, overturned the 0.330 kb kDNA band, only (Figure 6). The Table 1 showed the inhibitors’ concentration employed to treat the U937 macrophages at the *T. cruzi-*host cell ratio set at 5 to 1.

**Figure 6.**
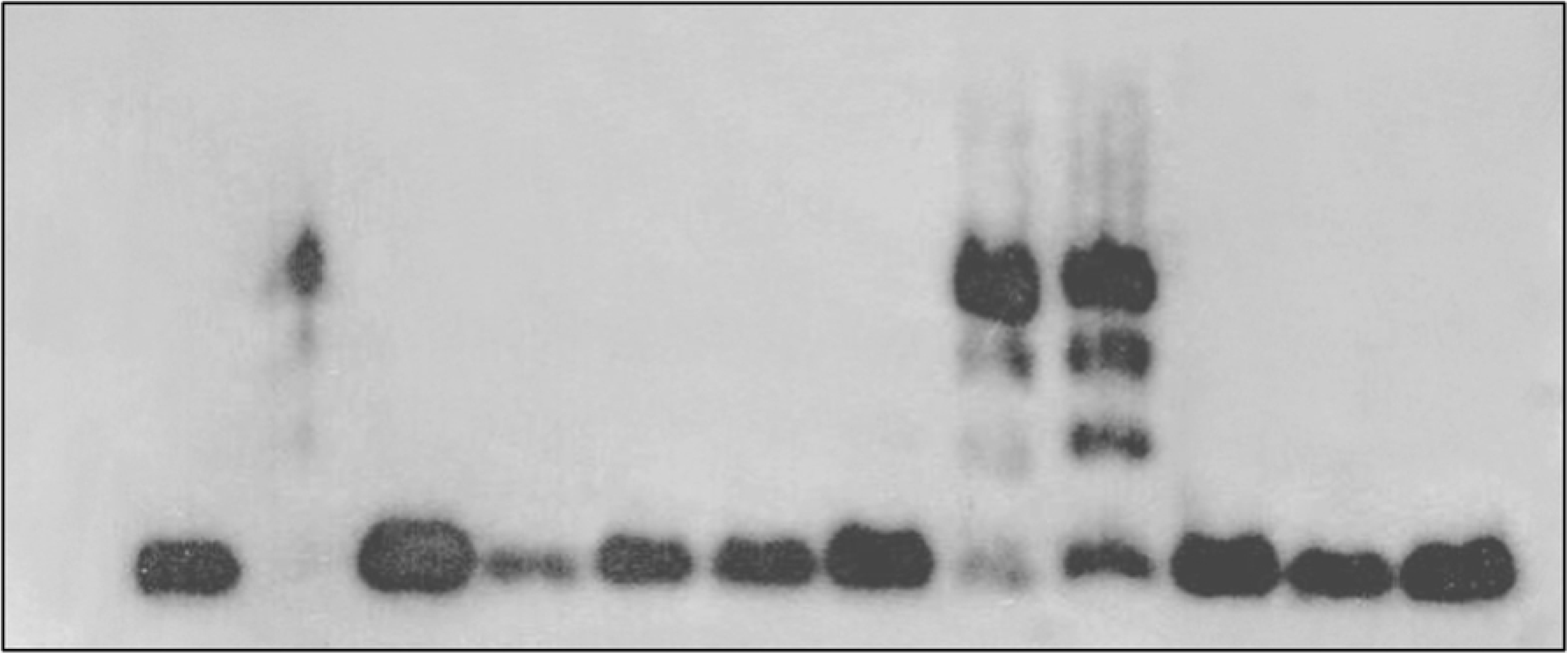
Inhibition of lateral transfer of the *Trypanosoma cruzi* kDNA minicircle sequence to the macrophage genome. The macrophages were collected at day 21 post-infection with the *T. cruzi* and treated with the inhibitor concentration showed in the Table 1. The DNA-PCR amplification products that hybridize with the radio labeled kCR minicircle probe revealed that phosphorylases, polymerases, and late phase cell division inhibitors precluded the transfer of the 1.2, 1.8 and 2.2 kDNA bands into the macrophage genome. The Microcystin, Camptothecin and Bromodeoxyuridine inhibitors did not prevent the transfer of the high molecular weight bands. The Microcystis prevented the 330 bp kDNA band. The positive control (*T. cruzi* DNA) revealed the 330 bp kDNA minicircle band, and the negative control (untreated macrophage DNA) showed none.

## Animal model systems

To confirm and dissect the effect of the drug inhibitors of the eukaryote cell growth, we moved our studies into rabbits and mice, which are amenable to *T. cruzi* infection, and chickens, which are refractory to persistent infection [35, 41, 57, 143–149] to show the kDNA minicircle sequences transfer sites into the vertebrate genome, aiming at the multidrug treatment of Chagas disease. The invasion of the blastula embryo by *T. cruzi* shown by tidy demonstration, and the possibility of *T. cruzi* uptake by stem cells *ex vivo* we first explored. The rabbit embryo stem cells were readily invaded by *T. cruzi* trypomastigotes. Rabbit zygote blast cells actively engulfed the protozoa, with penetration of the host plasma membrane 6 h after adhering to the plastic surface. After the fourth day in culture, dividing amastigotes were identified in the growing embryo stem cells (Figure 7A: a, b, c and d).

**Figure 7.**
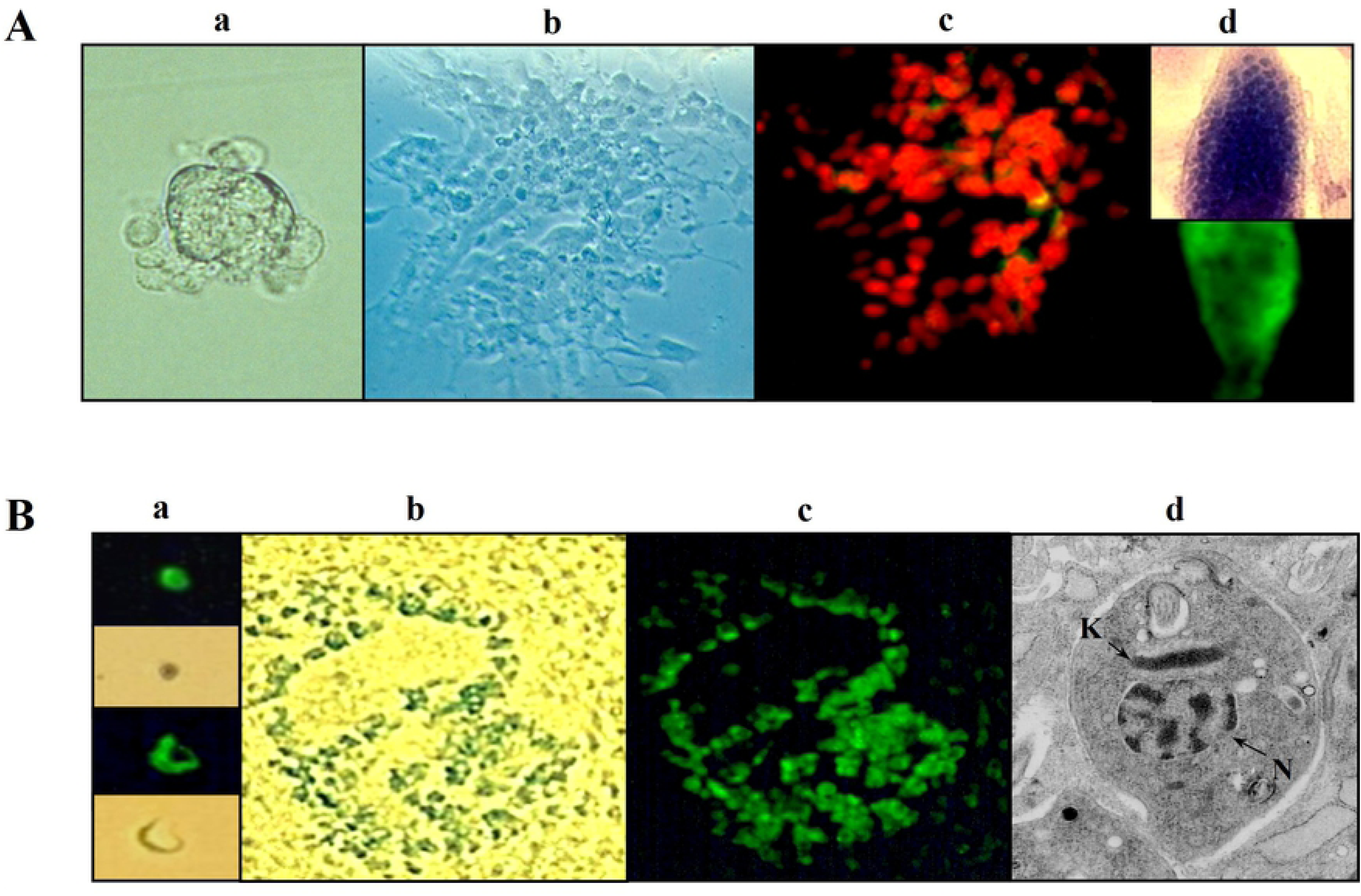
Growth of *T. cruzi* in embryonic stem cells of mammals and birds. A) A rabbit zygote at stage 2 days postcoital blasted off stem cells that adhered to the surface and was exposed to 1,000 *T. cruzi* trypomastigotes (a). A layer of stem cells formed on the fourth day in culture. The phase contrast microscopy showed intracellular *T. cruzi* amastigotes (b). Magnification 200x. Same field, in which nuclei of stem cells counterstained with ethidium bromide, revealed clump of immunofluorescence-staining intracellular amastigotes (c). Embryonic stem cell showed amastigotes counterstaining with Hematoxylin-Eosin; the same cell below showed immunofluorescent intracellular amastigotes of *T. cruzi* recognized by a specific antibody (d). Magnification 800x. B) The *T. cruzi* amastigote and trypomastigote forms showed by the specific immunofluorescent antibody, and brown forms of the same unstained *T. cruzi* stages detected by phase contrast microscopy (a). Magnification 1,200x. Paraffin embedded section of a chicken showing blue X-Gal-stained cells in the endoderm and mesoderm tissues of a 4-day-old embryo (b). Magnification 200x. The immunofluorescence staining colocalizes the *T. cruzi*-infected cells on the same embryo tissue section (c). Electron micrograph showing an intracellular amastigote (d): n, nucleus; k, kinetoplast.

The kinetics of the *T. cruzi* infections observed, also, in the fertile chicken eggs inoculated with 100 *T. cruz*i trypomastigotes prior to the incubation at 37 ° C. On the eight days of incubation, the embryo stem cells with the dividing amastigotes were identified by immunofluorescence and X-Gal staining. Using this immunohistochemical approach, we confirmed amastigotes of *T. cruzi* in chicken embryo endoderm and mesoderm tissues (Figure 7B: a, b, c, and d) at stages corresponding to 4 and 8 days old. The dividing *T. cruzi* amastigote in the embryo stem cell that was examined under the electron microscope showed typical disc-shaped mitochondrion kDNA network and membrane wrapped round nucleus containing DNA (Figure 7B, d).

The permissiveness of the embryonic stem cells to a *T. cruzi* infection at the blastula stage (2-day-old zygote) was an indication that differentiating germline cells in the genital crest, which appear at days 4-8.5 of gestation [149–151] could contain kDNA minicircles due to invasion. Thus, these cells were surrogates of the parasite-host cell interactions, such as early markers of the effect of the inhibitors on the cell growth and prevention of the kDNA transfer. The de novo transfer of kDNA to the genome of mammals and birds supports the contention that genetic transfer may be part and parcel of the normal course of the *T. cruzi* infections. To emphasize this point, we took advantage of our ability to infect the rabbit and the chicken embryo with *T. cruzi*.

### The rabbit models

#### The *Trypanosoma cruzi* infection of rabbits and treatment with the nitro-heterocycle Benznidazole

Initially we sought to reproduce the phenomenon in the rabbit model infection system [142]. The *T. cruzi-*infected rabbits died of the chronic Chagas heart disease indistinguishable to that described in humans [145–149]. The active process of genetic transfer between *T. cruzi* and rabbit was initiated with the examination of effect of the nitro-heterocycle Benznidazole lead in the treatment of the human Chagas disease [143-148, 156-158]. We run experiments in groups of *T. cruzi* infected rabbits to finding out whether the administration of the nitro-heterocycle that curtailed the parasitemia (Figure 8) could halt the transfer of the kDNA and the Chagas heart pathology. Two groups of eight two-month-old male and female rabbits in individual cages each inoculated with 2.5 x 10^6^ *T. cruzi* trypomastigotes subcutaneously. Forty days after the *T. cruzi* infections, the group A rabbits received 8 mg/kg/day of the Benznidazole intraperitoneal injection for 60 days. The therapeutic regime consisted of the drug powder suspension in 0.15 M saline, which was administered from the 40^th^ to the 100^th^ day post-infection. The levels of parasitemia detected by xenodiagnoses, in which each of 20 first instar nymph of the Reduviid bug *Dipetalogaster maximus*, obtained the blood fills from each *T. cruzi*-infected rabbit of groups A and B. The xenodiagnoses showed the *T. cruzi-*infected, Benznidazole treated group A rabbits did not show the parasite in the blood, as indicated by absence of the flagellate in the bugs’ excreta, during the 40^th^ day of the drug regime and thereafter. Contrastingly, the infected-untreated rabbits (group B) yielded persistently positive xenodiagnoses for 150 days. The *T. cruzi-*infected rabbits (groups A and B) showed the ELISA specific antibody titers above 1:80 serum dilutions throughout the long-lasting *T. cruzi* infections regardless of the drug interference. Despite of the drug treatment the serologic assays showed the specific antibody titers through the long-lasting course of the *T. cruzi* infections (asterisks, Figure 8).

**Figure 8.**
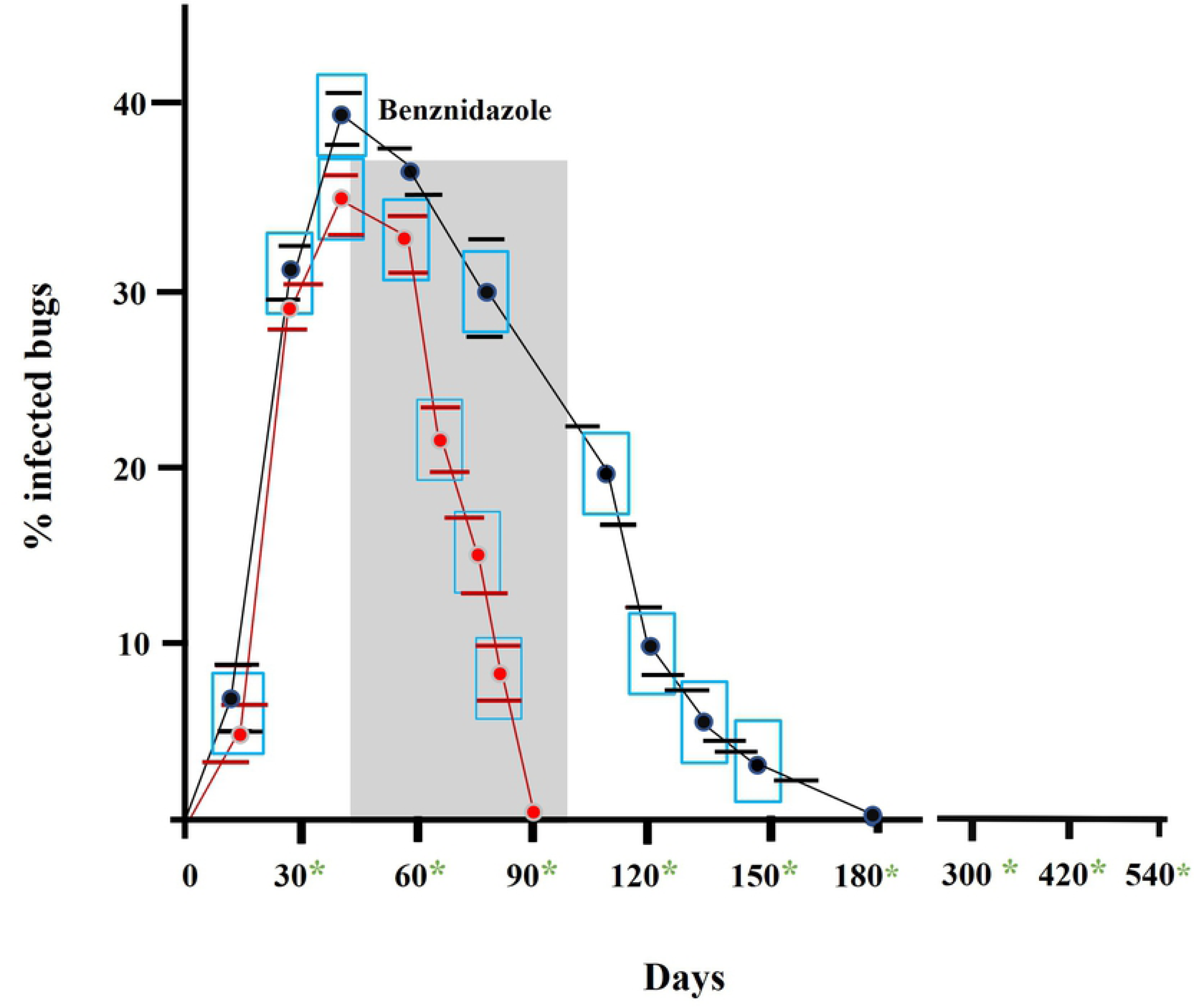
The Benznidazole effect on the levels of parasitemia in the *Trypanosoma cruzi-*infected rabbits. Two-month-old rabbits (groups A and B, n = 8) infected subcutaneously with 2.5 x 10^6^ flagellates. Forty days after the infection the group A rabbits treated with the Benznidazole, 8 mg/kg/daily for sixty days. The parasitemia of the groups of rabbits A and B determined by the number of clean first instar nymphs of *Dipetalogaster maximus* bugs fed on the Chagas rabbits. The percent parasitemia achieved by the total number of *D. maximus* that became infected with *T. cruzi /*divided by the number of bugs showing the *T. cruzi* in the excreta examined 30 and 60 days after a blood meal. Shaded area, period of the drug administration. Bars = standard deviation. Note the parasitemia curtailed at the 90^th^ day post infection of the Benznidazole treated rabbits, whereas the untreated *T. cruzi*-infected rabbits sustained the parasitemia until the 150^th^ day post infection.

#### Integration of *Trypanosoma cruzi* kDNA minicircle sequences into the rabbit genome

The kDNA integration sites associated with modification in the human genome could represent a critical biological element in host-pathogen interaction leading to the chronic clinical manifestations of Chagas disease. Initially we thought to reproduce the phenomenon in the rabbit model infection system. The active process of genetic transfer between *T. cruzi* and rabbit was initiated with the examination of DNA from tissues extracted from eight animals that had been infected and treated with Benznidazole for six months to three years. Digests of DNA extracted from heart, skeletal muscle, liver, intestine, and kidney were hybridized with the kCR constant region probe or the *T. cruzi* minicircle. A 2.2 kb band was obtained in the heart and large intestine derived DNAs, distinct from the minicircle unit-sized 1.4 kb band hybridizing in parasite DNA alone. DNA extracted from tissues of an uninfected rabbit showed no hybridization with the kCR probe. The tissue samples showed no hybridization with the other *T. cruzi*-specific nuclear DNA probes for high-copy number genes or repetitive sequences [153–155, 159]. To characterize the 2.2 kb band, 5’ race was employed on DNA from infected rabbit heart. This approach generated sequence containing both arms of rabbit DNA flanking the kDNA insertion (Figure 9). Integration of a CCA/ACC rich kDNA fragment (bp 1255-1907) occurred at attachment sites of direct CACCAACC repeats within the rabbit DNA. The extreme 3’ region (1908-2064 bp) has homology with the LBNL1-125D4 clone (GenBank accession number AC144399) a rabbit LINE containing interspersed SINE repeats. An ORF initiating in the host DNA and extending beyond the kDNA integration site (1217-1582 bp) translates a putative chimeric r45-like antigen (GenBank accession number AAR24603.1)

**Figure 9.**
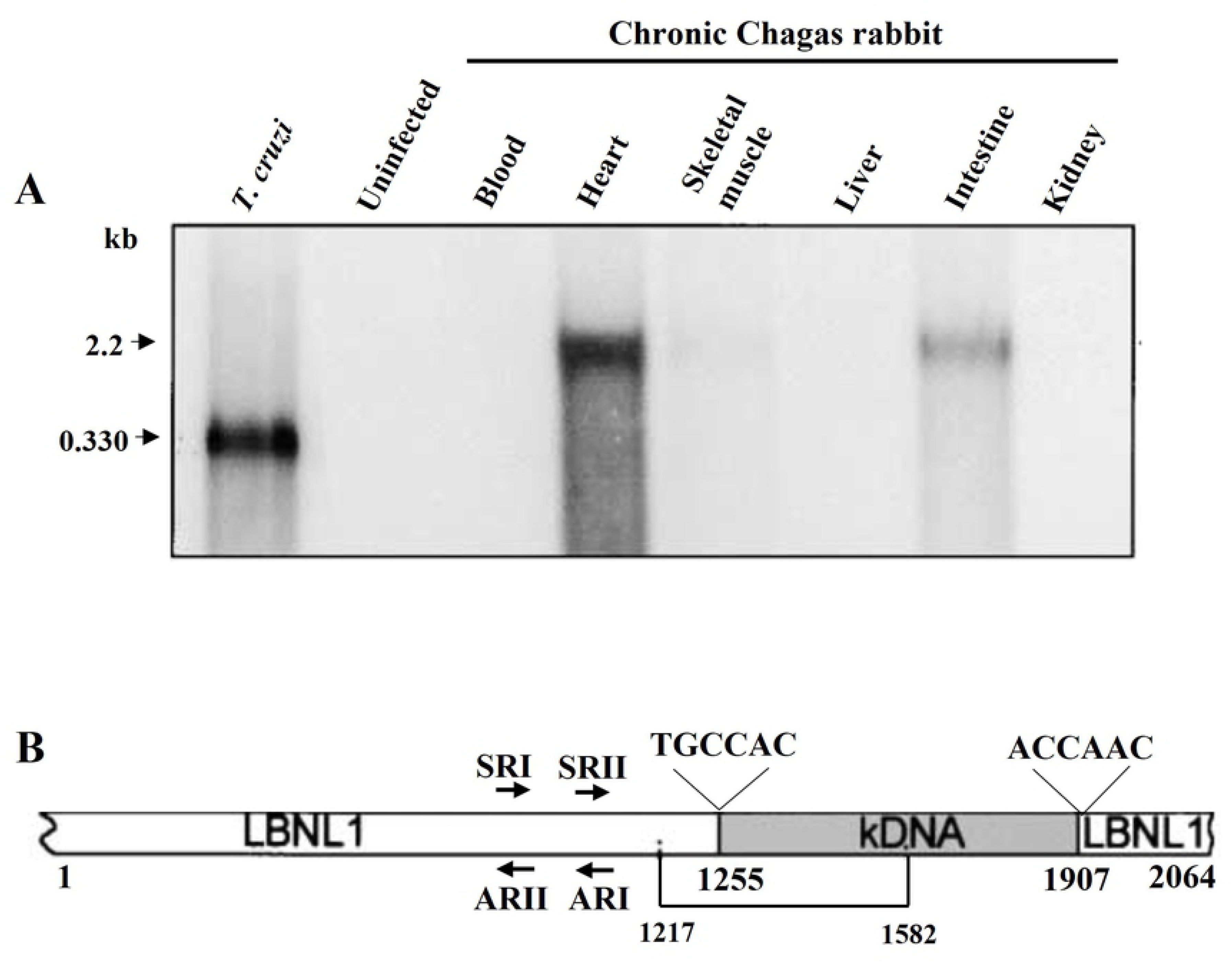
Integration of kDNA minicircle sequence into the genome of a Chagas disease rabbit. A) Hybridization of rabbit DNA with a specific kDNA probe. *Eco*R1 digested DNA (20 µg) separated in 1% agarose gel was used for Southern hybridization with 1 µg of *T. cruzi* kCR probe. B) Schematic representation of kDNA integration into rabbit DNA. Arrows show primers used in 5’ RACE to detect kDNA insertion into the rabbit genomic clone LBNL1. Integration of CCA/ACC-rich kDNA (bp 1255-1907) occurred within rabbit DNA showing attachment sites of direct short CCAACC repeats. An ORF spans the chimeric sequence bp 1217-1582.

To continue the investigation on the host-pathogen interactions in the rabbit, four sexually mature does and two bucks were crossbred during chronic infection. After three pregnancies, the does with chronic *T. cruzi* infections delivered 104 litters (26 ± 6 per doe), among which 23 (22.1%) were stillborn. Three controls, uninfected does in three pregnancies delivered 96 litter (32 ± 3 per doe), among which 22 (22.9%) were stillborn. PCR was carried out on DNA either from specific tissues of still born animals or from blood cells of individual surviving offspring of the *T. cruzi*-infected rabbits for both *T. cruzi* nDNA and kDNA. A sample showing the presence of kDNA but not nDNA (Figure 10A, offspring 1 of doe C) was examined by Southern hybridization of *Eco*RI-digested genomic DNA with a kDNA probe (Figure 10B). A band that was larger in size than the unit minicircle was detected, indicative of an integration event. The control uninfected rabbit DNA showed the absence of bands that hybridized with nDNA or kDNA probes. Out of the 104-surviving offspring from chronically infected parents, 15 (14.4%) contained nDNA, and 24 (23%) contained kDNA by the PCR assay. Nine stillborn yielded DNA from heart, skeletal muscle, liver, spleen, and large and small intestine, and each tissue type formed specific bands by hybridizing amplification products with kCR probe. Genomic DNAs of six kDNA-positive offspring were subjected to 5’RACE, yielding six integration sites of kDNA fragments (S3 Table). Three of these clones showed the CCAACA motif flanking integration sites and potential ORFs for chimeric proteins [159].

**Figure 10.**
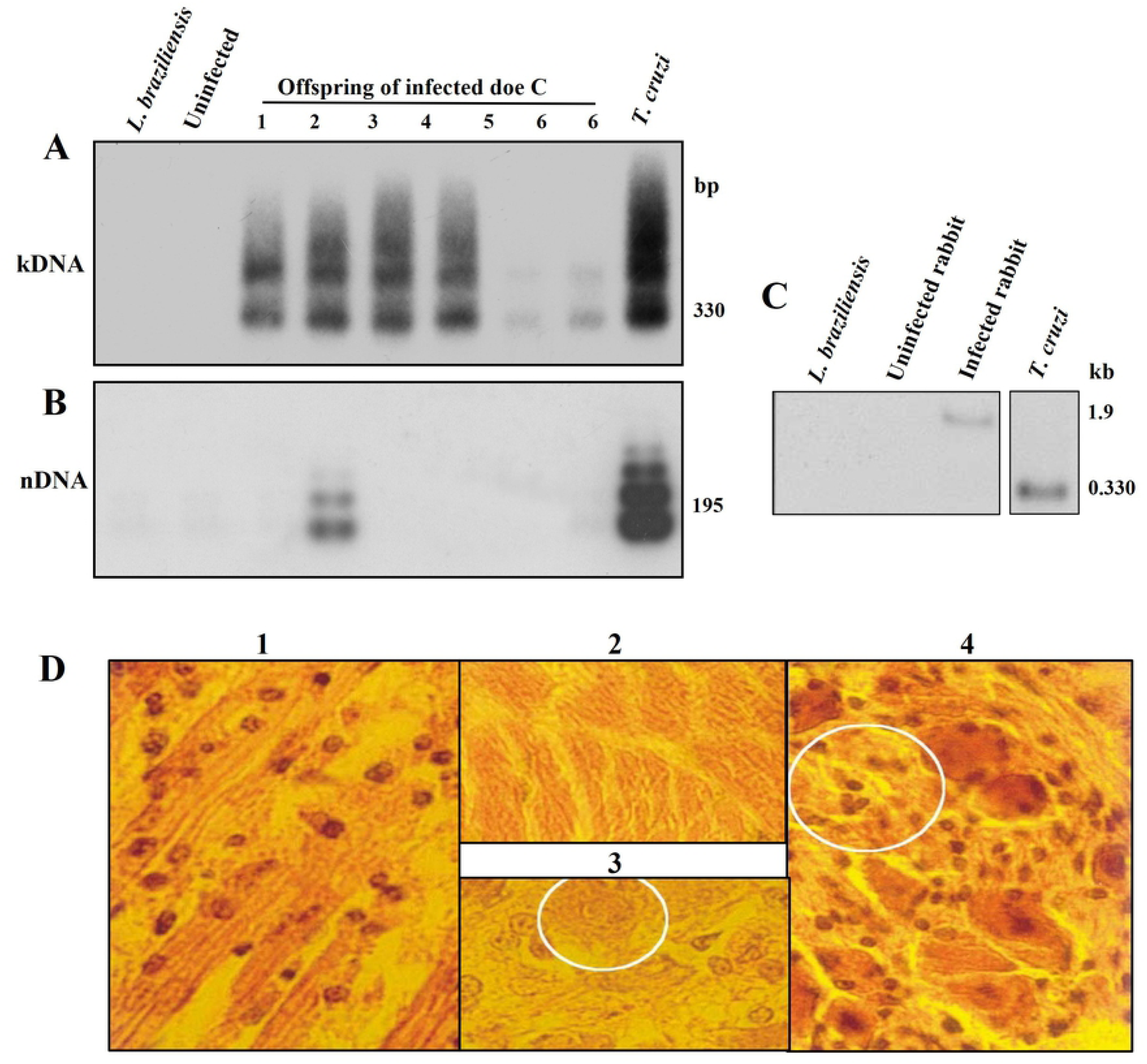
Genetic markers of *Trypanosoma cruzi* infection in offspring of Chagas disease rabbits, kDNA transfer, integration, and pathology. Hybridization of PCR amplification products from template DNA obtained from offspring of Chagas disease doe using sets of *T. cruzi* nDNA and kDNA primers, and specific radio labeled nCR and kCR probe. A) Evidence of genetic markers of the *T. cruzi* kDNA transfer. Specific hybridization of PCR amplification products from template DNA obtained from offspring of Chagas disease doe C using sets of *T. cruzi* kDNA specific primers. The *Eco*RI digested DNA products resolved in 1% agarose gels. PCR analysis of kDNA amplification showed bands of 330 bp and catamer from the parasite DNA and from genomic DNA of six progeny with hybridization with the kCR probe. This event showed the sexual transmission of the *T. cruzi* infection from Chagas rabbits’ parental to the offspring 2 of Doe C. B) PCR analysis of nDNA amplification showed band of 195 bp and catamer formed with parasite DNA and with genomic DNA from offspring 2 after hybridization with the specific internal probe. This event indicated the transfer of the *T. cruzi* nuclear DNA from the parental to progeny. C) Integration of kDNA minicircles into the genome of offspring C1 from a *T. cruzi-* infected doe. Test and control DNA (20 µg) was digested with *Eco*RI along with 10 ng of *T. cruzi* or *L. brasiliensis* DNA for Southern hybridization against the kCR probe. Separation of DNA products achieved as described in Figure 1. The presence of the 1.9 kb band size higher than the minicircle 330 bp band indicated the integration of the kDNA into the rabbit genome. D) Destructive myocarditis and lysis of target heart cells. Note the round lymphocytes adhered to the surface of the target cells. 1) Histopathological section showing mononuclear cell infiltration and lysis of target heart cells, and lymphocytes adhered to the surface of the target cells. 2 and 3) Normal histological features of myocardium and intracardiac ganglion cells from a control offspring of a noninfected rabbit. 4) Intracardiac parasympathetic ganglion showing mononuclear cell infiltration and neuron drop out (circle).

Moreover, the sections from the tissues of the offspring 1 from doe C showed destructive myocarditis and ganglionitis. Tissue specific histopathological lesions in muscle tissues, which were usually extensive in the peripheral nervous systems of stillborn offspring of *T. cruzi*-infected rabbits, were seen. Myocarditis and ganglionitis, which are typical of Chagas disease in humans were documented in offspring 1 of doe C, consisting of mononuclear cell infiltration and lyses of parasite-free target host cells (Figure 10D) like that seen in the adult *T. cruzi-*infected rabbit. None of these lesions was present in the spleen, in the liver, or in the kidney of offspring from Chagas disease rabbits or in any tissue of control offspring from uninfected rabbits.

This set of experiments demonstrated the high frequency of lateral kDNA transfer and integration *in vivo* into the vertebrate host genome. The resulting offspring could harbor persistent infection, but the presence of kDNA fragments in diverse tissue types suggested that some integrations occurred throughout the germ line of the host. In addition, the results showed that the *T. cruzi* infections in the rabbit as well as in humans can be vertically transmitted from males and females to mates by intercourse [33–36].

The *de novo* transfer of kDNA to the rabbit genome supports the contention that the genetic transfer phenomenon is identical to its analogue recognized in humans [6]. The pathologic findings fostered the search for exploring the Chagas disease manifestation in the benznidazole treated adult rabbit.

#### Clinical and Pathological findings in Chagas rabbit

The X-Rays anticipated the dilated cardiomegaly gross pathology that evolved to clinical heart failure [145, 146]. The heart of a *Trypanosoma cruzi-*infected and Benznidazole treated rabbit with dilation of the atria and the ventricles showed in Figure 11. The histopathology revealed features of the autoimmune heart pathology, whereby the cytotoxic lymphocyte lysis the target cell. The severe myocarditis in the Benznidazole treated (group A) rabbits was neither quantitatively nor qualitatively different from that seen in the untreated *T. cruzi-*infected, group B rabbits. Tissue-specific histopathological lesions in muscle tissues, which usually extended to the peripheral nervous systems of stillborn offspring of *T. cruzi*-infected rabbits, were seen. Myocarditis and ganglionitis, typical of Chagas disease in humans were documented in rabbits [146]. The lesion in the heart consisted of mononuclear cells infiltration and lysis of parasite-free target host cells. None of these lesions were seen in any tissue of control, uninfected rabbits.

**Figure 11.**
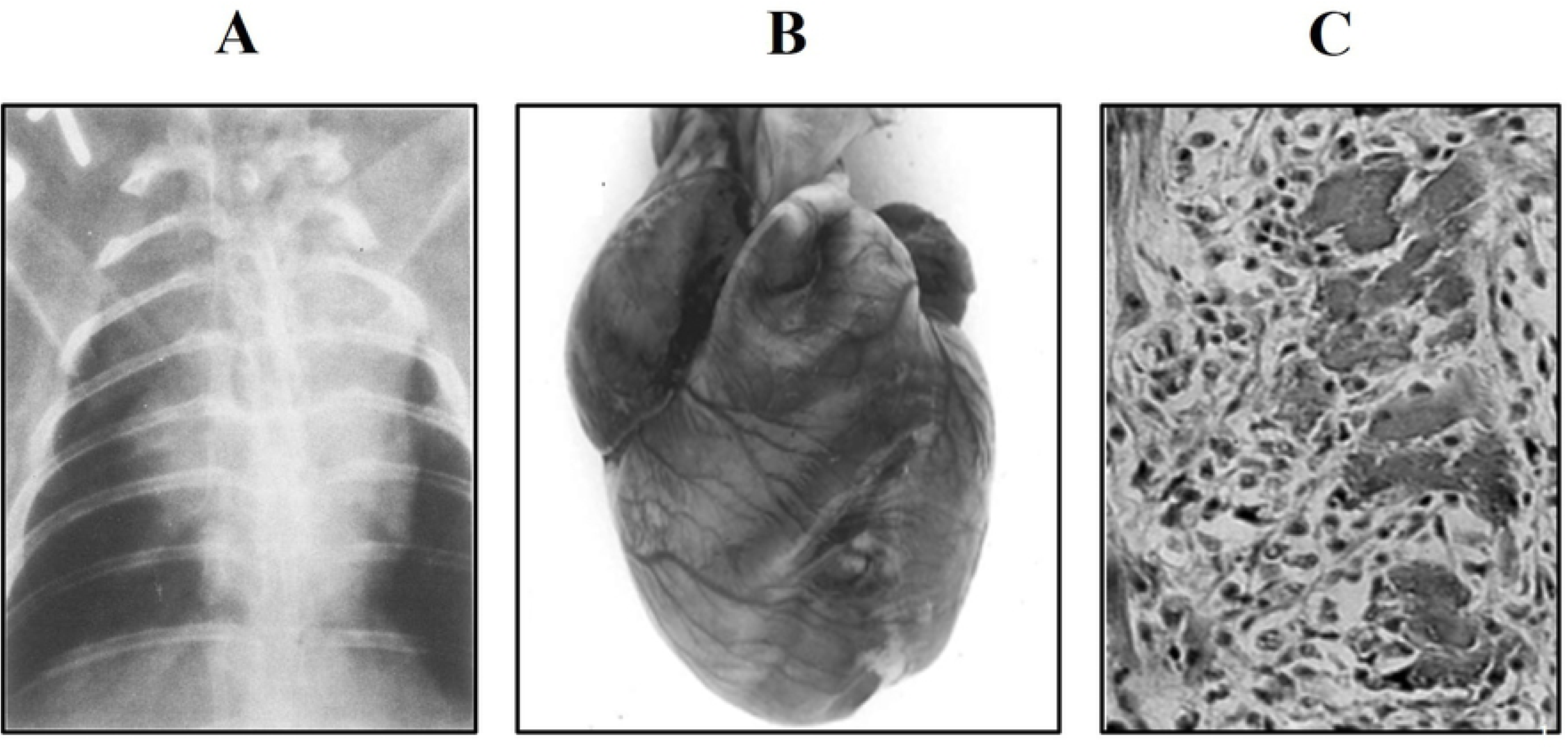
The pathology in the *Trypanosoma cruzi-*infected rabbits treated with Benznidazole. A) Chest X-Rays silhouette showing the dilated heart. B) Cardiomegaly with dilation of the right atrium and ventricle, and of the left ventricle. C) The histopathology with infiltration of cytotoxic lymphocytes that lysis the target heart cells.

This set of experiments demonstrated the vertical transmission of the *T. cruzi* infection in one offspring of the Doe C, whereas the lateral kDNA transfer was documented in the genome of six progeny of the Chagas’ rabbit. Although the number 2 offspring harbored the persistent infection, the progeny showing the kDNA fragments in diverse tissue types suggested that some integrations occurred shortly after parasite invasion thus, resulting in the transfer of the kDNA throughout the germline of the host. To clearly show the postulate we carried out the experiments to search for the lateral transfer of kDNA in the chicken refractory to the *T. cruzi* infection.

### The chicken model system

#### Heritable integration of kDNA minicircle sequences from *Trypanosoma cruzi* into the chicken genome

The permissiveness of the Aves embryonic stem cells to *T. cruzi* infection at the blastula stage (2-day-old zygote) indicated that differentiating germline cells in the genital crest, which appear at days 4—8.5 of gestation [145–152] could contain kDNA minicircles due to invasion. Thus, these cells considered candidates for lateral or germline transfer of parasite DNA.

The experiments in the chicken model system explored to dissociate the kDNA integration from cryptic *T. cruzi* infection. We demonstrated that the chickens are susceptible to *T. cruzi* only at the early embryonic stage, after which they are refractory to persistent infection [111]. Therefore, a kDNA integration event occurring early in the embryonic developmental process could result in the generation of a sexually mature chicken with kDNA integrated into gonadal tissue.

Thirty-six fertile chicken eggs were each injected with 100 *T. cruzi* trypomastigotes. The embryo tissue collected on the second, fourth, end eight days post infection yielded nDNA and kDNA amplification products; however, tissue collected on the tenth, 12^th^, 18^th^ and 20^th^ days of incubation yielded amplification products only for kDNA, indicative of the clearance of the early active infection (Figure 12). Two roosters (4938 and 4973) and two hens (4948 and 4979) that hatched from *T. cruzi-*infected eggs, then showed positive hybridization bands with the kCR in a Southern blot performed on DNA isolated from sperm and from nonfertilized eggs, with their pattern of migration differing from that of free-relaxed or of free-catenated minicircle DNA. These birds were raised for crossbreeding. In the control group, 14 fertile chicken eggs were subjected to PCR, and neither nDNA nor kDNA was detected. Additionally, we inoculated naked minicircle or cloned minicircle sequences in the air chamber of 30 fertile chicken eggs. Absence of PCR amplification products from these embryo’s template DNAs tested weekly prior to hatching indicated that, as in rabbits, transfer of minicircle kDNA sequences to a bird’s genome required a living *T. cruzi* infection. To determine whether kDNA-transfected birds harbored the *T. cruzi-*specific DNA in the germline cells, we collected sperm from 4938 and 4965, and eggs at an early stage of development from the ovary of 4970 and 4980. DNA templates that were extracted from these samples amplified kDNA but not nDNA (Figure 12). DNA from testes and ovaries of control, uninfected birds 4963 and 4964 did not yield products. Histopathological lesions in muscle tissues and in peripheral nervous systems were seen in offspring that hatched from *T. cruzi-*infected eggs. Myocarditis and ganglionitis were visible, like those lesions described in the rabbit’s offspring [144, 145]. These diagnostic lesions were absent in tissues of control offspring from chicks hatched from uninfected eggs. This rejection of parasite-free target host cells had been observed in adult birds with kDNA integration [111].

**Figure 12.**
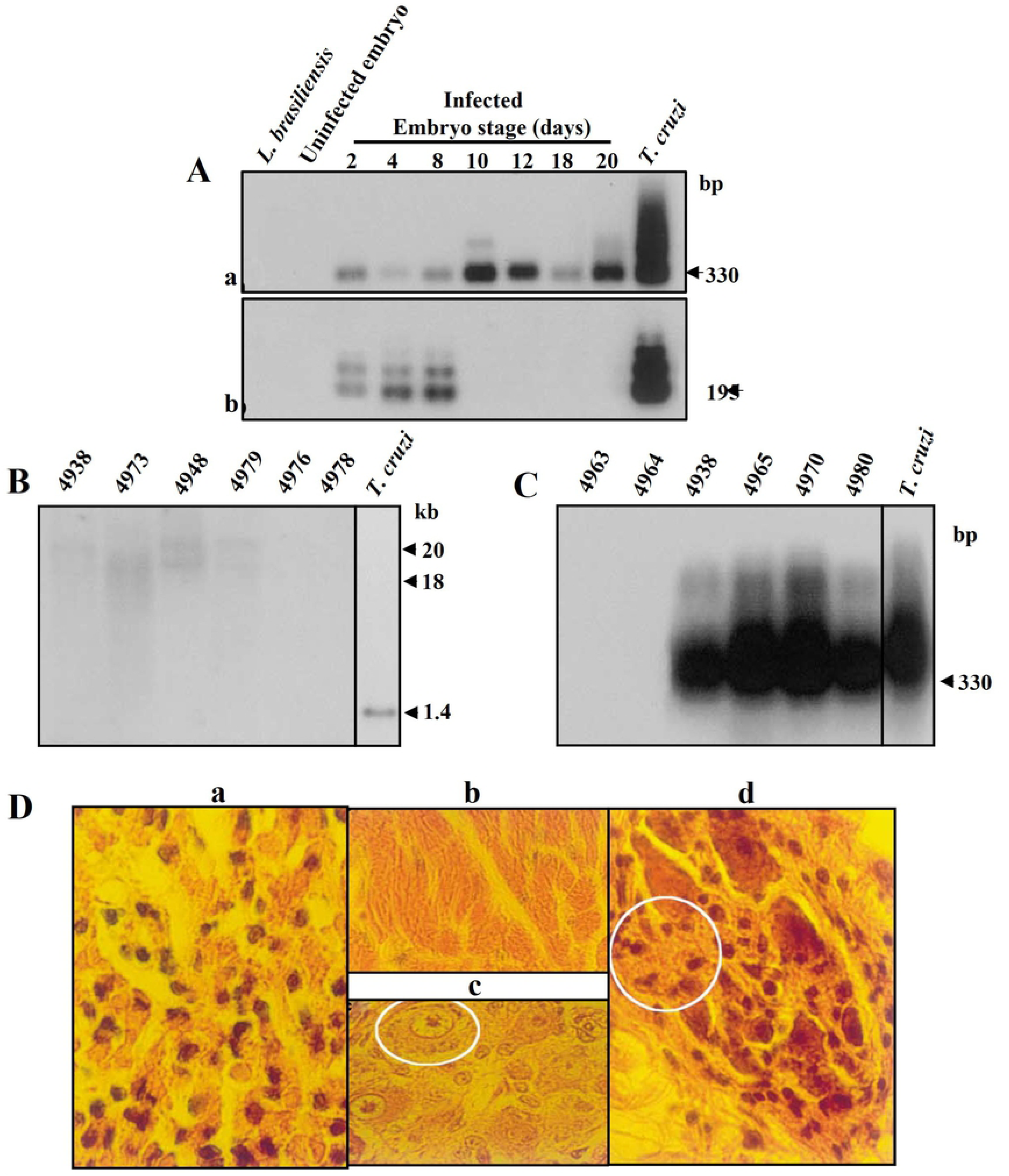
Evidence of kDNA integration in germ line cells and tissues from birds hatched from *T. cruzi-*infected eggs, with accompanying pathology. A) Establishment of *T. cruzi* infection early in embryonic development followed by loss at 10 days of incubation. PCR, size separation, blotting, and hybridization were performed as described in Figure 2a. Aa) Hybridization of PCR amplified bands of kDNA minicircles. KDNA products were amplified from DNAs harvested from tissues at several stages in the embryonic development of the chicken as indicated. Ab) Hybridization of PCR amplification of sequences of nDNA. Bands of 195 bp were diagnostic of nDNA presence in the template DNA. B) Southern hybridization of *Eco*RI digests of genomic DNA derived from *T. cruzi*- infected or uninfected eggs with a kDNA specific probe. Band sizes of approximately 20 and 18 kb formed with DNA from *T. cruzi*-infected birds 4938, 4973, 4984, and 4079 but not with DNA from uninfected, control birds 4976 and 4978. The positive control consisted of 10 ng of *T. cruzi* DNA. C) Integration of kDNA fragments into the avian germline. PCR hybridization analysis of template DNAs from sperm (4938 and 4985) and from nonfertilized eggs (4970 and 4980) from experimentally infected birds, compared to control birds 4963 and 4964. Probe and controls used are describe in B). D) Destructive myocarditis and ganglionitis in 2-week-old F1 Chick 1072. Da), Histopathological section showing mononuclear cell infiltration and lysis of target heart cells, like that described in Figure 9C. Db and c) Normal histological features of myocardium and intracardiac ganglion cells from a control offspring of a noninfected chick. Dd) Section from an intracardiac parasympathetic ganglion showing lymphocyte infiltration and drop out of neuronal cells (circle).

These experiments documented the generation and confirmation of the kDNA minicircle sequences integrated into the chicken germline and somatic cells in the absence of persistent infection. The presence of kDNA with mobility differential from naïve minicircles in the gonadal DNA of individual chickens combined with the absence of *T. cruzi* nDNA attested to the success of the integration event and of the subsequent eradication of *T. cruzi* [35, 56, 57, 172].

#### Germline transmission of integrated kDNA in *Gallus gallus*

With the creation of hens and roosters with integrated kDNA in their ova and sperm, we were poised to examine the transmission of integrated kDNA to the resulting progeny independent of persistent or cryptic *T. cruzi* infection.

The kDNA transfected rooster 4938 and hens 4973 and 4984 were bred to produce vertical, germline transmission of kDNA to their offspring. Twelve chicks born from these crosses carried the kDNA in their genomes, as shown by amplification from blood cell DNA. The kDNA positive F1 birds then crossed to obtain F2 hybrids. Lineages of kDNA transfected progeny are shown in Figure 13. KDNA transfected rooster 4979, and control, uninfected hens 4976 and 4978 bred to detect the frequency of vertical inheritance of kDNA to F1 and F2 offspring from a single kDNA donor parent. All chicks born from these crosses show the kDNA in their genomes thus, indicating that integrated kDNA can be inherited through the male. The cloning and sequencing reveal kDNA integrated in the chickens’ genome (Figure 13C). In a control PCR for nDNA, no bands obtained from any of the offspring DNAs.

**Figure 13.**
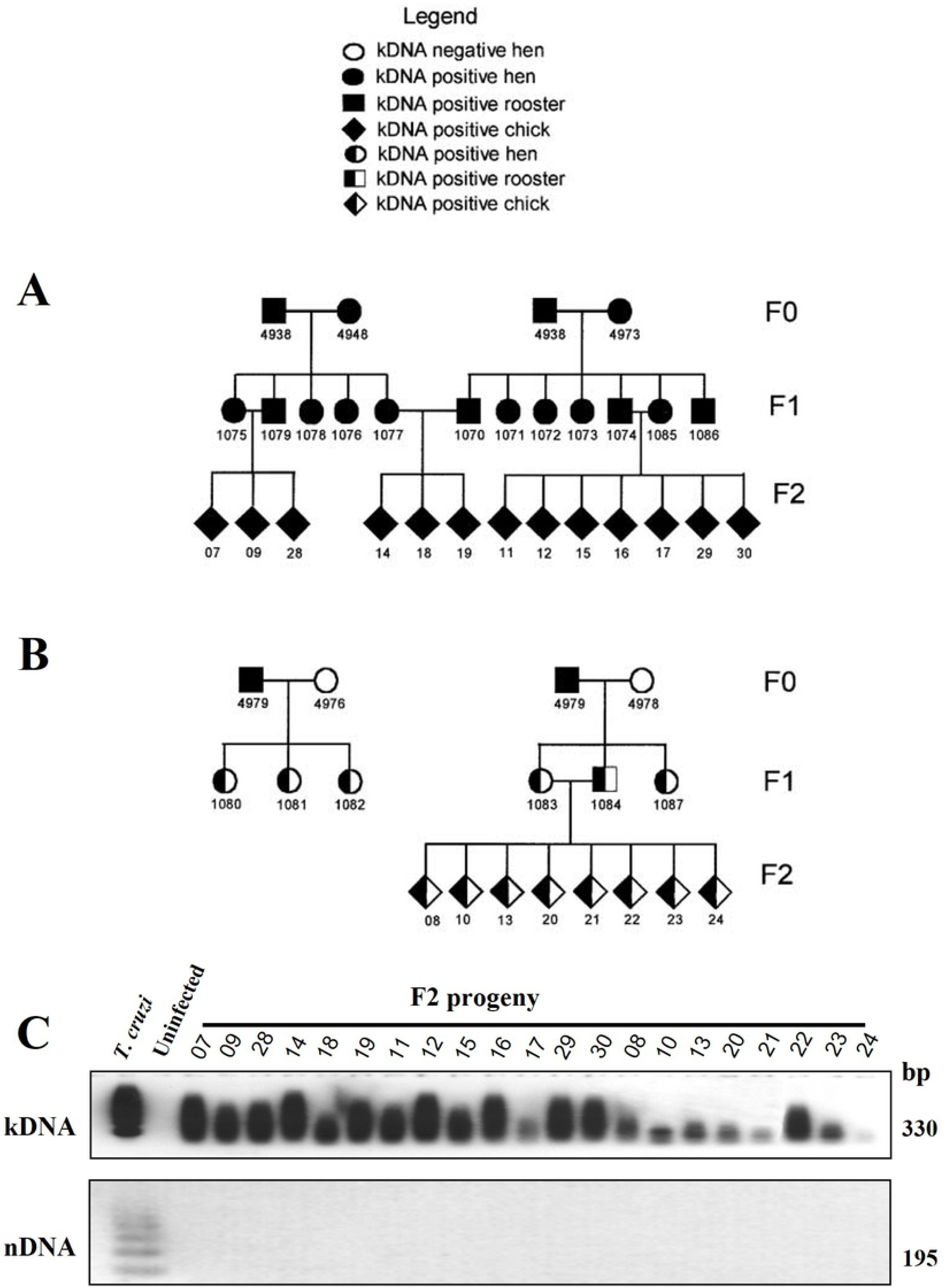
Pedigree of chickens carrying integrated kDNA. A) Parentals kDNA-positive rooster 4938 kDNA-positive hen 4948 (FO) were crossed, yielding five offspring each carrying the kDNA genotype. Similarly, kDNA-positive rooster 4938 cross with kDNA-positive hen 4973, yield seven offspring (F1) each carrying the kDNA genotype. Further crossings of F1 result in offspring each carrying homozygous kDNA genotype (F2). B) Parentals kDNA-positive rooster 4979 cross with kDNA negative hen 4976 (FO), yield three offspring each carrying the kDNA genotype. Additionally, kDNA positive rooster 4979 crosses with kDNA negative hen 4978 yield three offspring carrying the kDNA genotype (F1). Further crossings of F1 positive rooster with negative hen resulted in offspring carrying hemizygous kDNA genotype. C) Evidence of persisting kDNA but not nDNA in tissues from progeny of birds hatched from *T. cruzi-*infected eggs. The PCR hybridization assay performed as described in Figure 1A. Ca) Hybridization of PCR amplification of kDNA minicircles utilizing the kDNA primer set shows band size 330 bp and its catamers in each F2 progeny, which were not formed in uninfected, control DNA. Controls consisted of 10 ng of *T. cruzi* DNA and 20 µg of uninfected chick DNA. Cb) Absence of *T. cruzi* nDNA in progeny of germline kDNA parents. The 195 nts band and its catamers formed with the *T. cruzi* DNA only.

The mapping of the kDNA minicircle sequence integrations into the genome of 35 chickens hatched from the *T. cruzi-*inoculated eggs, which disclosed 72.8% of the lateral kDNA transfer events into the LINE-1 elements at several chromosomes. The databank analysis reveals the kDNA integration with shearing of the transfers to the large chromosomes into the chickens’ genome [6]. The details of those specific *loci* of the integrations more frequently seen in the chicken genomes, showed in the S5 Table.

The separation of the kDNA minicircle sequence integration event from active infection with *T. cruzi* quells any doubt that our observations are spurious. The fertility of roosters and hens harboring specific integration events bodes well for additional studies on the biological effects of DNA acquisition.

#### The Chagas-like pathology in chickens

The gross pathology in the birds hatched from *T.* cruzi-infected eggs consisted of cardiomegaly like that seen in the human Chagas heart disease. The heart showed thickening of the walls of the ventricles and dilation of the chambers. The microscopy revealed myocarditis with infiltrates of mononuclear cells and lymphocyte lysis of the target host cell (Figure 14). In addition, ganglionitis and lysis of the parasympathetic neurons were seen in the heart, in the wall of the intestines and in the skeletal muscles. The finding of the parasite in the heart was unexpected because the chicken’s innate immunity eradicates the *T. cruzi* infection.

**Figure 14.**
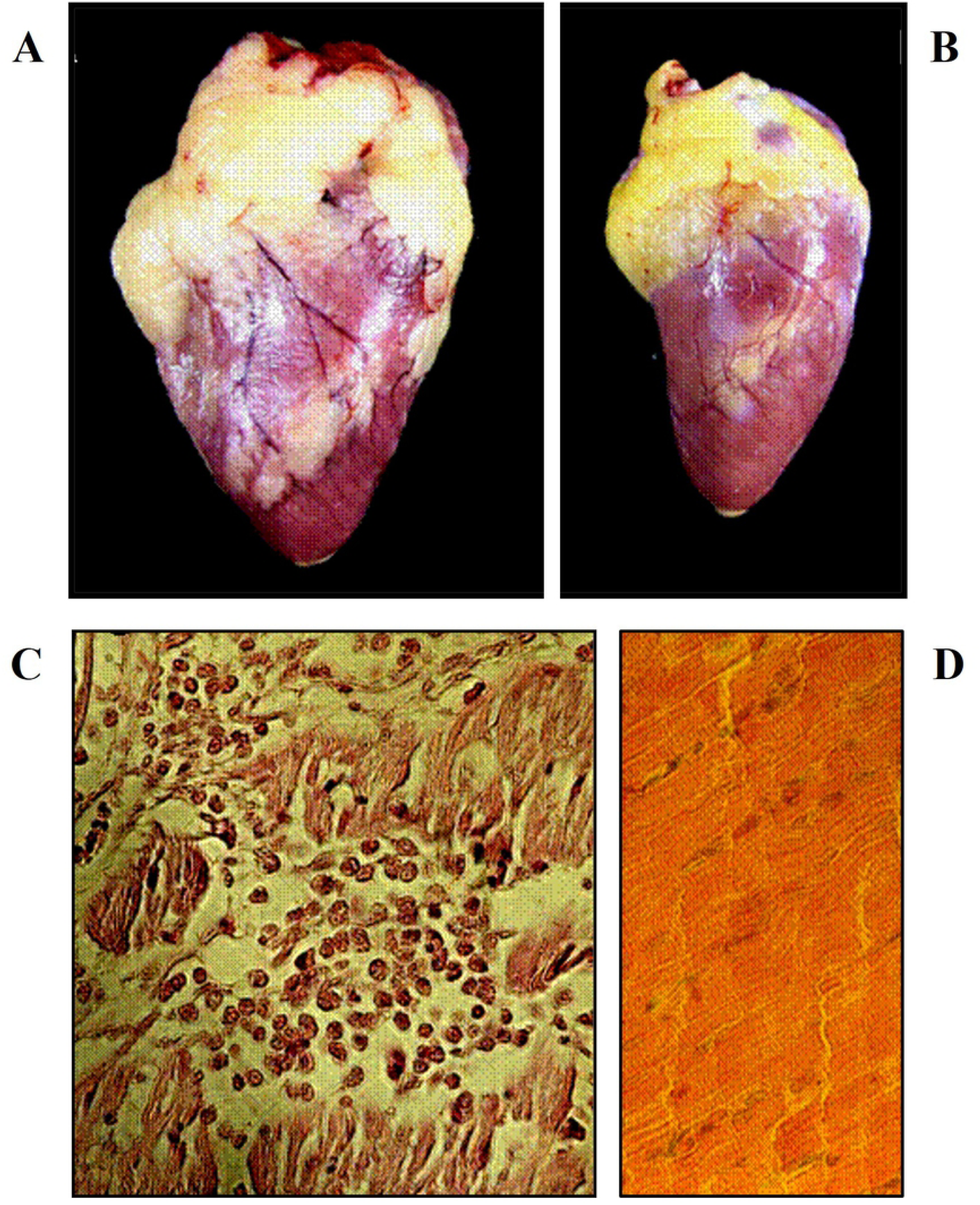
The parasite-free pathology in the chickens hatched from the *T. cruzi-*infected eggs. The conspicuous cardiomegaly (A) in the kDNA positive chicken contrasts with the small heart size of the kDNA negative control chicken (B). C) The histopathology of the heart from kDNA positive chickens hatched from *T. cruzi*-infected eggs shows the typical myocarditis like that of the human Chagas. Note the cytotoxic lymphocytes infiltrate and lysis of the target cells open space for a tunnel in the myocardium of a kDNA positive rooster. D) The normal histology of a control rooster of the same age. The kDNA positive, as well as the negative control chicken that hatched from eggs that did not receive the *T. cruzi* inoculation in the air chamber, in the absence of the amastigote forms in the body tissues [39].

In brief, the experiments showed the persistence of the *T. cruzi* nDNA and of the kDNA into the chronically infected Chagas rabbit genome, whereas the kDNA alone remained in the genome of the chicken refractory to the infection. Moreover, the chickens hatched from the *T. cruzi-*infected eggs developed the Chagas-like heart pathology identical to that described in rabbits. With this respect, the efficacy of any therapeutic regime against the Chagas disease should abrogate the lateral transfer of the kDNA minicircle sequences into the host’s genome, biomarker in charge of the heart pathology.

In view of these results, we initiated the investigation to finding out whether the multidrug treatment of *T. cruzi* infection with the lead Benznidazole in combination with drug inhibitors, which prevented the eukaryote cells division and growth *ex vivo*, could either prevent or lessen the lateral kDNA transfer phenomenon and the pathology in the mouse models of Chagas disease.

### The mouse models

#### The multidrug treatment of *T. cruzi-*infected BALB/c mouse

The employment of the Ofloxacin and of the Azidothymidine to treat bacterial and viral infections and their inhibition of the growth of *T. cruzi* forms without detectable toxicity against the host murine muscle L6 cells, in keeping with the criteria to preventing the transfer of the kDNA minicircle into the macrophage’s genome (Figures 4 to 6), led us to anticipate a therapeutic regime encompassing the combination of these inhibitors (Table 1) with the Benznidazole lead for the treatment of Chagas disease. On the one hand, we used the Azidothymidine that intercalates thymidine to forming the DNA adduct and inhibition of reverse transcription at the G2/M phase, caspase-induced cell division arrest, and apoptosis. On the other, the Ofloxacin 4-Fluorquinolone that inhibits the polymerases metabolic pathway at G2/M phase, and apoptosis. This rational aimed at to unravelling whether the envisage multidrug treatment that curtailed the infection would prevent the lateral transfer of the kDNA integration into the mouse genome, and pathology.

The results of these experiments showed that the independent administration of Benznidazole, of Azidothymidine, and of Ofloxacin to treat each of eight acutely *T.* cruzi-infected BALB/c mice curtailed from 40 to 20 days the presence of the protozoan flagellates in the blood (Figure 15A); and showed further that the curtailment of the flagellates in the blood was achieved with the two-by-two combination of Benznidazole with Azidothymidine, and with Benznidazole and Ofloxacin. Furthermore, the employment of the nitro heterocycle lead in combination with the Azidothymidine and the Ofloxacin, inducers of cell growth arrest and apoptosis, curtailed the parasitemia from 40 to 15 days (Figure 15B).

**Figure 15.**
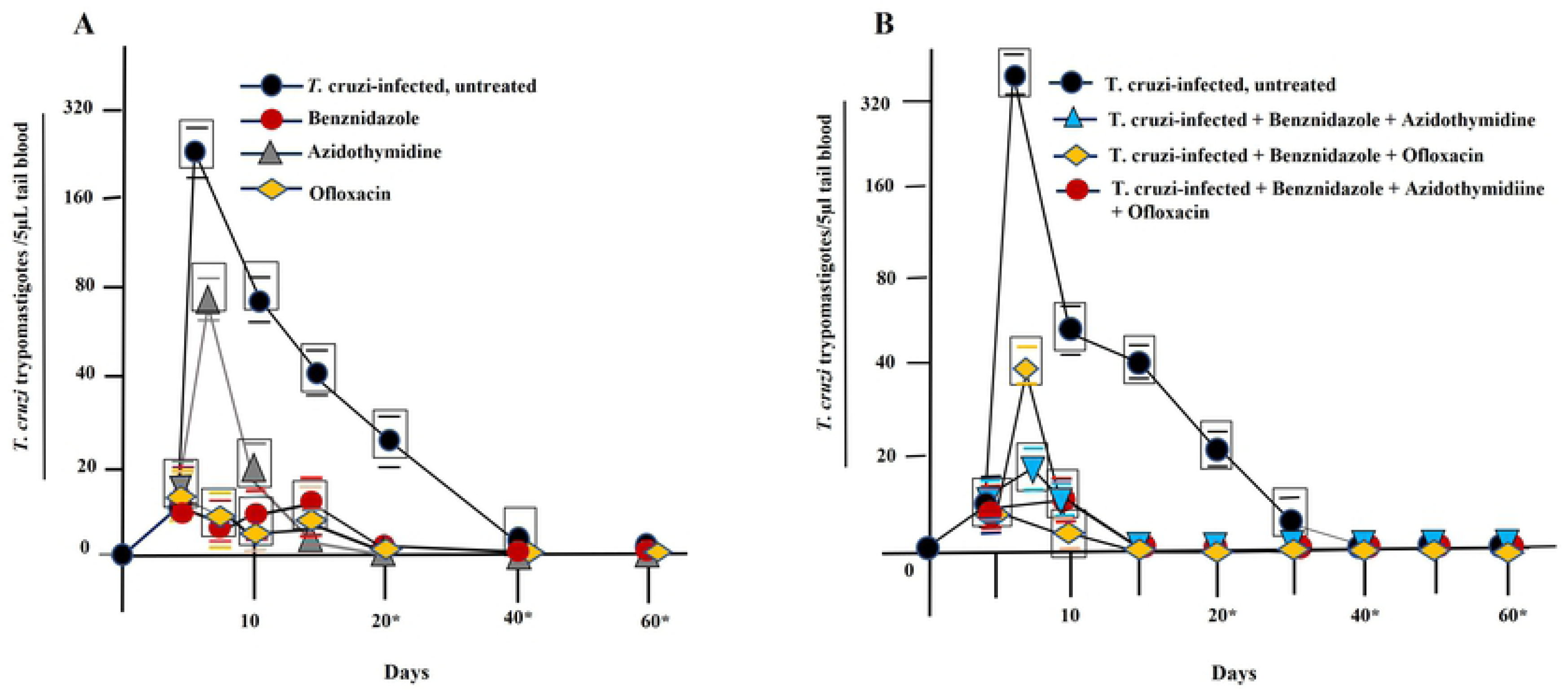
Profiles of the parasitemia in *T. cruzi-*infected mice treated with drug inhibitors of eukaryote cells growth. Groups of eight mice each inoculated with the *T. cruzi* trypomastigotes (2 x 10^3^) intraperitoneal and the multidrug treatment regime by gavage initiated at the 5^th^ day post infection; the *T. cruzi-*infected-untreated was the positive control group. A) The peak of the parasitemia detected in *T. cruzi-*infected-untreated mice, and a quick fall of the parasitemia observed in the *T. cruzi-*infected groups of mice treated with either Benznidazole, or Azidothymidine, or Ofloxacin. B) The parasitemia undetectable at the 40^th^ day in the untreated group of mice, whereas it lasted for 20 days in the *T. cruzi*-infected mice treated with the combination Benznidazole-Ofloxacin or with the Benznidazole-Azidothymidine regime; The results further showed the parasitemia abbreviate to 15 days in the group of mice treated with Benznidazole, Azidothymidine and Ofloxacin. Note the asterisks under the abscissa indicated, however, that the hemocultures set at the 20^th^, at the 40^th^, and at the 60^th^ days post infections yielded *T. cruzi* positive.

At 250^th^ day post infection the mice of all groups sacrificed under anesthesia and the tissues secured by necropsy. The body tissue samples were minced and processed for the DNA purification and analysis. The persistence of the *T. cruzi* infections in the mice of each group was demonstrated further by the PCR amplification with the specific primer sets and Southern hybridization of the products separated in 1% agarose gels, and blot. The results of these experiments clearly demonstrated (Figure 16) the persistence of the infections in the chronically *T. cruzi*-infected mice showing the positive nDNA 195 bp band: the nDNA-PCR triplicate amplification experiments detected a minimum of 15 femtogram DNA, which is 20-fold below the total DNA of the diploid *T. cruzi* parasite [41, 56].

**Figure 16.**
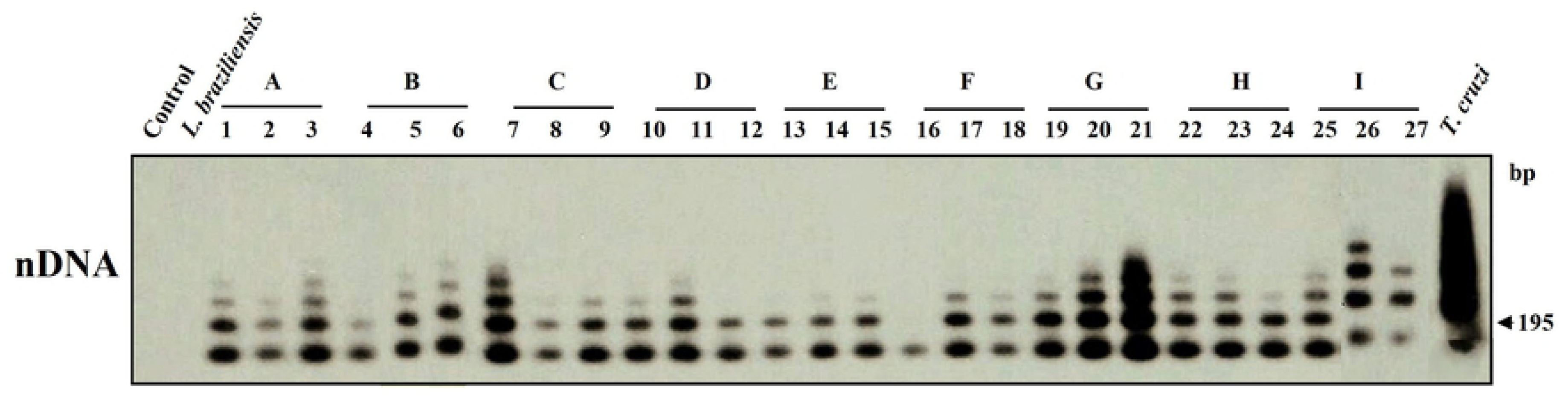
The PCR amplification of the nDNA from the chronically *Trypanosoma cruzi-*infected mice subjected to the multidrug therapeutic regime. Southern hybridization of the *T. cruzi* nDNA-PCR amplification product from DNA extracted from the *Mus musculus* subjected to multidrug therapeutic regime. All the DNA samples yielded the 195 pb bands that revealed the *T. cruzi* infection.

The Ofloxacin and Praziquantel piperazine derivative enantiomers curtailed the *T. cruzi* growth *in vitro*. However, a discrete moderate cytotoxicity of the Praziquantel against the murine muscle cell suggested its exclusion from the therapeutic regime to the *T*. cruzi-infected mouse. In this regard, we continued the experiments in the search for the inhibition of the lateral transfer of the kDNA minicircle sequence to the Chagas mouse genome. The results of the *T. cruzi* kDNA PCR-amplification with the S35/S36 primers and separation of the products in 0.8% agarose gel, blot, and radio label with the specific kCR probe, showed in Figure 17. The presence of the 330 bp band demonstrated some kDNA minicircle sequence integrated into the genome of the *T. cruzi*-infected mice treated with the lead multidrug therapeutic regime.

**Figure 17.** The PCR amplification of the kDNA minicircle sequence from the *T. cruzi-*infected mice subjected to multidrug therapeutic regime. The solid tissues from each group of mice minced with sterile blades, and the mix subjected to DNA extraction and purification. The PCR amplification products separated in agarose gel, blot, and Southern hybridization with the radio label kCR probe. Note the 330 bp kDNA band in the groups of *T. cruzi-*infected mice: *i*) untreated;*ii)* Benznidazole-treated; *iii*) Benznidazole and Azidothymidine treated; *iv*) Benznidazole, and Ofloxacin; and *v*) Benznidazole, Azidothymidine and Ofloxacin. B, blank; C1 and C2, uninfected controls; *T. cruzi*, positive control.

In previous experiments we showed the kDNA minicircle integration into the genome of the *T. cruzi-*infected rabbits and in the chickens hatched from the *T. cruzi* inoculated eggs. The *tp*TAIL-PCR techniques with the specific primer sets employed [41, 55], respectively, to disclose the main site of the kDNA minicircle sequence integrations into the rabbit (LBNL-1) and into the chicken (CR-1) retrotransposons LINE-1 at several chromosomes. The Figure 18 depicted the schematic representation of the *tp*TAIL-PCR employed to disclose the kDNA mutations in the *Mus musculus* genome. The primers set specific to the mouse LINE-1, and the annealing temperature to show in the S6 and S7 Tables. The procedure included the uninfected mouse DNA and *T. cruzi* DNA internal controls, and the last cycle of the *tp*TAIL-PCR reaction showed the chimeras host-*T. cruzi* DNA, only. The A/C rich nucleotides microhomologies at the site of the integration in the mouse DNA, and as well in the kDNA minicircle constant sequence repeats, showed in the **S2 Figure,** which intermediated the end-joining homologous recombination and the lateral transfer of the *T. cruzi* mitochondrion minicircle sequence.

**Figure 18.**
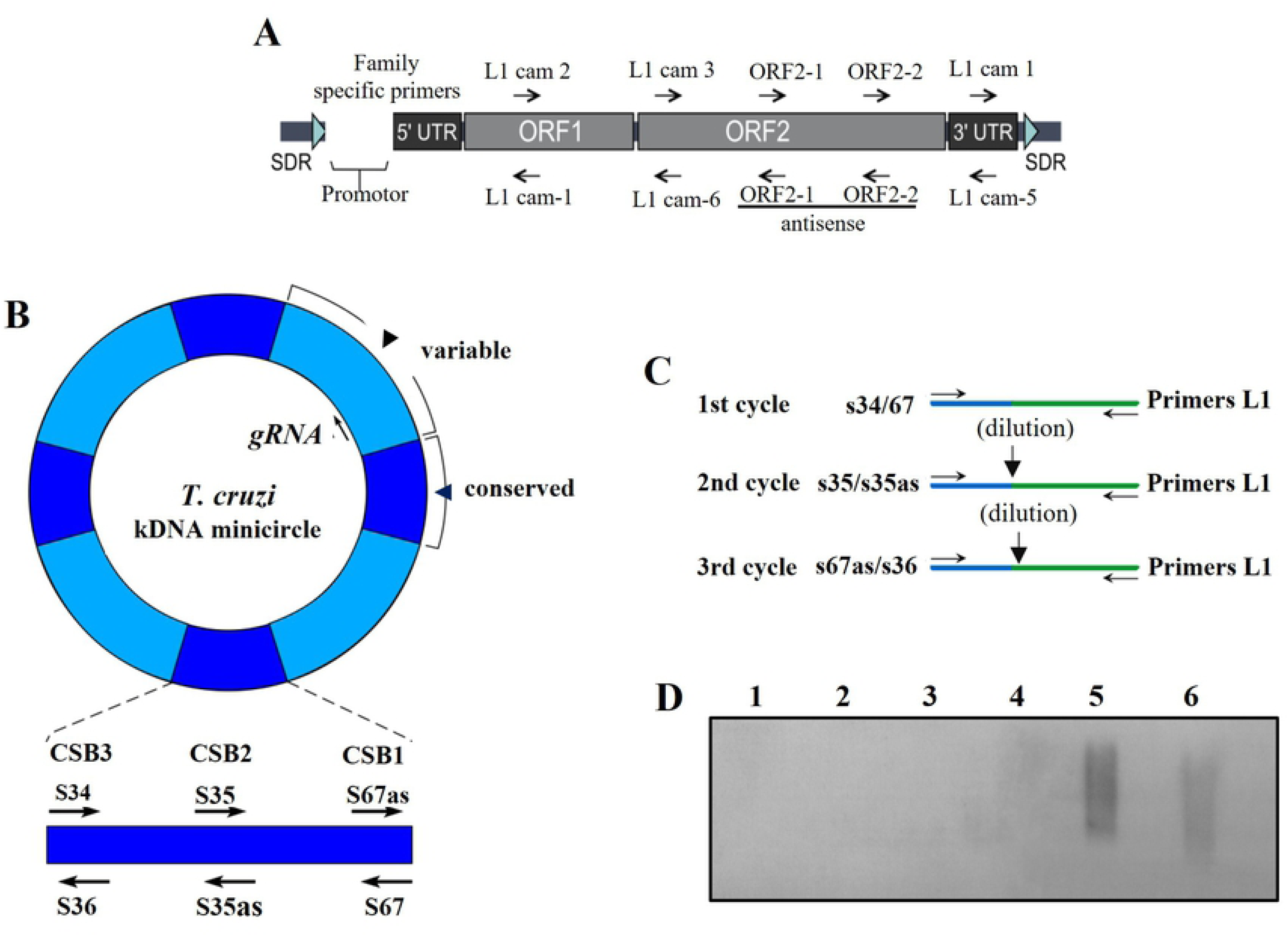
Infographic showing the *tp*TAIL-PCR amplification of the *Trypanosoma cruzi* kDNA minicircle transfer into the mouse genome. A) The scheme showing the *Mus musculus* retrotransposon LINE-1 elements and the combination of primers (arrows) used in the reaction. B) The *T. cruzi* kDNA minicircle showing the constant (light blue) and the variable (dark blue) interspersed regions and the straight demonstration of the CSBs 3, 2 and 1 and primers anneal (arrows) downstream and upstream. C) Reamplification cycles used during the *tp*TAIL-PCR reaction. Notice that each amplification cycle employs the previous cycle amplification product for the subsequent amplification with the kDNA internal primers in combination with the LINE-1 primers shown in the scheme A. D) Validation of the *tp*TAIL-PCR products after the last cycle of amplification. The products of the third cycle are separated in agarose gel, blot, and radio label kCR probe are cloned and sequence. The scheme depicted the 5’ short direct repeat (SDR), the 3’ and 5’ untranslated regions (UTR), the open reading frames (ORF1 and ORF2), and the poli-A tail (A_n)_ of the LINE-1. In addition, it showed the conserved sequence block (CSB), the kDNA minicircle (blue) and the mouse DNA (gray).

Each mice group mix tissue DNA subjected to four independent repeat triplicate samplings *tp*TAIL-PCR and the average non-redundant hybrid sequences, which were obtained from each of five experimental group of mice, showed in the Figure 19. The average non-redundant hybrid sequences obtained with the DNA from the mice group treated with Benznidazole + Ofloxacin + Azidothymidine (n = 18) was 2.44-fold below that much (n = 44) obtained with the DNA mix from the *T. cruzi*-infected, untreated mice. Interestingly, although these data obtained in quadruplicate experiments represented one set point in the course of the chronic infection, the biological implications resulting from overturning the lateral transfer of the kDNA cannot be underemphasized. The cut short inhibition of the lateral transfer of the *T. cruzi* kDNA to the mouse genome suggested that checkpoint inhibitors of the cell growth and differentiation could prevent the lateral transfer of the kDNA into the mouse genome.

**Figure 19.**
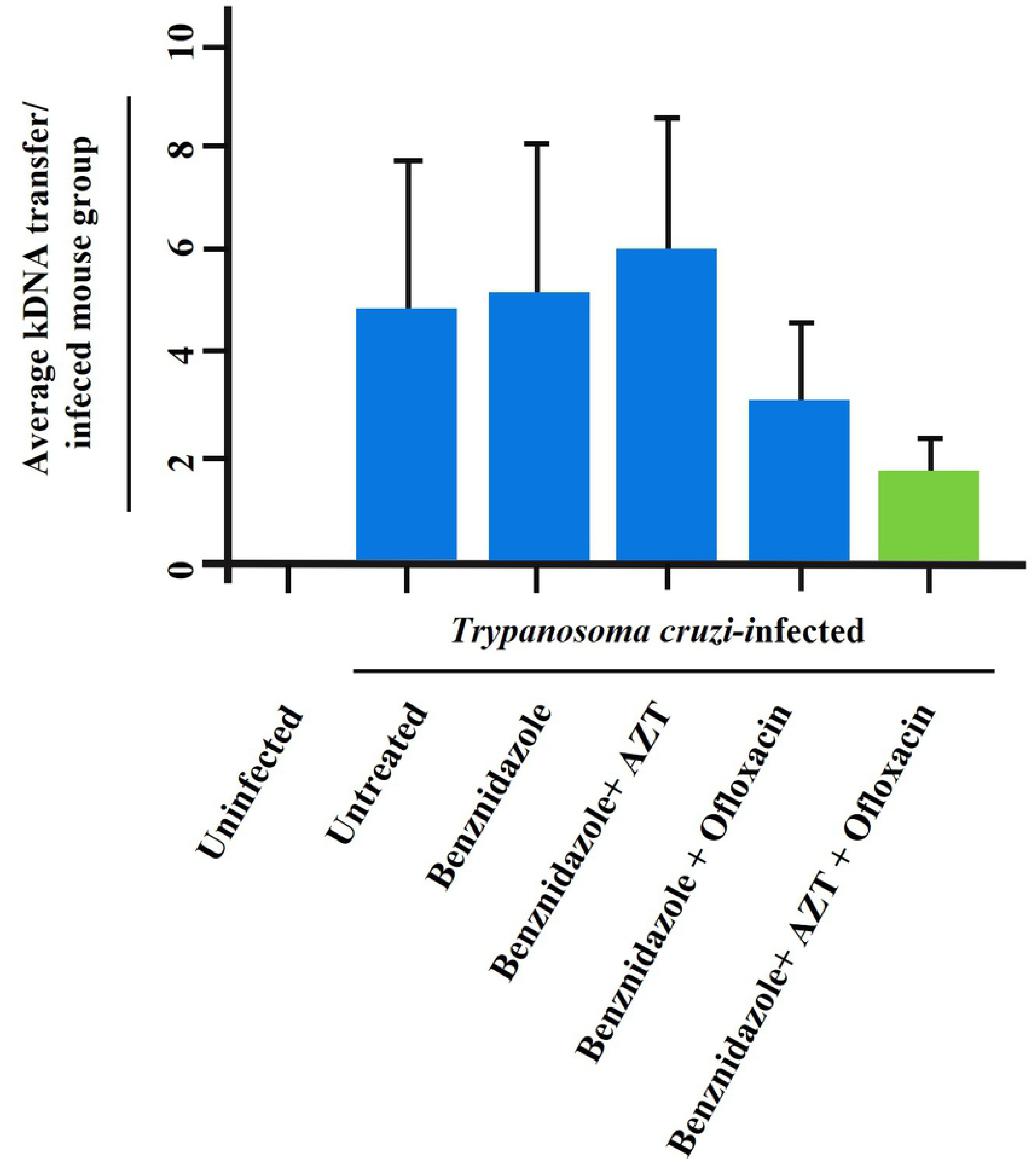
The inhibition of the lateral transfer of the *Trypanosoma cruzi* kDNA minicircle sequence into the *Mus musculus* genome. Schematic representation of the average kDNA transfer in the *T. cruzi*-infected groups of mice treated with the Benznidazole lead in combination with Azidothymidine or with Ofloxacin. Note a 2.44-fold inhibition of the lateral transfer of the kDNA sequence by the multidrug administration of Benznidazole, Azidothymidine and Ofloxacin.

The grand total of 94 hybrid sequences, average size 505 ± 260 nts, displayed in the S8 **Table**. The data revealed that 79% of the lateral transfer of the kDNA minicircle non redundant sequences took place at the Family A, LINE-1 into the genome of the *T. cruzi-*infected mice (Table 2).

**Table 2.**
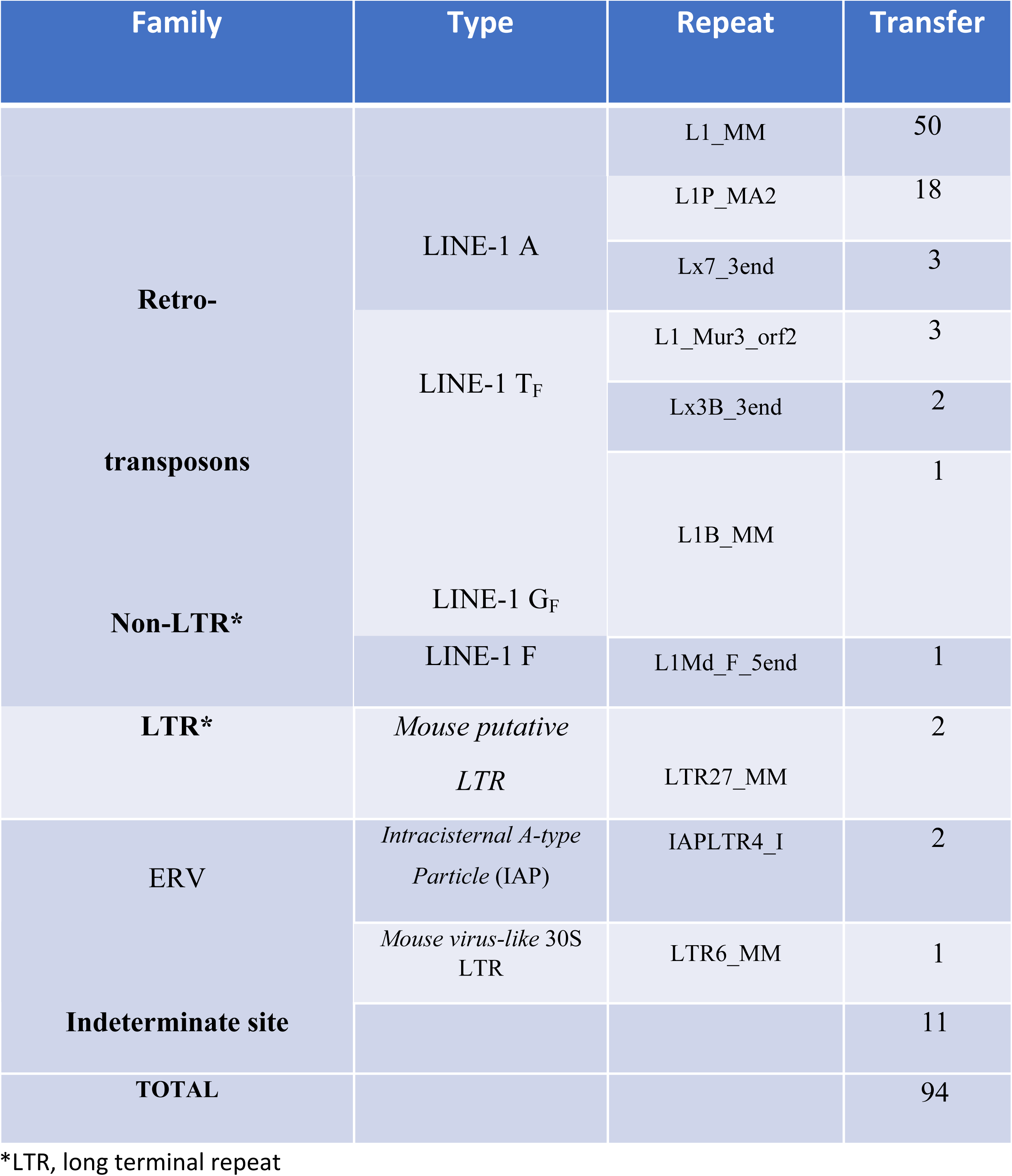
Transfer of *Trypanosoma cruzi* kDNA minicircle sequence and repeat spread into retrotransposon elements of the mouse genome

The development of the Chagas heart disease (Figure 20) in two *T. cruzi*-infected mice and, also, in two mice that had the infection treated with the lead Benznidazole were documented. Moreover, the histopathological study revealed the group of mice treated with Benznidazole + Ofloxacin + Azidothymidine had much less severe myocarditis than the mice treated with Benznidazole alone. The histopathology showed that the Benznidazole, Azidothymidine, and Ofloxacin therapeutic regimens overturned the myocarditis, and a moderate target cells lysis hallmark of the autoimmune Chagas cardiomyopathy, in the absence of the parasite *in situ,* was documented.

**Figure 20.**
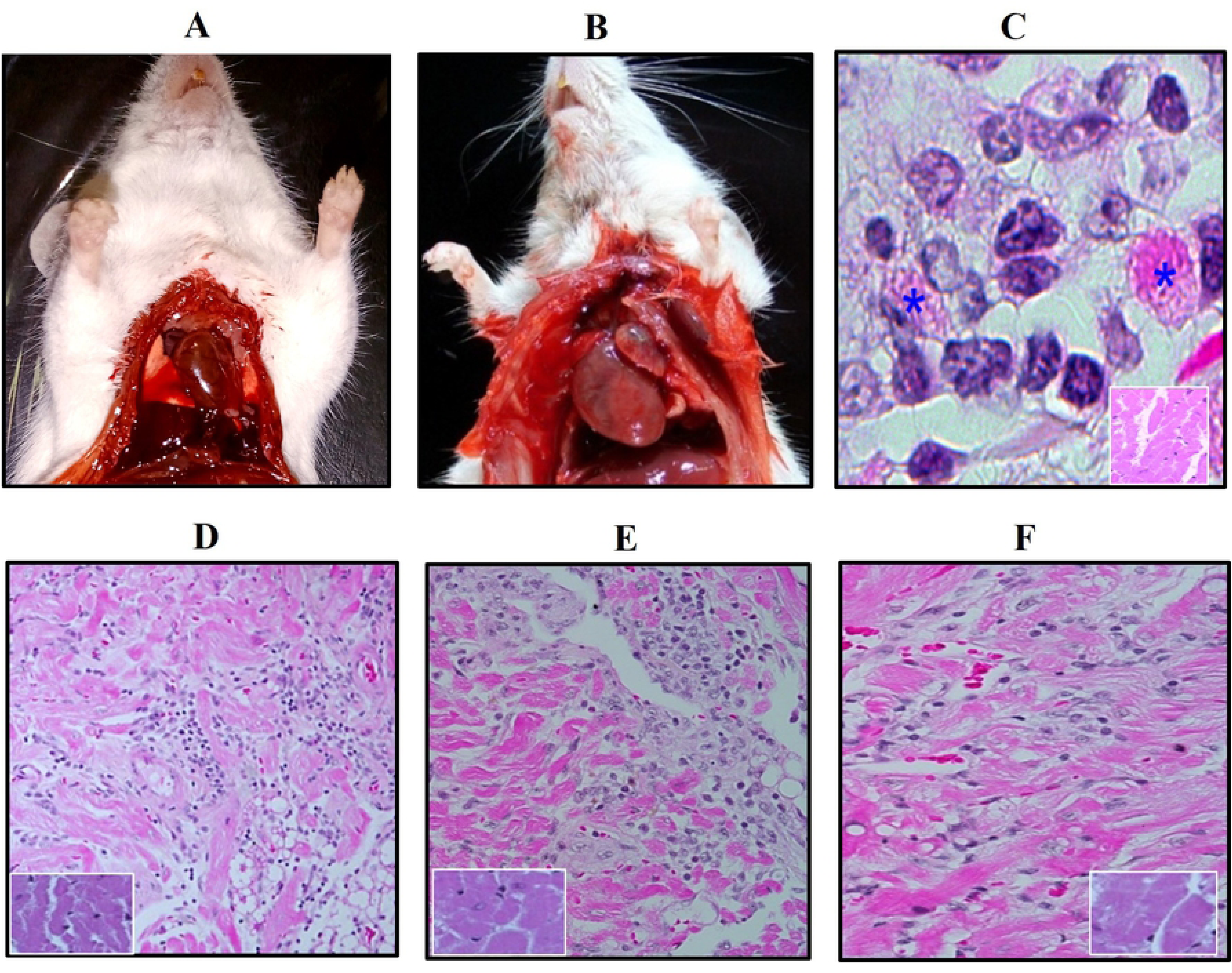
The Chagas heart pathology in the mouse. A) The normal, control adult mouse heart size. B) Dilated cardiomyopathy of a Benznidazole-treated Chagas’ mouse. C) The destructive myocarditis with lysis of the target cells in the *T. cruzi*-infected mouse. D) Myocarditis of a mouse treated with Benznidazole and azidothymidine. E) Ibid, treated with Benznidazole and Ofloxacin. F) Ibid, treated with the Benznidazole, Azidothymidine and Ofloxacin. Notice that the representative histopathological finding in a mouse of group F was consistent with the lessening of the kDNA integrations into the host’s genome.

## Discussion

The prologue of the scientific knowledge on the pathogenic mechanisms calling out the multifaceted clinic manifestations foresaw the transfer of the *T. cruzi* kDNA minicircle sequences could drive the development of the cardiomyopathy hallmark of Chagas disease. With this respect, a strategy compatible with the disease pathogenesis was set forth aiming at its treatment [2, 39, 56, 111]. For in the lack of irreversible drug arrest of the *T. cruzi* infection, we sought for the reduction of the parasite load, and to lessening the accumulation of the kDNA minicircle sequences integration into the host’s genome, therefore, underpinning polygenic modifications that could flare up the parasite-induced, genetically driven mechanism lethal to circa one third of the Chagas people [39, 56, 57]. The postulate addressed to the acute phase of the infection, which subsided spontaneously in the *T. cruzi*-infected, untreated mouse group, followed by chronic phase, and the late myocarditis in the absence of the parasite *in situ* [56, 57]. The observed autoimmune rejection of self-tissue in Chagas disease should be considered to attaining multidrug drug protocols for the treatment of the experimental *T. cruzi* infections, and the prevention of the transfer of the kDNA minicircle sequence into germline and somatic host’s cells.

In view of the demonstration that the nude mitochondrion kDNA minicircle sequences cannot integrate in the macrophage genome, and of the infection stress-induced growth of the host and of the parasite eukaryotic cells [53, 105–110], we decided to test an array of drug inhibitors of metabolic pathways checkpoints. The optimal concentration of each of 12 drug inhibitors was tested *in vitro* against the *T. cruzi* forms. These assays revealed the growth inhibitory efficacy of Bromodeoxyuridine, Azidothymidine Ofloxacin, and Praziquantel in the independent triplicate assays. The trypan blue exclusion dye test showed that these inhibitors of the cells growth and differentiation killed over 75% of the flagellates after 4-day incubation at 27 °C. To finding out the cytotoxicity of an array of 12 inhibitors we incubated each drug with the monolayers of the *T. cruzi*-infected L6 murine muscle cells fed culture medium at 37 °C temperature in a 5% CO2 incubator. The results showed the severe cytotoxicity of the Microcystin phosphatases inhibitor [124–126], and of Mitomycin [127–128], and of Bromodeoxyuridine [117–118] cross linkers forming DNA adducts by covalent attachment to the target macromolecules. The severe cytotoxicity against the eukaryotic cells under the microscope revealed loss of the cell’s membrane boundaries, foamy cytoplasm, and death.

Independent assays that searched for the integration of the *T. cruzi* kDNA minicircle sequences into the human macrophage genome showed that an array of the polymerase’s inhibitors and, also, arrest of the cell division prevented the transfer of the minicircle sequences and no longer the presence of the 1.2, 1.8 and 2.2 kb bands, different from the 330 bp mitochondrion kDNA band. Of interest, we noticed that the Microcystis inhibitor hold back the 330 bp band, but it did not prevent the integration of upper 1.2, 1.8 and 1.2 kb kDNA minicircle sequence bands into the macrophage genome, different from the Camptothecin and the Bromodeoxyuridine that did not prevent any of those bands at all. Nevertheless, the Microcystin serine/threonine protein phosphatase inhibitor-induced cell death should undergo further study towards the development of a new drug to kill the *T. cruzi* agent of Chagas disease.

To approaching the multidrug treatment of the *T. cruzi* infections we selected the polymerases inhibitor 4-Fluorquinolone Ofloxacin [136–137] and the Azidothymidine reverse transcriptase inhibitor [114, 115], and the lead nitro heterocycle Benznidazole. The treatment of the acutely *T. cruzi-*infected BALB/c mice with the Benznidazole curtailed the parasitemia at the 15^th^ day of the drug administration, whereas the flagellates lasted for 40 days in the blood of the infected-untreated mice. Also, the *T. cruzi-*infected mice showed the hemoculture, the ELISA assay for the specific antibody, and the *T. cruzi* nuclear DNA-PCR yielded positive results, consistently, in the infected mice that had received thrice as much the dose of Benznidazole used to treat the human infection [149]. Two out of eight mice each in the *T. cruzi-*infected-untreated and in the infected-treated groups developed the identical Chagas heart disease and died with the dilated cardiomegaly due to a severe myocarditis. This caveat provides the information showing that the administration of the Benznidazole to treat the *T. cruzi* infections is unsatisfactory. The information is in keeping with the observations stemmed from experimental protocols in animal models [144–149] and, accordingly, with the data obtained from human Chagas disease clinical trials [156–162], although the controversy remains [161–169]. In addition, the acutely infected mice permitted parasitemia for 10 days after the administration of Benznidazole + AZT + Ofloxacin.

### Insights into human Chagas disease

The comparative pathology experiments caried out in the chicken refractory to the *T. cruzi* and in the rabbit amenable to the infection revealed that the pathogenesis of the Chagas disease stems from the lateral transfer of the kDNA-inducer of the genetically driven autoimmune myocarditis and target cell lysis by the lymphocytes [56, 57, 111, 145–147, 170–172]. The identical heart pathology in the chicken refractory to the *T. cruzi* infection, in the rabbits, and in the mouse susceptible to the persistent parasite infection was essential to demonstrate the lateral kDNA transfer triggers the pathogenesis of the severe Chagas disease lesions in the vertebrate animal models [35, 172], whereby clones of the kDNA mutated lymphocytes rejected the target heart cells. The study continued to disclose overturning of the late outcome of the Chagas heart disease by the employment of a multidrug cocktail of inhibitors of the main metabolic pathway checkpoints of eukaryotic cells’ growth and differentiation, and the prevention of the lateral transfer of the kDNA minicircle sequence into the host’s genome. Firstly, we found out that the *T. cruzi*-infected mice treated either with Benznidazole and Azidothymidine, or with the Benznidazole and Ofloxacin remained susceptible to the integration of the kDNA sequence into the genome. Secondly, the treatment of chronically *T. cruzi-*infected mice with a cocktail of Benznidazole with Azidothymidine and Ofloxacin showed a 2.44-fold lessening of the kDNA minicircle sequences into the mouse genome. Accordingly, we documented that the mice in this multidrug-treated group had less severe myocarditis than those in the *T. cruzi-*infected and treated with Benznidazole alone.

The lateral transfer of the kDNA minicircle sequence is a natural consequence of the *T. cruzi* infection in the vertebrate host cell genome. The intracellular invasion by the *T. cruzi* stress-induced metabolic burst and eukaryotic cell growth is required for minicircles to integrate into the vertebrate host’s genome. Therefore, intracellular growth and differentiation is included among the environmental factors associated with kDNA transfer, integration, and reshuffling the genome of the host species. The vertebrate genomes contain repetitive long and short elements (LINEs and SINEs) that persists by vertical transmission within a host, and the integration of kDNA sequences into these retrotransposons has implications for further mobilization of foreign DNA within the genome. The vertebrate genome contains variable number of copies of retro transposable elements. The human has circa 400 copies of active LINEs belonging to different subsets, whereas the mouse contains over 3,000 copies belonging to various family and type subsets [173, 174]. These elements are likely progenitors of mutagenic insertions and providers of means for the mobilization and reshuffling of DNA sequences around the genome.

### Cross-Kingdom DNA transfer, evolution, natural selection and Chagas disease

From a historical perspective the lateral DNA transfer phenomenon between organisms of distant evolutionary relationship is equivalent to the events of migration of endosymbiotic prokaryotes organelles to the eukaryotic cells, since circa 1.5 billion years ago [175–177]. The Protist mitochondrion kDNA is considered the remains of microorganisms that invaded the then emerging single celled protozoan with nucleus and was ultimately adapted to become its own component [175]. Our data show that the eukaryotic cell DNA can be frequently transferred from the endosymbiotic mitochondrion to the nucleus. The investigation documented a robust flow of DNA among different creatures of far apart kingdoms. The specific detection of the *T. cruzi* kDNA minicircle fragments integrated into the genomes of vertebrate hosts is reminiscent of the microorganism assimilation of organelles such as mitochondrion and chloroplast [175–178], a prevailing mechanism of evolution at the molecular level. These observations provided compelling evidence of lateral and vertical kDNA transfer stemmed from the experiments. The heritable fashion through germ line cells Cross Kingdom genome growth sustained, certainly in our laboratory.

The integration of the protozoan kDNA minicircle sequence into the germline of rabbit, chickens and mouse cells’ LINEs and SINEs transposable elements at several chromosomes are usually neutral mutation but hitchhiking to coding regions require further studies of DNA sequence modification and evolution; and the creation of new genes, pseudogenes and silencing of genes may produce polygenic alteration and genetic mechanism of disease [179–183]. The sheer number of minicircles, each with four conserved regions containing CA-rich sequence motifs, may be the dominant characteristic influencing the high frequency of this serendipitous mutagenic events that may influence the endogenous host genes [184, 185]. The transposable elements that replicate within the host genome may contribute to somatic mosaicism and, thus potent regulatory polygenic modifications in the diseased state [186, 187] play a role in the pathogenesis of inflammatory autoimmune Chagas disease [35, 41, 57, 172, 188]. In so far, the polygenic modifications involved in the genesis of cancer [189–193] have not been ascribed to any Ambiental factor inducer of the lateral exogenous kDNA transfer. In this regard, evolution is inherently opportunistic.

We postulate that the kDNA insertion mutations underpinning phenotype alterations of the effector and of the target host’s cells undergone polygenic modifications, stemming from the LINE-1 elements intermediate genome reshuffling, called out the *T.* cruzi- induced autoimmune driven Chagas disease lesions over time [33, 36, 39, 41, 57, 114, 172]. Although the transfer of mitochondrion kDNA events are neutral for the sake of the species evolution, some mutations that could be deleterious hindered identification because the lifelong chronically infected individual usually retains normal health. The threatening heart disease flareup in circa one third the chronically infected individuals bore short survival of two to five years. The timely investigation of those Chagas heart disease cases revealed accumulation of lateral transfer of the kDNA mutation range from a minimum of 4 to 8 into the LINEs located at various chromosomes [41].

The sexual transmission of the *T. cruzi* infection from males and females to naïve mates and reshuffle to the family members has been documented throughout subsequent generations [33, 36, 57]. The Chagas family born acute infections usually go unrecognized. The epidemiologic studies to unravelling an on-going Chagas disease pandemic shall consider that the *T. cruzi-*infected youngster usually does not become sick at a time when tissue parasitism is high but may develop striking lesions several decades later when the parasite is difficult to find. Certainly, disease manifestation may not be dependent on the parasite’s direct action upon a host cell, and autoimmune rejection is likely to play a role in the pathogenesis of the late Chagas disease.

The role of autoimmunity in Chagas disease has been examined in experiments carried out in the chicken refractory to the *T. cruzi* [41, 57]. The experiments showed that the innate immunity of chicks hatched from the *T. cruzi*-infected eggs eradicate the parasitic infection at the end of the first week of the egg incubation. Interestingly, the kDNA minicircle sequences remain integrated in the chicken genome, and the mutations in the parental and progeny were found in the chicken LINE-1 repeats sheered to several chromosomes. Furthermore, the kDNA mutated adult chickens develop the parasite-independent Chagas-like dilated myocarditis identical to that seen in the Chagas heart disease cases in humans and died.

The refractoriness of the Aves to the *T. cruzi* flagellate protozoan was timely convenient for this study aiming at the inhibition of the autoimmune Chagas heart pathology because it did not require precleaning the infection. In view of the immune tolerance observed in the progeny of the Chagas disease parental, in the absence of the specific antibody, the traditional mechanism of target cell destruction by antigen-specific antibody acquired immunity does not hold up [41, 57]. Otherwise, the organ specificity of the autoimmune phenomenon that targets the heart is the result either of germline or of somatic mutation in the genome of the cytotoxic lymphocyte and of the target, heart cells. With this respect, each kDNA-integrated immune system mononuclear cell involved in the “self” tissue destruction is essentially a mutated clone homing to the target organ where the cytotoxic lymphocyte promoted the lysis of the target cells. The disadvantageous kDNA mutations have been encountered in the Chagas disease late phase histopathology evidenced in the experimental rabbit and chicken model systems of the heart disease [33, 36, 57, 61, 145–149]. We suggest that the kDNA insertion could produce polygenic alteration and phenotypic modification of mutated host cells is the factor triggering the parasite-induced autoimmune tissue-specific rejection in Chagas disease. Alternatively, in the absence of deleterious polygenic alteration and phenotype modification, it could explain the long-lasting asymptomatic infection in a major parcel of the *T. cruzi*-infected people, and in some patients with mutagenic kDNA fragments dispersal by active LINE-1 mobilization within the genome and reversal of the clinical manifestations, as documented in family-based study [162].

Two questions wait for comprehensive knowledge stemming from the molecular biology of evolution and natural selection spurred transfer of the Protist’s kDNA into the vertebrate’s genome. Firstly, we deal with the organ specificity of the muscle and of the parasympathetic neuronal cells targets of the genetically driven autoimmune rejection. Secondly, the homing of the bone marrow derived lymphocytes towards the counterpart kDNA mutated target cells. We postulate that the selective effector-target cells affinity could be a genetically acquired trait dependent upon the *T. cruzi* kDNA minicircle with four interspersed constant and variable sequences, showing the CSB1, CSB2 and CSB3 AC-reach repeats intermediating the lateral transfer of the kDNA to circa 850 homologous repeats, average 8 to 25 nts share into the human genome [33, 36]. Additionally, the share repeats intermediate, bending and loop forming tri-dimensional structures, binding sites for charged protein electron emitted signaling pathway, and a closed lock selection site of the foreign DNA transfer end-joining recombination mechanism [36, 41 and 57]. The specificity of the target organ rejection now, could be called out through the AC-reach repeats stoichiometry thus, bend and loop tri-dimensional structures [194], source of cell-to-cell signaling the homing roadway [33, 36, 39, 41, 57, 172]. Occasionally, groups of kDNA minicircle sequence integration mutations, and polygenic modifications thereof, may explain the lymphocyte activation and the rejection of the target host’s cells, showing the multifaceted clinical manifestations translated into the pathology. The studies of chromosome skewing and instability-generated from signaling interactions may substantiate the genetically driven mechanism ensuing the rupture of immune tolerance and to explaining the eventual attenuation due to chromosome upright remodeling.

The practical value of animal models basic research supported the treatment of the Chagas heart disease by bone marrow transplantation [57]. The experiments aimed at the detection of parasite-independent heart homograft rejection in congenic kDNA mutated chicken strains showed that the killing of the bone marrow from the kDNA-positive chickens with the cytostatic chlorambucil and replacement with the bone marrow cells of the histocompatibility chicken strain prevented the Chagas heart lesions described in the bird’s model system [57]. Moreover, an advantage of the chicken model is the provision of new insights that showed the genetically driven autoimmunity process in the mammalian required the combination of the nitro heterocyclic with the inhibitors for the multidrug treatment to halt the genetically driven Chagas heart pathology, since the *T. cruzi* eradication by the bird’s natural immunity is evident [35, 36, 172].

The kDNA insertion mutations could play different functional roles ranging from advantageous, neutral, and disadvantageous to the host. We shall emphasize studies of the advantageous kDNA mutations that could associate emergence of adaptive character that may be rapidly driven toward fixation by Darwinian natural selection [176]. The long-lasting, cryptic *T. cruzi* infection of vertebrate hosts is consistent with neutral mutations resulting from lateral kDNA transfer and its vertical inheritance in a vast number of offspring that could represent a prevailing mechanism of evolution at the molecular level. Neutral kDNA mutation, while having no recognizable benefit to the host, can provide a substrate for natural selection. This type of mutation could play a major role in evolution mainly by loss, alteration, and refinement in certain groups of organisms subjected to environmental conditions over the evolutionary time. Furthermore, we show Cross-Kingdom molecular chimeras created by the lateral transfer of a Protist’s kDNA, hitchhiking, rearrangement and reshuffling of the eukaryotic host genome, therefore, an unexplored force in speciation [177]. The DNA mutation resulting from a persisting infection could explain the spectrum of Chagas disease manifestations seen several decades after the onset of the *T. cruzi* infection and surreptitious phenotype alterations that contribute to the activation of CD8α+ and CD8ß+ effector cells that express vß1 and v ß2 receptors involved in the target cell rejection [57].

### Perspective

The reproducibility validated experiments reported herein shall foster new research tools for the understanding of the biology of the Chagas parasite and of the chronic heart disease. Much fundamental research is needed to achieve the prevention and control of the forthcoming family-based pandemic disease. In view of the immune tolerance documented in cases of the vertical transmission of the *T. cruzi* infection the antibody assays may not contribute to the disease surveillance and control. The accuracy of the diagnosis of the *T. cruzi* infections requires the nucleic acids assays. The development of a thorough put platform is needed for the accurate diagnosis in hospitals and clinical outposts. In addition, the prevention and control of Chagas disease requires much paradigm research with new approaches and innovation technologies to repurposing of inhibitors of the protozoan parasite metabolic pathways [195]. The development of a new drug [196] devoid of toxic effect and proved to eliminate the cryptic chronic infection is essential to obtaining the control of the Chagas disease spread worldwide [197–200].

For the disease spread worldwide requires surveillance-based education program to control the silent pandemic Chagas disease. An inevitable question is set forth: how to explain the autochthonous Chagas disease acquisition [199–206] in the Northern Hemisphere? Notably, the sexual transmission of the *T. cruzi* introduced substantial changes in the concepts of the health care, control, and surveillance of the Chagas disease [36]. In this regard, we postulate that some genetically driven autoimmune idiopathic dilated cardiomyopathy could stem from vertically inherited kDNA-like mutation. The odds for the vertical transfer of the parasite-induced kDNA mutation, underpinning the genetically driven autoimmunity, with high ratios of morbidity and mortality indicate the roadway for the multidrug treatment and prevention of the kDNA mutation genomic alteration and, possibly, the polygenic modifications that could call out the pathogenesis of the Chagas disease. Meanwhile, the prevention and control of Chagas disease shall rely on the sustainable education, information, and communication program for health.

## Materials and Methods

### Growth of the eukaryotic cells

The archetype Berenice *T. cruzi* (Chagas 1909) [35] grown in 5 ml of Liver-Infusion Tryptose (LIT) medium in screw-cap glass tubes kept on a shaker-incubator at 27 °C, and a clone of the Tulahuén *T. cruzi* stock expressing β-galactose [207] grown in Dulbecco Minimum Essential Medium (DMEM) with 20% fetal bovine serum (FBS), in 5% CO_2_ atmosphere, were used. The *T. cruzi* epimastigotes grown in the axenic LIT medium employed to determine the optimal doses of the drug inhibitors of the cell growth and development (Table 1). The *Leishmania brasiliensis* LTB300 stock promastigote forms harvested in the exponential growth phase were collected from the culture medium; the cells washed trice in PBS, pH 7.4, were collected by centrifugation and used in the experiments [35].

The L6 murine muscle cell and the U937 human macrophages purchased from the Bio-Rio Foundation, Rio de Janeiro. The cell lines were maintained by serial passages in sterile culture flasks containing freshly prepared DMEM, pH 7.2, with 10% FBS, 100 µm/mL penicillin,100 ug/mL streptomycin, and 250 nM L-glutamine at 5% CO2, at 37 °C (complete DMEM). The L6 murine muscle cell line grown in 15 mL culture flasks were inoculated with 10 x 10^4^ trypomastigotes that underwent differentiation to amastigote-dividing forms into the host cell cytoplasm. After several cycles of division each 14 hours the host cell-loaded amastigotes differentiated the trypomastigotes that burst-out in the supernatant medium. The flagellates in the supernatant collected by centrifugation at 3,000 x g for 15 min in a swing bucket rotor were washed trice, resuspended in FBS-free DMEM, and quantified in a counting chamber. The free-swimming trypomastigotes were used in the *ex vivo* drug inhibitor experiments and, also, to infect the experimental animal models of Chagas disease.

### Collection of Human Blood

We collected 10 ml of blood from Chagas patients who volunteered to participate in this study. Serum samples were subjected to immunological testing for specific antibodies, and the blood-nucleated cells yielded DNA for biotechnological analyses [35].

### Interactions of the macrophage with the *T. cruzi* kDNA minicircle sequence

In the kDNA minicircle experiments, aliquots of 10 x 10^4^ uninfected macrophages suspended in 10 mL SFB-free DMEM were incubated with: *i*) 5 µg of lipofection pBlueBacHis2/CAT plasmid (Invitrogen); *ii*) DNA from uninfected, control macrophage; *iii*) crude *T. cruzi* kDNA; *iv*) cloned kDNA sequence. Each aliquot then incubated with the macrophage monolayers in flasks with complete DMEM, at 37 °C, at 5% CO_2_ atmosphere. At set-points 1- 7-, 14- and 21- day post-incubation the cells were harvested for the DNA analyses.

In the other experiment, 10 x 10^4^ uninfected macrophage fed the complete DMEM at pH 7.2, incubated at 5% CO_2_ atmosphere, at 37 °C, forming monolayers in the culture flasks were inoculated with 50 x 10^4^ *T. cruzi* trypomastigotes suspension in 15 mL of the DMEM. At set-points day-7 and day-31 post-infection the macrophages released with 0.25M trypsin, washed thrice with PBS, pH 7.2, and subjected to DNA extraction.

### The assessment of cytotoxic effect of drug inhibitors on the *T. cruzi-*infected muscle cells

The L6 muscle cells monolayers grown in glass chamber (Tissue-Tek, Thomas Scientific) were inoculated with 10 x 10^3^ trypomastigotes aliquots suspended in 3 mL DMEM. One week after the *T. cruzi* inoculation the muscle cells loaded with the dividing amastigotes received a concentration of a specific cell growth inhibitor. The drug inhibitors of eukaryotic cells check-point showed in Table 1. The chemical structure of each inhibitor shown in the S1 Figure. In these experiments, the drug concentration determined in dose response assays, which showed either absence, or minimal, or severe cytotoxic effects on the *T.* cruzi-infected L6. The optimal concentration of the drug inhibitor did not produce pathogenic effect on the L6 muscle cells identified by the search under an inverted microscope, during two weeks in a CO2 incubator at 37 °C. The images showing cytotoxic effects of some inhibitors on the host cells were captured with a DP72 camera coupled to an Olympus BX51 microscope.

### Hosts cells’ DNA extraction

The macrophages and the L6 muscle cells monolayers subjected either to the *T. cruzi* infections or to the lipofection with different sources of DNA sequences were collected by centrifugation and washed once with PBS, pH 7.4, After two washings with TBS (20 mM Tris-HCl pH 7.2, 0.5M NaCl) the cell pellets obtained by centrifugation at 1.500 x g for 15 min and resuspended in 2 mL of extraction buffer (1 mM Tris-HCL pH 8.0, 0.1M EDTA pH 8.0, 0.5% SDS, RNase, 200 ng/mL). Thereafter, the proteinase K (100 µg/mL) added, and the incubation period extended for 12 h. The cells in the pellet were subjected to two extractions with equal volume of chlorophenol (phenol: chloroform: isoamyl, 25:24:1) and a further extraction with chlorophyl (chloroform: isoamyl alcohol, 24:1) under light agitation at room temperature. The aqueous and organic phases separated by centrifugation at 5,000 x g for 10 min, and the DNA precipitated with 1:10 volume of sodium acetate 3 M, pH 4.7 and 2.5 volume of cold 100% ethanol. After 30 min stand at –80 °C the DNA was sedimented by centrifugation at 12,000 x g for 15 min. The DNA washed twice with cold 70% ethanol dried out before resuspension in TE buffer (10 mM Tris-HCl, pH 8.0, 1 mM EDTA, pH 8.0). The DNA subject to the analysis in 1% gel electrophoresis was stored at –4° C.

#### Trypanosoma cruzi and Leishmania brasiliensis nuclear DNA (nDNA)

The *T. cruzi* epimastigote and the *L. brasiliensis* promastigote forms in exponential growth phase were collected by centrifugation at 3,000 x g for 15 min. The pellets washed twice in TBS (5 x 10^7^ flagellates/mL) were resuspended in the extraction buffer. After a period of 1 h at 37 °C, and the proteinase-K (100 µg/mL) added, the incubation proceeded for 12 h, following the step analogues to those of the host cells.

#### *Trypanosoma cruzi* mitochondrion DNA (kDNA)

The kDNA extracted from *T. cruzi* and from the *L. brasiliensis* by the methods described elsewhere [49]. In brief, either 5 x 10^7^ epimastigotes or equal quantity of promastigotes, respectively, were washed thrice in PBS and collected by centrifugation and resuspended in 630 µL of NET buffer (10 mM Tris-HCL and 100 mM EDTA, 100 mM NaCl, pH 8.0). The cells underwent lysis with the addition of 70 µL of 10% SDS and 7 µL of proteinase-K 20 mg/mL. After incubation for further 12 h the lysate homogenized by sheering, and, thereafter added 690 µL NET buffer with 100 µL 20% sucrose. The mix centrifuge at 14,000 x g for 15 min and the supernatant removed with a pipette tip. The remnant 30 µL was resuspended then in the NET buffer containing sucrose. The pellet collected by the centrifugation and resuspended in 100 µL of distilled water was subjected to subsequent extractions as described for the hosts’ cells. The kDNA precipitated with 2.5 volumes of 100% ethanol and 0.1 volume of sodium acetate 3 M, pH 8.0. The pellet washed twice with ethanol 70%, and the kDNA resuspended in 200 µL of TE buffer and stored at 4 °C.

#### Quantification, enzyme digestion and electrophoretic analysis of DNA

The DNA samples quantified by the spectrophotometric method, and the integrity and purity demonstrated by the analysis in 1% gel electrophoresis run in TAE buffer (Tris-acetate 90 mM pH 8.0, EDTA 25 mM). The digestion of the DNA was carried out by restriction enzymes (Invitrogen); and 2 units of the enzyme cut 1µg of DNA. After 4 h incubation, the DNA fragments were separated by gel electrophoresis. The fragments excised from the gel purified by the DNA Purification kit (Promega) according to the maker instructions.

#### Southern blot

The purified DNA fragments separated by electrophoresis in 1% agarose gel and transferred by the alkaline capillarity method to the Bio-dyne nylon membrane (Invitrogen) in a denaturing solution (NaOH 0.4 M). After the transfer, the DNA dried out and fixed to the membrane at room temperature.

#### Labelling probes

The probes were radio labeled by the Random Primer Labelling System (Invitrogen) by the insertion of [α-^32^P]-dATP in the DNA sequence synthesis by the Klenow polymerase in the presence of the hexamer random primers, in agreement with the maker’s instructions. The radiolabeled probes purified by chromatography in Sephadex G-10 column, and the radioactivity confirmed by scintigraphy showed >10^8^ counts/µg of the DNA. The membrane pre-hybridization caried out for 4 h at 65 °C in the SDS 0.5%, Denhardt solution 5X with 100 µg salmon DNA, according to the Bio-dyne nylon membrane maker’s instructions.

#### PCR amplifications and hybridizations

The *T. cruzi* kDNA sequence amplification carried out with specific primer sets S34/37 and S35/36 [153] that amplifies the kinetoplast minicircle sequence. The minicircle internal oligonucleotides constant region amplification sequence obtained from the *T. cruzi* kDNA, showing primers S34 and S67, and the nested S35 antisense primer from nts 65 to 46. The kinetoplast minicircle constant region kCR probe was PCR-amplified from the *T. cruzi-* infected host cells’ DNA template with primers S34/S67, cloned in the TA vector and sequence. The primer set S34/37 [153] detected as little as 0.15 femtogram of the kDNA minicircle sequence from the *T. cruzi-*infected hosts’ cells. The kDNA and the nDNA specific primers and probes showed in the S1, S2, S4, S6, and S7 Tables.

The *T. cruzi* nuclear DNA primer sequences Tcz1 and Tcz2 [154] amplified the *T. cruzi* nuclear repetitive telomere sequence. The *T. cruzi* nuclear conserved region (nCR), showing 195 nucleotides and the Tcz1/2 primers anneal underlines (S1 and S2 Tables), used probe for hybridization. The PCRs carried out with template DNA 20-fold above the levels of detection, 10 ng of each pair of primers, 0,5 IU of *Taq,* 0.2 mM of each dNTP in a final volume 25 µL. The kCR and the nCR probe radio label [α-^32^P]-dATP showed specific activity 3000 ci/mM and maximum stringency 0.1% SSC and 0.1 SDS, at 65 °C for 1 h.

The amplifications secured with 200 ng of test DNA template and 10 ng of each primer set in the reaction buffer (50 mM KCl, 10 mM Tris-HCL, pH 9.0, and 1.5 mM MgC l_2_), 0.2 mM dNTPs and 2.5 units *Taq-*polymerase. The *T. cruzi* and the *L. brasiliensis* DNA were the controls. The PCR reactions carried out in a PTC-100 MJ Research thermocycler: 94 °C/5min, followed by 32 cycles at 94° C/30 secs. The primer Tm °C/1min, 72 °C/1min, 72°C/7min, and stop at 4 °C. The reactions in triplicates yielded amplification products shown in 1% agarose gel staining by ethidium bromide. The bands in the gels transferred to the nylon membrane and hybridized with the radio labeled specific probes.

#### *tp*TAIL-PCR

A targeting primer thermal asymmetric interlaced-PCR (*tp*TAIL-PCR) [33, 35, 57], using the vertebrate animal LINE-1 specific primers combined with the successive kDNA minicircle internal primers in three cycles of the reaction. The *tp*TAIL-PCR reaction proceeded with different temperature towards the next cycle (S7 Table). A scheme with the *tp*TAIL-pCR to amplify the kDNA integration event into the *Mus musculus* genome showed in the Figure 18.

The *tp*TAIL-PCR was employed to amplification of the *T. cruzi* kDNA minicircle sequence integrated into the host’s genome. The temperature combination with alternate primers and probe (S1, S2, S4, S6 and S7 Tables) specific to the *T. cruzi* kDNA minicircle regions and the oligonucleotides anneal to LINE-1 sequence set forth three straight reamplification cycles (Figure 18). The LINE-1 primer concentration was 10-fold below that of the kDNA primers employed. The primers anneal to a gamut of contigs render amplification of each of the transposon families.

#### The rabbit model: *Trypanosoma cruzi* infection, treatment, and pathology

The two-month-old male and female New Zealand white rabbits purchased from Asa Alimentos (Recanto das Emas, Federal District, Brazil), fed commercial pellet chow and water *ad libitum,* and housed in individual cages at the animal room, at 65% humidity and temperature 24 °C, at the Faculty of Medicine of the University of Brasília. The rabbits of groups A and B were inoculated subcutaneously with 2.5 x 10^6^ *T. cruzi* trypomastigotes collected from the muscle cell cultures.

The therapeutic effect of the nitro-heterocycle Benznidazole searched in the rabbits of the group A. Sixty days after the infection, the *T. cruzi*-infected rabbits of group A treated with intraperitoneal injections of the nitro heterocycle Benznidazole (8 mg/kg body weight) for 60 days. The parasitemia in the *T. cruzi*-infected rabbits’ groups A and B determined by the xenodiagnoses. This consisted of 20 first instar nymphs of the reduviid bug *Dipetalogaster maximus* obtaining blood fills from the A and B rabbits every 15 days for six months. The search for the flagellates in the bugs excreta examined by microscopy at 30- and 60-days post-feedings. The positive exams count derived the curves of parasitemia throughout the course of the infection. The rabbits under clinical inspection weekly subjected to chest X-Rays, and complete necropsy carried out in each rabbit of groups A, B, and C. The tissues fixed in formalin 0.5% were embedded in paraffin, cut 5 µm thin sections in a rotatory micrometer, and H-E staining.

#### Mating the *T. cruzi*-infected rabbits and embryo culture

The sexually mature rabbits mated, and a doe subjected to a Cesarean section under ketamine anesthesia had the fallopian tube flushed with 5 mL of culture medium. The eggs stem cell culture was established from blastocyst harvested 2 days postcoital. The zygotes collected to the wells of a 24-well plate at 5% CO2, 37 °C in a humidified atmosphere. The growing blast cells monolayer bursting the egg adhered to the plastic surface and interacted with the *β-*galactose expressing *T. cruzi* trypomastigotes. In another set of experiments the offspring of a chronically *T. cruzi*-infected dam used to search for the specific antibody and for the genetic markers of Chagas disease. The blood drawn from a wing vein yielded serum and the mononuclear cells, and, also, the solid tissues of the *T. cruzi-*infected and -treated (A), of the *T. cruzi-*infected (B), and of the control (C) rabbits used for the DNA extraction [35], and the serum samples used for the serologic immunofluorescence antibody assay.

#### The chicken model: *Trypanosoma cruzi* inoculation into fertile chicken eggs

The White Ross fertile chicken eggs obtained from Asa Alimentos, Recanto das Emas, Brasília-DF, were inoculated with 100 *T. cruzi* trypomastigotes through a hole made at the eggshell air chamber and sealed with adhesive tape. The mock control eggs received culture medium, only. These eggs hatched after 21-day in chamber incubation at 37 °C, with 65% humidity, while rolling for 1 min every 30 min, in a chicken room maintained at an average 24 °C, under positive pressure filtered air and exhaust. The chicks hatched were kept in incubatory for 24 h and thereafter at 32 ° C for one month. In addition, F0 hens fertilized by artificial insemination laid eggs that hatched the F1 chicks, and the repeat insemination of adult F1 chicken eggs generated F2 chicks. The F0, F1, and F2 chickens used in the experiments [56, 57]. Chicken stem cell cultures were obtained from the blast cells bursting out the zygote membrane six hours after the egg incubation. The germline cells adhered to the plastic surface. The early embryo blast cells growing at 37 °C and 65% humidity were infected with *β-*galactosidase expressing *T. cruzi* trypomastigotes.

The peripheral blood mononuclear cells, semen, and solid tissues from 48 chickens that hatched from eggs inoculated with 100 forms of *T. cruzi,* and from mock control chickens that received 10 µL of culture medium alone. These flocks grew to adults in individual cages, and they fed commercial chow and water *ad libitum*.

#### DNA extraction and nucleic acids analyzes

The semen extracted from roosters, from the embryo tissues of non-inseminated eggs at set points of incubation, from the tissues of the chickens hatched from the *T. cruzi-*inoculated eggs, and from mock controls were processed for DNA extraction. The DNA was extracted from the heart, skeletal muscle, and large bowel of the *T. cruzi* inoculated eggs and from mock control chickens. The genomic DNA were templates for PCR amplification run in triplicates [33]. The amplicons resolved in 1.3% agarose gel, transferred to a Bio-dyne nylon membrane, and hybridized with specific [µ^32^P]-dATP-radio labeled probe. The Random Primer Labelling kit (Invitrogen) employed to label the nCR and kCR probes [33, 36].

The Southern hybridization of DNA from the *T. cruzi*-infected, from mock control eggs, and from the *T. cruzi* kDNA positive control performed in agreement with the protocols described elsewhere [33, 35]. The *tp*TAIL-PCR, cloning and sequence carried out with the chicken primers Gg1 to Gg6 (S4 Table) anneal to a specific locus of the chimera sequence (GB FN599618) in the *locus* NW_001471673.1 on chromosome 3 [10. 68]. The primers 0.004 µM dilution and their annealing temperature used in combination with the kDNA primers [66]. The clones revealed by hybridization with the radio labeled kDNA and sequenced commercially. The *tp*TAIL-PCR validated in a mix of *T. cruzi* kDNA with DNA from naïve control birds.

### The mouse models

#### *Trypanosoma cruzi* infection, parasitemia, and pathology

The two-month-old BALB/c mouse weighing 20 to 25 grams were used. The mice were inoculated intraperitoneal with 10 x 10^3^ *T. cruzi* trypomastigotes harvested from L6 muscle cell cultures and suspended in DMEM. The groups of 8 male and female mice were housed, separately, in cages maintained at 24 °C, 65% humidity, under pressure air exhaust, and fed commercial pellets and water *ad libitum*. The parasitemia in the *T. cruzi-*infected groups of mice determined by direct microscopic exam of the tail blood obtained at set points throughout the course of the infections. In addition, the demonstration of the *T. cruzi* in the blood was made by the hemoculture in screwcap glass culture tube containing 5 mL of blood-agar slant plus 5 mL of LIT medium overlayer. These culture tubes inoculated with 100 µL of the blood drawn by the heart puncture from the *T. cruzi-*infected mice, were kept in a shaker incubator at 27 °C, during a period of 60 days. The cultures examined by microscopy at 20, 40 and 60 day-set points. The A, B, and C groups of *T. cruzi-*infected mice that died during the *T. cruzi* infections or by accident during heart puncture, and tissue samples removed. At the end of the experiments the mice that were sacrificed under anesthesia had the body tissue samples subjected to DNA extraction, and samples fixed in formalin used in the histopathology study. These procedures employed to search for tissue lesions in the groups of *T. cruzi-*infected mice treated with the drug inhibitors of eukaryotic cells growth and differentiation.

#### Multi-drug treatment of the *T. cruzi-*infected mice

Groups of eight, two-month-old BALB/c mice in cages separated for the multidrug treatment of the *T. cruzi* infection. Each mouse in the groups A to G received intraperitoneal inoculation of 10^4^ *T. cruzi* trypomastigotes derived from L6 muscle cell cultures: A) *T. cruzi-*infected-positive control, untreated. Each *T. cruzi-*infected mice in the groups B to G received 0.86 mg Benznidazole daily (See Methods). In addition to Benznidazole, each mouse of group received AZT (1.2 mg), whereas group D received Ofloxacin (1.2 mg). Also, a multidrug regime with the above drug concentration administered to mice groups: E) Benznidazole and AZT. F) Benznidazole and Ofloxacin. G) Benznidazole, AZT, and Ofloxacin. The group H) negative control, uninfected. The drug daily administration regime initiated at the fifth day post-infection and continued for 60 days. Each drug powder stock suspended in distilled water to the concentration indicated in the Table 1 and the aliquot mix administered through gavage using a 4 cm long urethral catheter n° 3.

The parasitemia in the *T. cruzi-*infected mice determined by direct search of the flagellates in a drop of the tail blood at the 5^th^ day post-infection and thereafter each 15 days set points until the 60^th^. The parasite count in 5 µL of blood under 2 x 2 coverslip on glass slide achieved by microscopy. Also, de presence of the late phase parasites determined by hemoculture of 100 µL of blood collected through heart puncture at 20-, 40- and 60-days post-infection. The mortality in the A-to-H groups of mice recorded, and the deceased subjected to necropsy had the tissue samples fixed in formalin and subjected to histopathology. The mice survivals sacrificed under anesthesia at the end of the experiments at the 120^th^ day.

#### ELISA and nucleic acids assays

In the absence of directly demonstrable blood parasitemia, the presence of the *T. cruzi* antibody revealed by the enzyme-linked immunosorbent assay (ELISA). The soluble antigen obtained by three cycles of freezing and thawing the *T. cruzi* epimastigote forms harvested from the LIT medium and washed thrice in PBS, pH 7.4. The antigen suspension (0.2 µg/100 µL) used to sensitize each of 96 well in carbonate buffer, pH 9.6, at 4 °C for 12 h. After three washings with PBS-*Tween* 0.05% the wells received an equal volume of 2% defatted milk to block unspecific antigenic sites. Each triplicate wells thrice washed with PBS-*Tween* received 100 L of the 1/100 serum dilution and the plates set to incubate for 2 h at room temperature. After the serum dilutions were discarded and the plates subjected to further washings with PBS-tween, and equal volume of the peroxidase-labeled goat anti-mouse IgG 1/1000 dilution added to the wells and incubated for 2 h. The reaction proceeded in the citrate buffer pH 5 containing H_2_0_2_ at 30 and 20 µg of the OPD substrate. The development of color yellow interrupted by addition of 4N sulfuric acid. The plate reading made in a Synergy HT spectrophotometer (Bio Systems) at wavelength 480 nm. The optical density above 0.064 cut off the positive from the negative antibody reaction.

#### Extraction of DNA from the *Trypanosoma cruzi-*infected, infected-treated, and from control uninfected mice

The mice survivals were sacrificed under ketamine anesthesia at the 120-day-end experiment. Sterile blade used for each mouse and the tissues obtained from the heart, skeletal muscle, large intestine, and spleen. Small fragments (circa 200 mg) of each tissue transferred to sterile Petri dish and minced. The tissue mixes from each mouse transferred to sterile 15 ml screw-cap plastic tubes and incubated with 1 µg/mL proteinase K (Gibco) for 12 h at 37 °C. The tissues subjected then to one extraction with chlorophane (phenol: chloroform: isoamyl alcohol, 25: 24: 1) and a further extraction with chlorophyl (chloroform: isoamyl, 24:1). The sample mixes agitated by inversion of the tubes and centrifuged at 4,500 x g for 10 min at room temperature. The aqueous supernatant collected, and the DNA precipitated with absolute ethanol (1:10 v/v) at – 8 °C and washed twice with 70% ethanol at 20,800 x g for 15 min at 4° C. After washings and evaporation of ethylic alcohol the DNA resuspended in buffer TE (Tris-HCl 10 mM, EDTA 1 mM, pH 8.0) containing RNase, 200 ug/mL (Invitrogen). The DNA quantified by 1% agarose gel electrophoresis and ethidium bromide staining. The detailed PCR protocol for the nucleic acids’ diagnosis of the *T. cruzi* infection with the tidy methods to identifying the *T. cruzi* nuclear DNA (nDNA) in the tissues is described elsewhere [33, 35, 57].

#### Targeting primer Thermal Asymmetric Interlaced PCR (*tp*TAIL-PCR)

The *tp*TAIL-PCR combine the kDNA primers with the primer sets specific to the LINEs (S6 and S7 Tables). The reactions run under set temperature changes for the anneal of the oligonucleotides along three reaction cycles (Figure 18). The DNA samples amplified after each reaction cycle initiated by the kDNA minicircle sequence primer combination with the mouse LINE retrotransposon specific primer. The *tp*TAIL-PCR cycles proceeded in High Fidelity buffer (Invitrogen) containing 2 mM MgCL_2_, 0.4 µL primers (IDT Technologies) and 200 ng DNA in the Bio-Rad My Cycler^TM^ thermocycler. The transposon LINE primers and the *T. cruzi* kDNA probe sets shown on the S1, S2, S4, and S6 Tables.

#### Southern blot and radio label probe hybridization

The *tp*TAIL-PCR amplified sequences separated by electrophoresis in 1% agarose gel and transferred to positively charged nylon membrane (HybondTM-N+, AmershamGE Healthcare) by the alkaline transfer method [33, 35]. The *T. cruzi* kDNA kCR subjected to [α-^32^P] -dATP (PerkinElmer). The Random Primers Labelling System (Invitrogen) used in the reaction with the triphosphate deoxynucleotides’ dATP, dCTP, dGTP and dTTP (20 µM), three units of the Klenow DNA-polymerase-I, 15.0 µL random primer buffer mixture, and 5 µL radio labeled dATP at 3,000 µCi activity. The reaction took place for 3 h and the radio labeled kCR probe purified in a Sephadex G25 column (Amersham, GE Healthcare).

The DNA immobilized on the nylon membrane had positive charges blocked with the pre-hybridization solution (PEG 800 10%, SDS 7%, SSPE 1.5%, salmon sperm, 100 µg/mL) at 65 °C for 15 min. The membranes then inserted into cassette (Kodak Biomax) for film exposure to the X-rays (Kodak T-Mat) and stored at –80 °C for several days. The films washed with the developing and fixation solutions (Kodak) in the dark to show the DNA bands.

#### Cloning and sequencing

The DNA sequences obtained after the *tp*TAIL-PCR third amplification cycle were cloned. Each mouse DNA sample subdivided in eight aliquots and the amplification from each *tp*TAIL-PCR third cycle products carried out four repeat runs. The sequences ligated by the T4 ligase at 4 °C for 12 h were cloned in the pGEM T-easy vector (Promega). The competent *Escherichia coli* X10-Gold (Stratagene) treated with rubidium chloride and transformed by thermal shock (Promega Protocols and Applications Guide). The cells inoculated in LB broth (USP Corp) at 37 °C for 3 h to the maximal growth achieve at OD 600. The broth then centrifuged, and the cells transferred to Petri dish to grow in LB Luria Betani agar (HIMEDIA Labs) with X-Gal at 4.8 x 10^-2^ µg/mL and ampicillin, 0.1 µg/µL.

The recombinant selection of the ampicillin resistant white colonies, whose cells were placed to grow on nylon membranes treated as for the hybridization with the radio labeled kCR probe. The recombinants showing intense hybridization signal selected for the plasmid extraction using the IllustraTM plasmidPrep Mini Spin Kit (GE Healthcare). The inserts released by the *Eco*RI (Invitrogen) enzyme digestion, showing strong hybridization signals were commercially sequenced (Molecular Genomic, São Paulo). The BygDye^(R)^ Terminator v3.1 Cycle Sequencing Kit employed to sequence the products in a 3130xl Genetic Analyzer (Applied Biosystems). Four rounds of the *tp*TAIL-PCR third cycle amplification yielded selection of the non-redundant sequences from the DNA of each of 5 groups of mice (Figure 19).

#### Sequence analyzes

The sequences subjected to the analyzes of similarity with those from *Mus musculus* and from *Trypanosoma cruzi* retrieved from the GenBank/NCBI. The non-redundant (nr) searches made throughout the GenBank/EMBL/DDBJ and RefSeqs data bases. The Blastn version 2.2.25 parameters used: e-threshold 10; word size 11. The scores secured: Match/Mismatch 1, -2; gap costs, existence 5 and extension 2. The repeat masker CENSOR used to map the sequences, and the repeats classification obtained by alignment with those in the LINEs families A, T_F_, and G_F_. The Table 2 show the reference sequences L1Md_-_A2 (M13002.1) for family A, L1spa (AF016099.1) for family T_F_, and L1Md_-_GF62 (MG1:2178803) for the G_F_.

#### Statistics

The One-way ANOVA-F employed for the statistical analysis and evaluation of distributions of the kDNA mutations among the experimental data.

#### Histopathology

The tissues from the mice of groups A to H were washed with PBS 1X, pH 7.2, and fixed in 4% formaldehyde solution for 72 h. The samples were washed in distilled water and dehydrated in ethanol for 30 min at 70%, 80% and (90%) concentrations. The samples then washed thrice in 100% ethanol were transferred to a solution of absolute ethanol/xylene (1:2) for 15 min. After three washings in xylene the samples were immersed in paraffin at 60 °C for 1 min and then twice for 30 min. The paraffin blocks transferred to room temperature and the blocks sections cut 4 mm thick in a rotatory microtome (Leica RM2135) and mounted in glass slides. The paraffin removed by immersion in xylene for 5 min and rehydrated in gradient ethanol dilutions from 100% to 70%, for 2 min. The slides dried out at room temperature and subjected to Hematoxylin-Eosin staining.

## Ethics

The Human and the Animal Research Committees of the Faculty of Medicine of the University of Brasilia approved all the procedures with human subjects and laboratory animals research protocols 2500.167567 and 054/09, respectively. The laboratory animals received humane care; the mice under anesthesia subjected to heart puncture before sacrifice.

## Abbreviations

AZT: Azidothymidine
CSB: constant sequence block
DMEM: Dulbecco minimal essential medium
kDNA: *T. cruzi* mitochondrion kinetoplast DNA
kCR: kDNA constant region internal probe
LINE-1: long interspersed nuclear element-1
LIT: liver infusion-tryptose nutrient broth
mya: millions of years ago
nDNA: *T. cruzi* nuclear DNA
nCR: nDNA telomere constant region probe
SINE: short interspersed nuclear element
[α-^32^P]-dATP: ^32^P-(32) p_-_labeled; 2-deoxyadenosine triphosphate

## Data Availability

All relevant data are within the manuscript and its Supporting Information files. Further core data information found at https://repositorio.unb.br

## ACKNOWLEDGMENTS

We acknowledge José Tavares for the animal care, and Rafael Andrade for the surveillance of the animal rooms and delivery of humane care.

## SUPPORTING INFORMATION

**S1 Figure. Chemical inhibitors of the cell growth and differentiation.** The figure depicted the chemical structure and theoretical formula of the range of inhibitors of eukaryote cells growth and differentiation showed in the Table 1.

**S2 Figure. *Trypanosoma cruzi* minicircle sequences repeats end-joining homologous recombination into the mouse LINE-1 DNA.** The CSB1, CSB2 and CSB3 consensus regions shared by the kDNA minicircle and the mouse LINE-1 and SINE. The blue sequences indicated the *T. cruzi* kDNA and the green showed the mouse DNA. The consensus regions appear underline. The AC/TG-reach repeats are common features intermediate the end-joining homologous recombination of the Protist DNA with the mammalian DNA.

**S1 Table. The *Trypanosoma cruzi* primer used in the PCR amplification of the mitochondrion kDNA minicircle sequence.** The primers S35, S36, and S37 specific anneal to the kDNA minicircle constant sequence block depicted in the S2 Fig [152]. The highly sensitive nDNA telomere primer set specific anneal detected 15 femtogram of the *T. cruzi* nDNA used in the nucleic acids assay to the accurate diagnosis of the protozoan infection [153].

**S2 Table. The Trypanosoma cruzi nucleus and mitochondrion DNA derived probe sequences with primer underlines.** The specific nDNA telomere (nCR) sequence and the kDNA conserved sequence fragment (kCR) used in the *tp*TAIL-PCR amplification to show the *T. cruzi-mouse DNA* hybrid sequence that indicated the lateral transfer of the Protist kDNA into the mammalian genome.

**S3 Table. Mapping of the sites of *Trypanosoma* kDNA integration into the chromosomes of the rabbit.** The 5’Race PCR products were reamplified and directly cloned in pGEM T easy vector. The clones confirmed by hybridization with the radio labeled kCR probe were sequenced commercially. The amplifications revealed the hybrid sequences with the kDNA primer sets at both extremities of the rabbit retrotransposon LBNL-1 intermediate sequence.

**S4 Table. *Gallus gallus* primer sets used in the tpTAIL-PCR amplification of *T. cruzi* mitochondrion kDNA-chicken’s transposon hybrid sequence**. The primer sets were used in combination with the S35/36 kDNA primers in three cycles of reactions, accordingly, with the annealing temperature indicated in the S6 Table. The amplification products hybridized with the radio labeled kCR probe revealed the lateral transfer of the kDNA minicircle sequence into the chicken transposable elements.

**S5 Table. Mapping the sites of *Trypanosoma cruzi* kDNA integration into the chromosomes of the chicken**. The *tp*TAIL-PCR employed the chicken specific primer sets in combination with the kDNA specific primer sets (S3 Table). The amplification products that hybridized with the kCR probe were cloned and sequence. The sequence analysis showed the AC-reach repeats (Fig S2) intermediate the integration of the *T. cruzi* mitochondrion sequence into the chicken genome, and that 72.8% of the integrations took place in the long transposable chicken CR-1 repeats.

**Table S5A.**
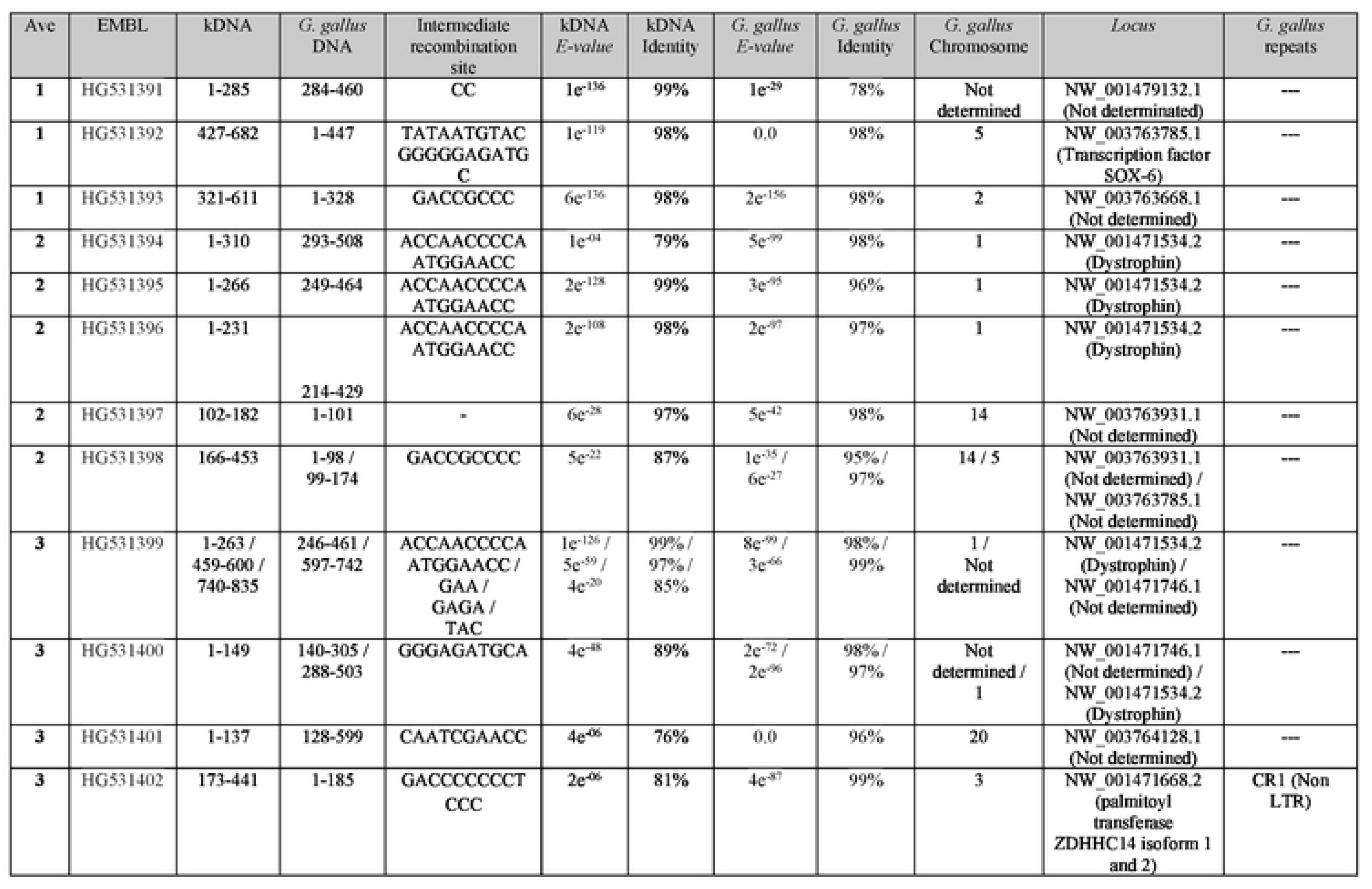
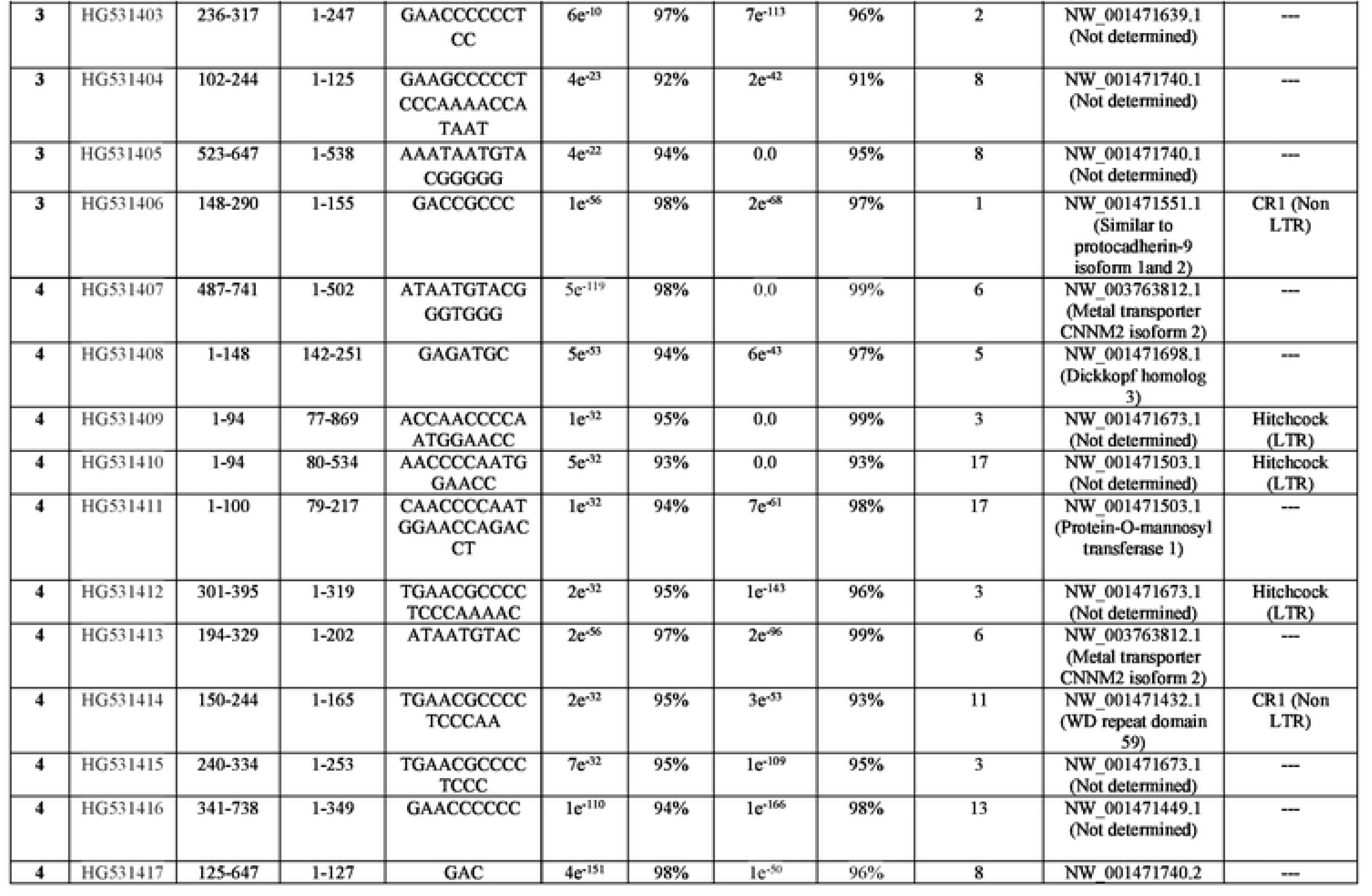
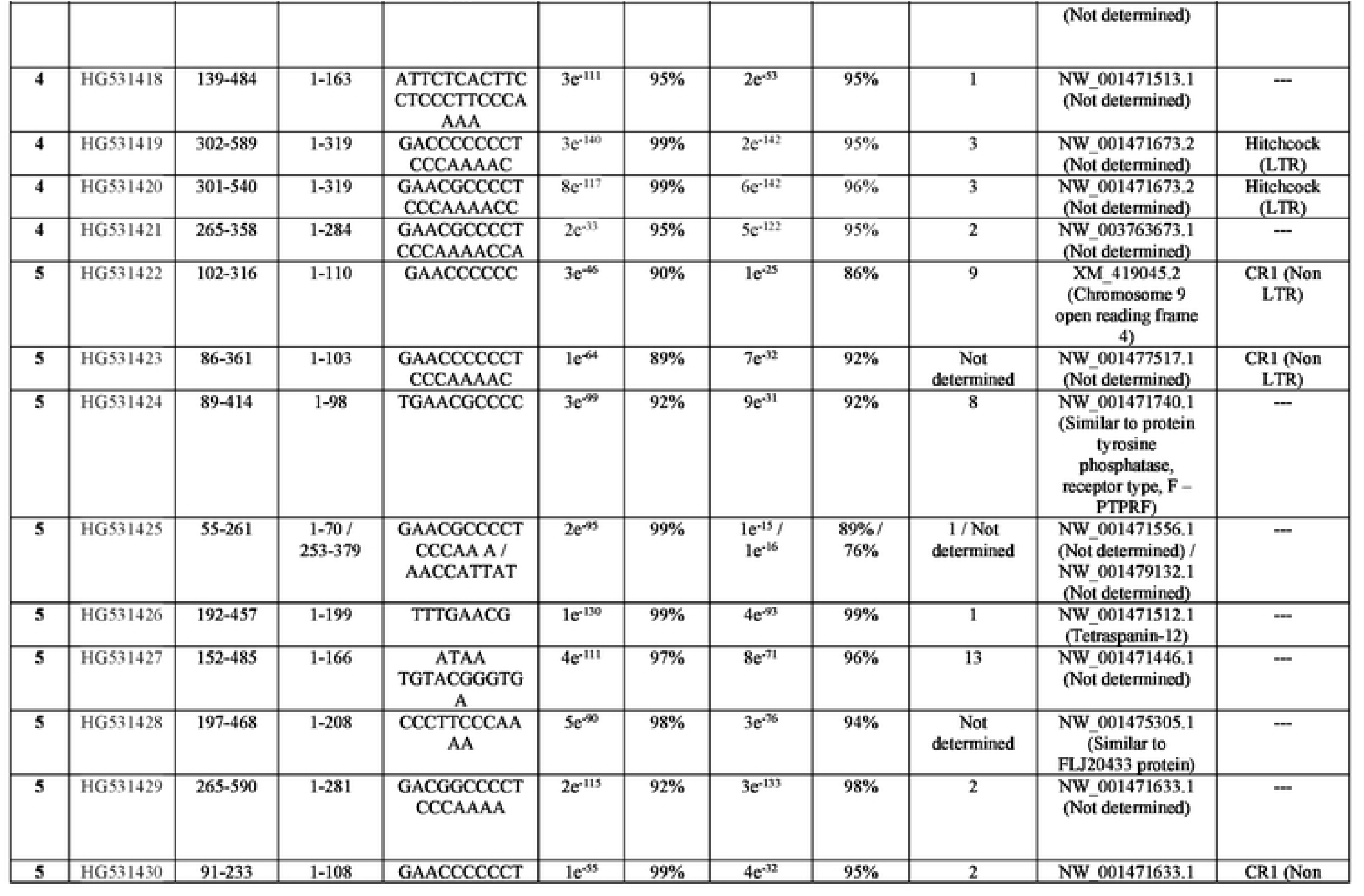
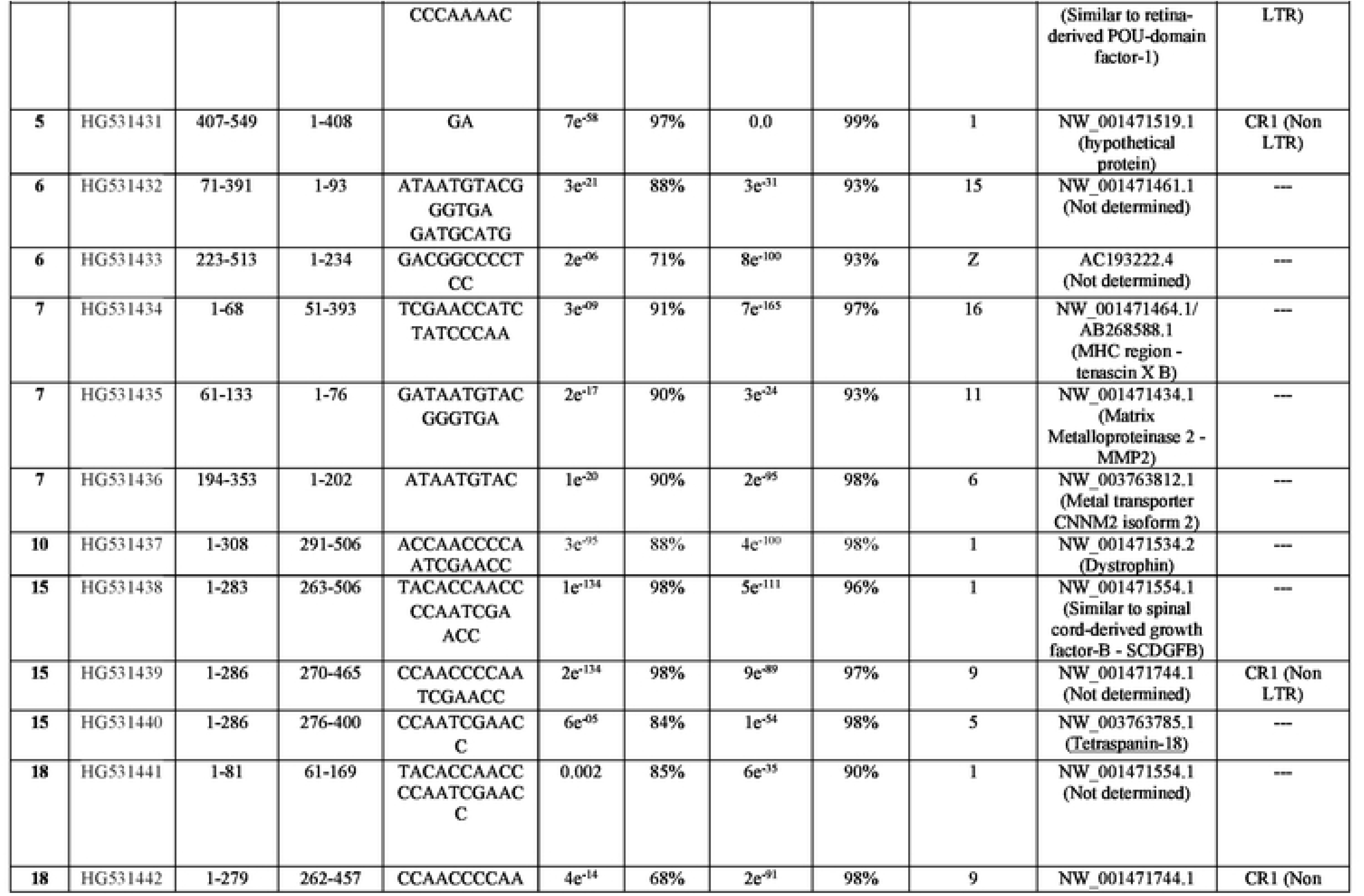
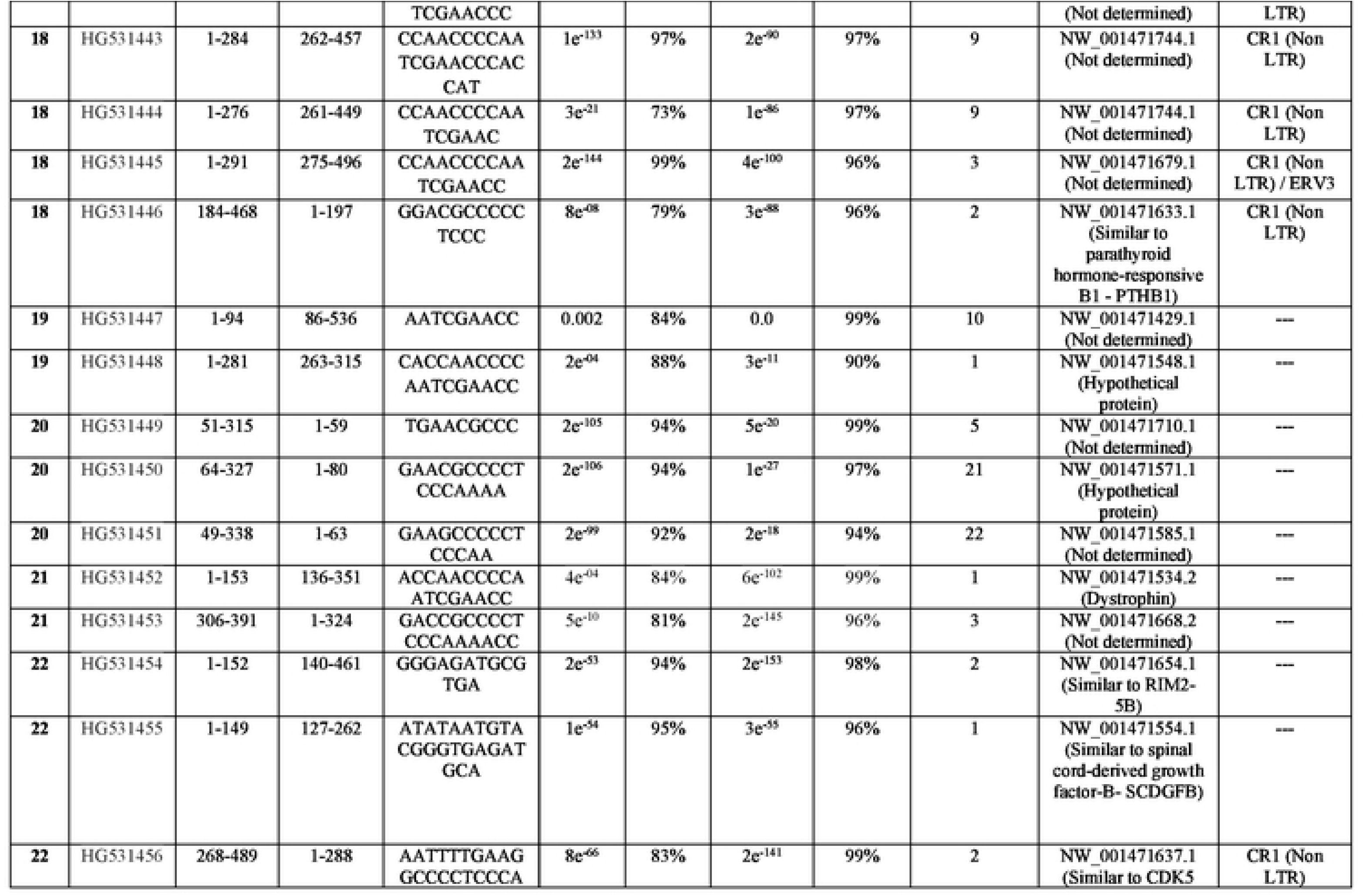
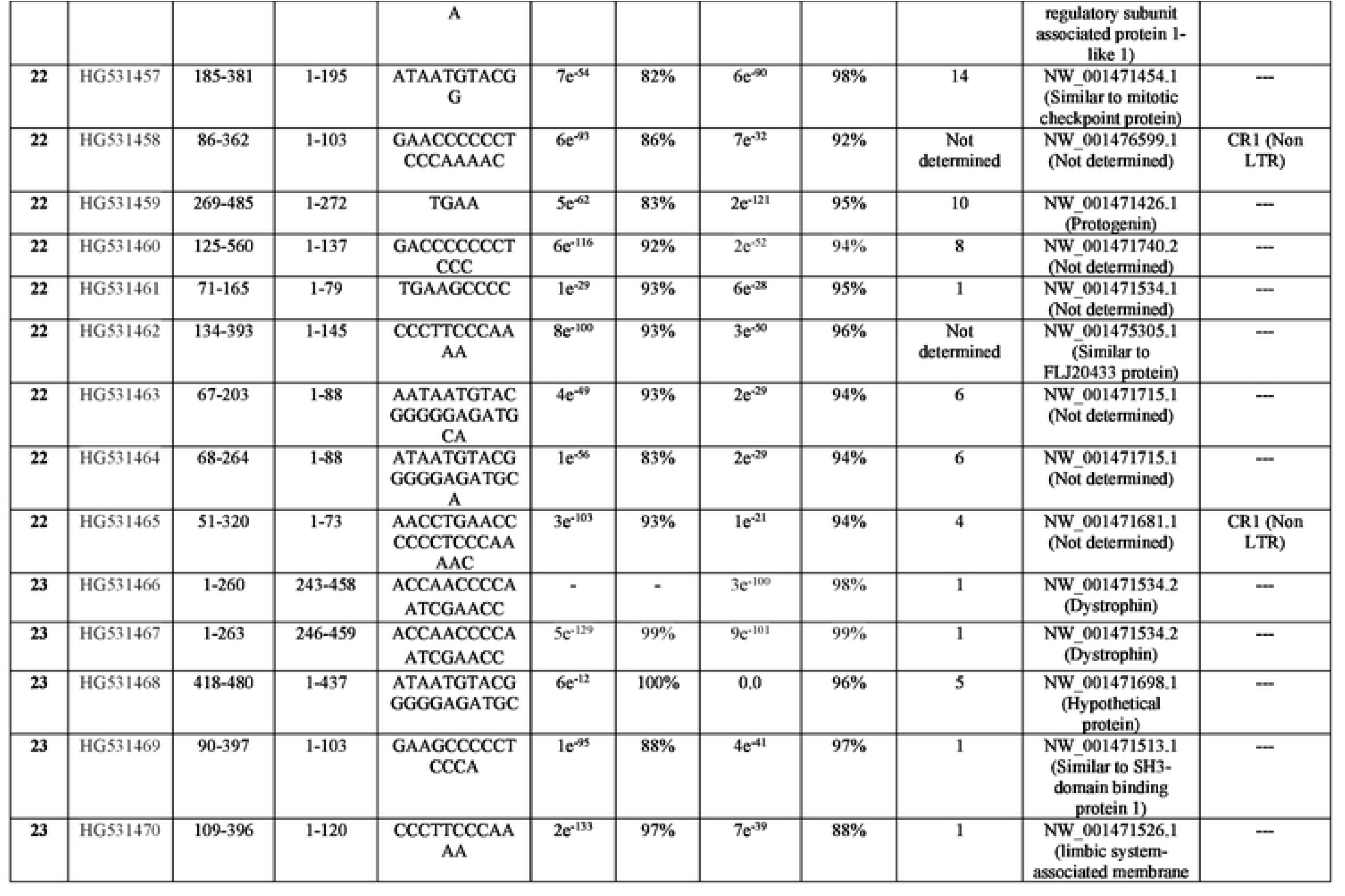
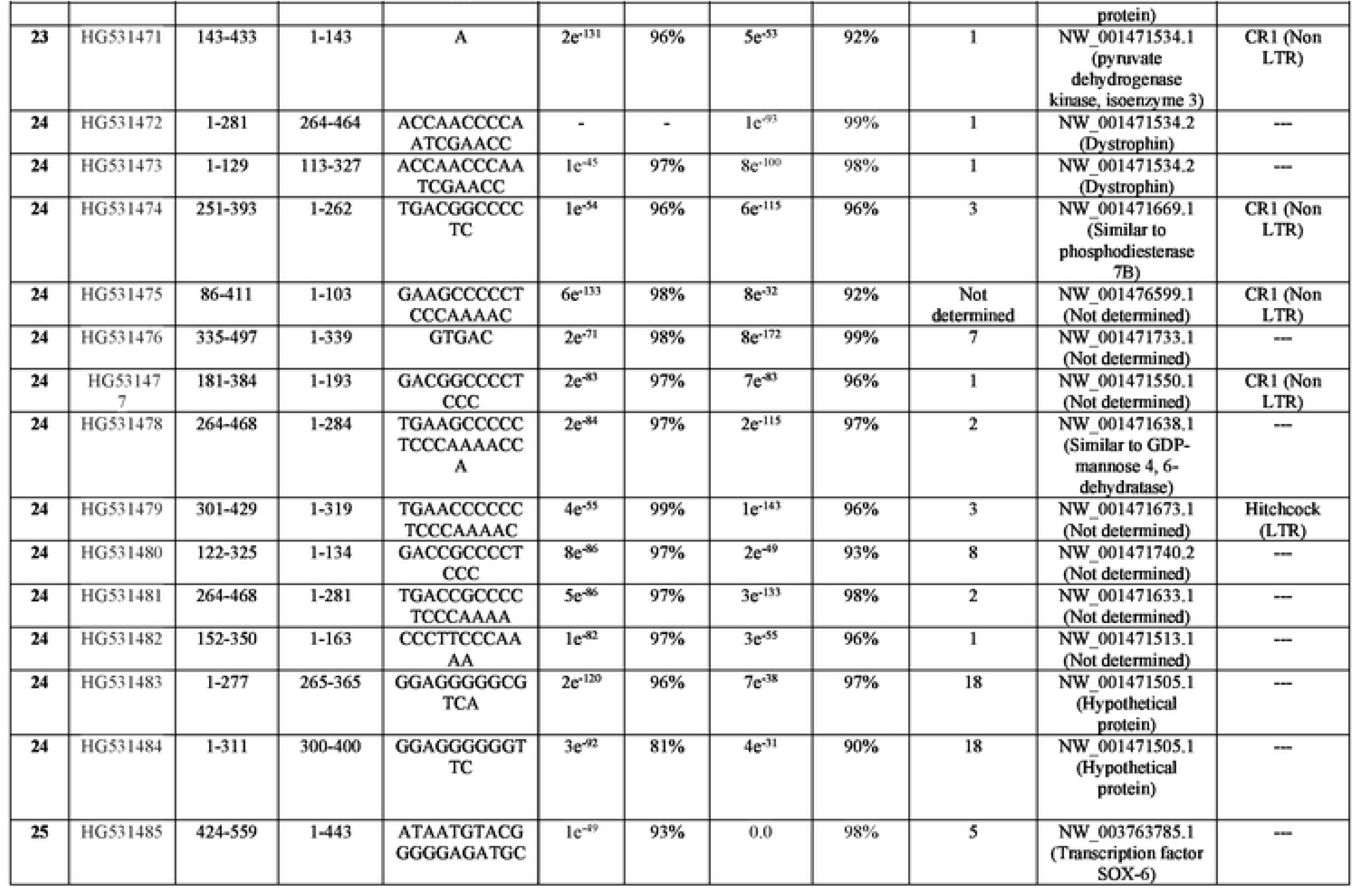
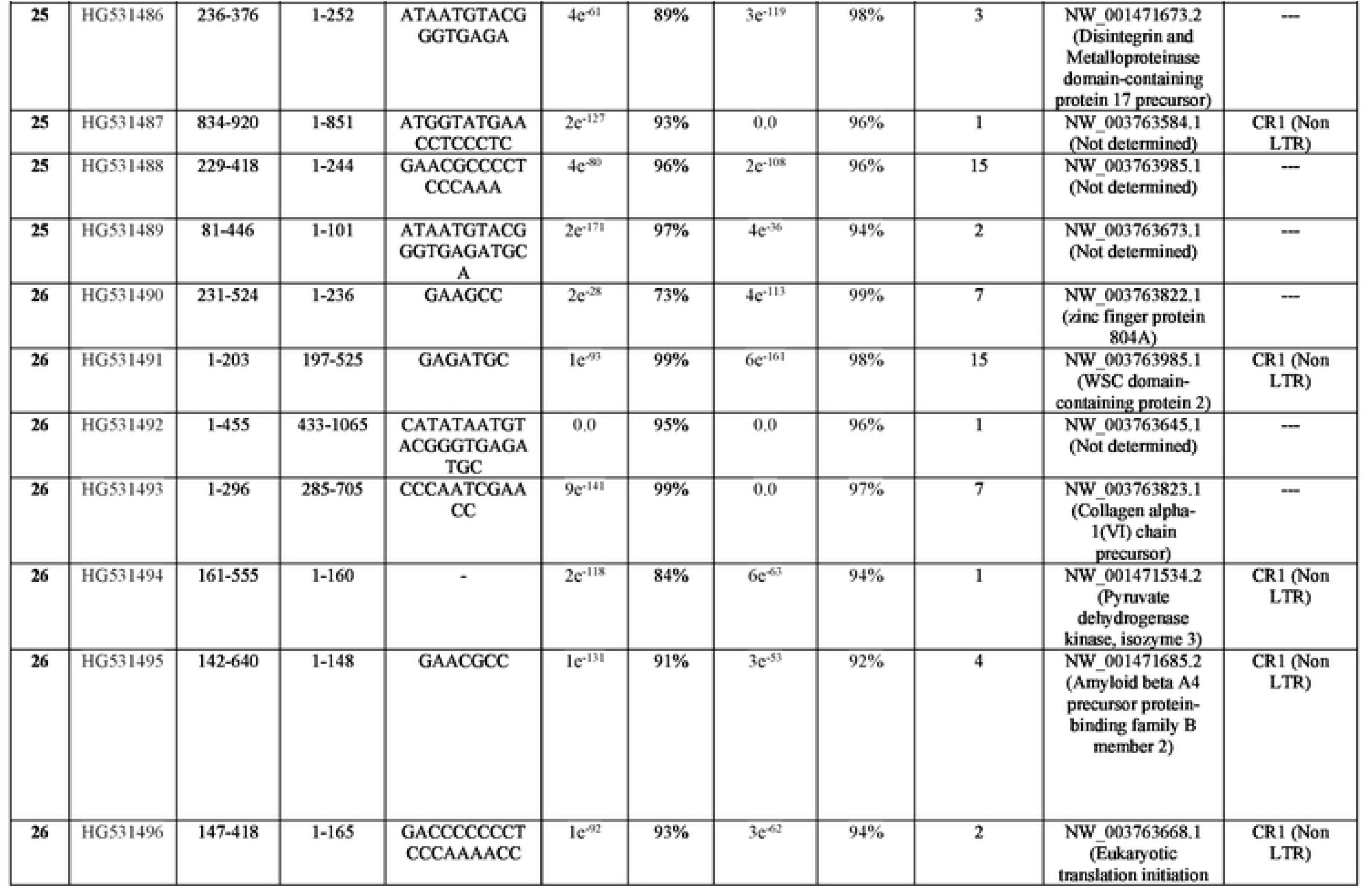
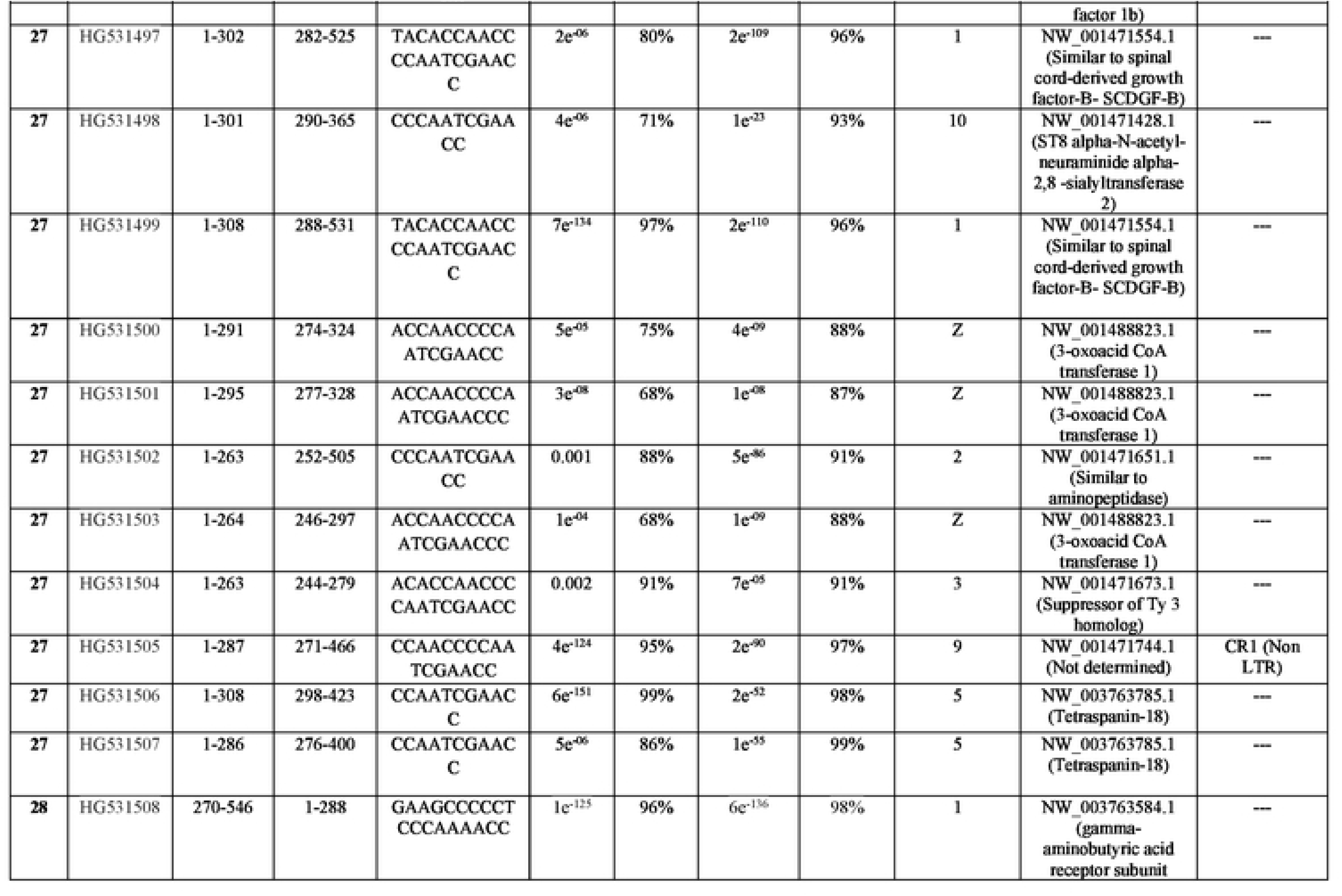
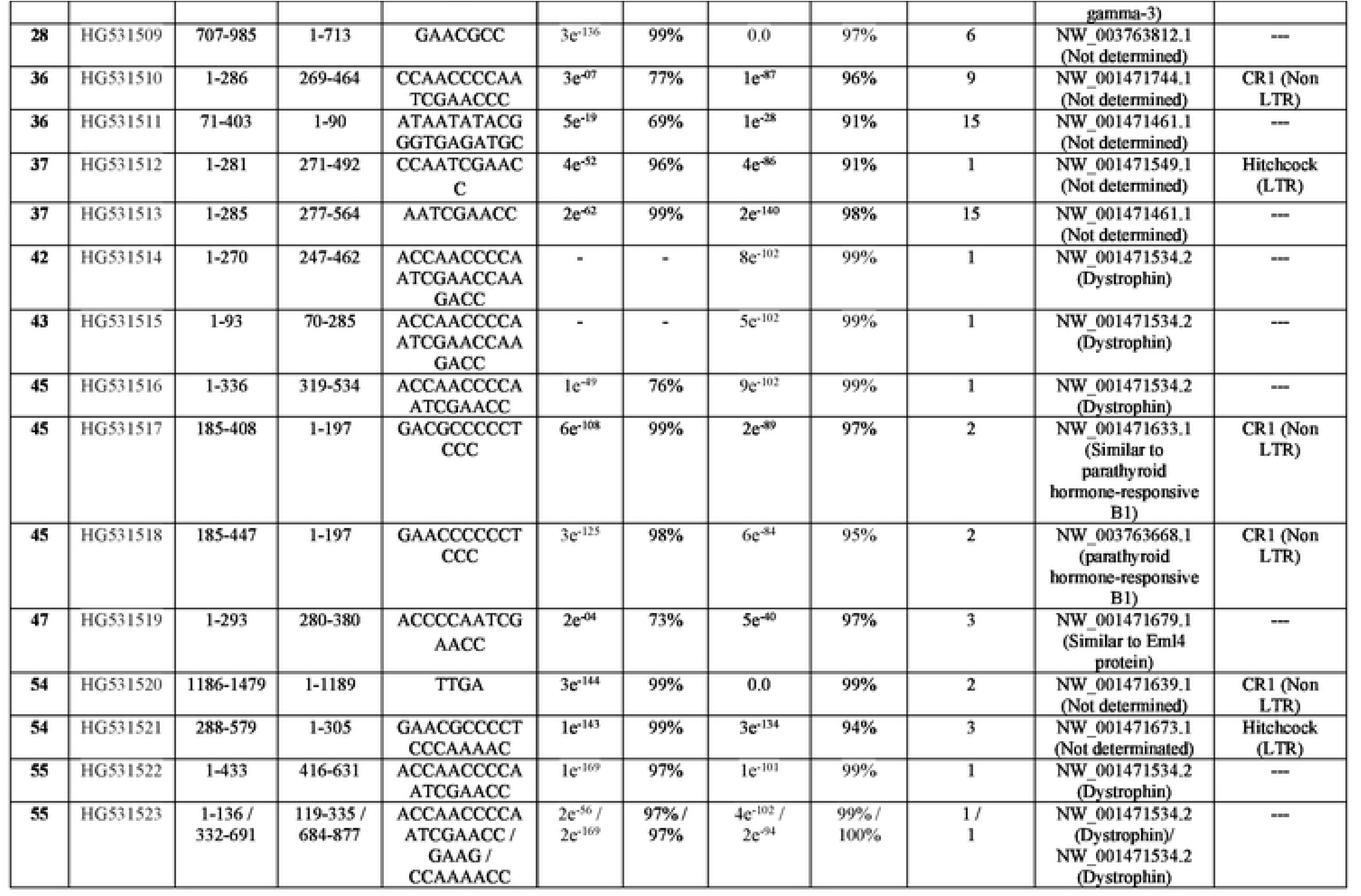
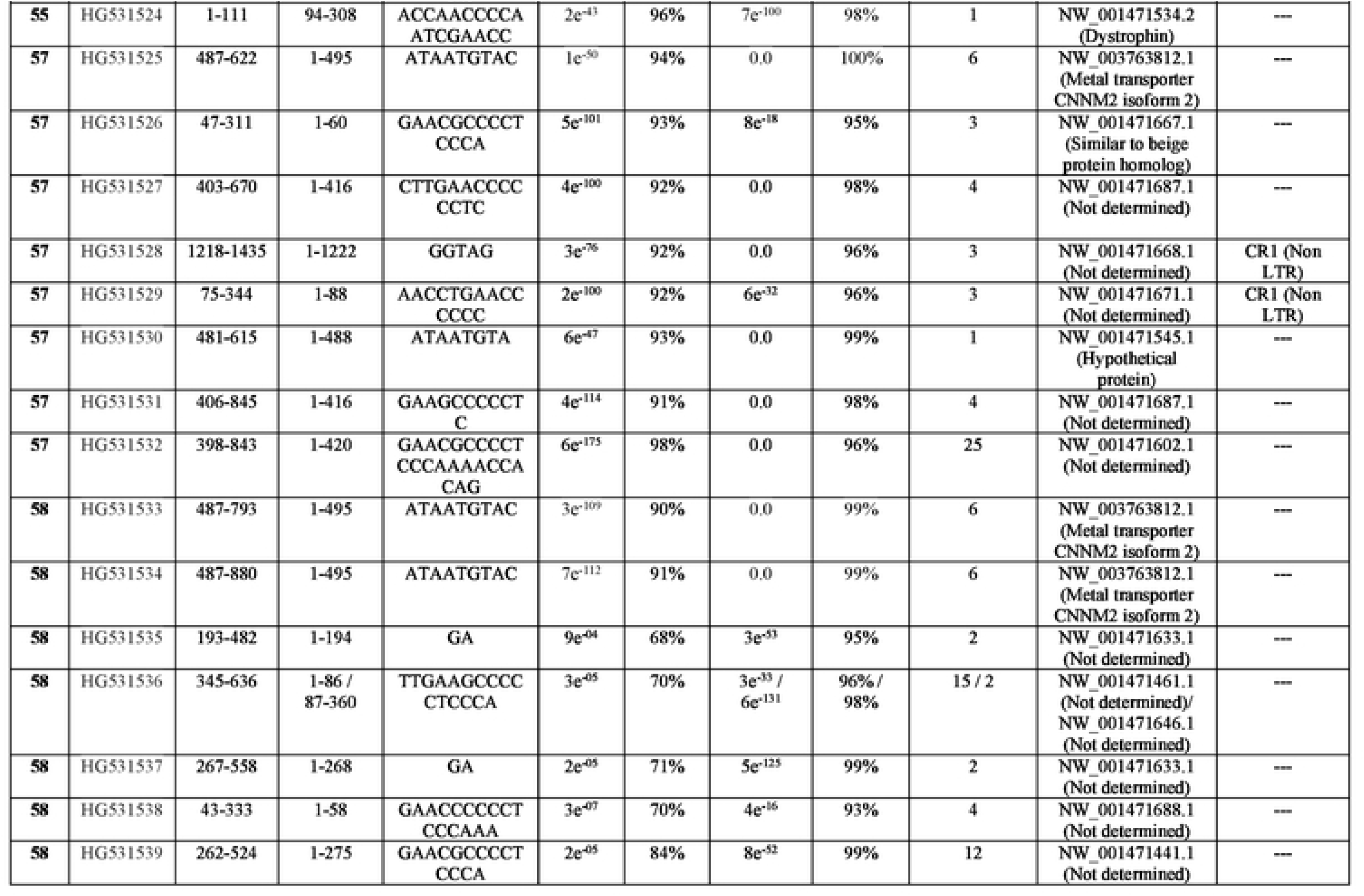
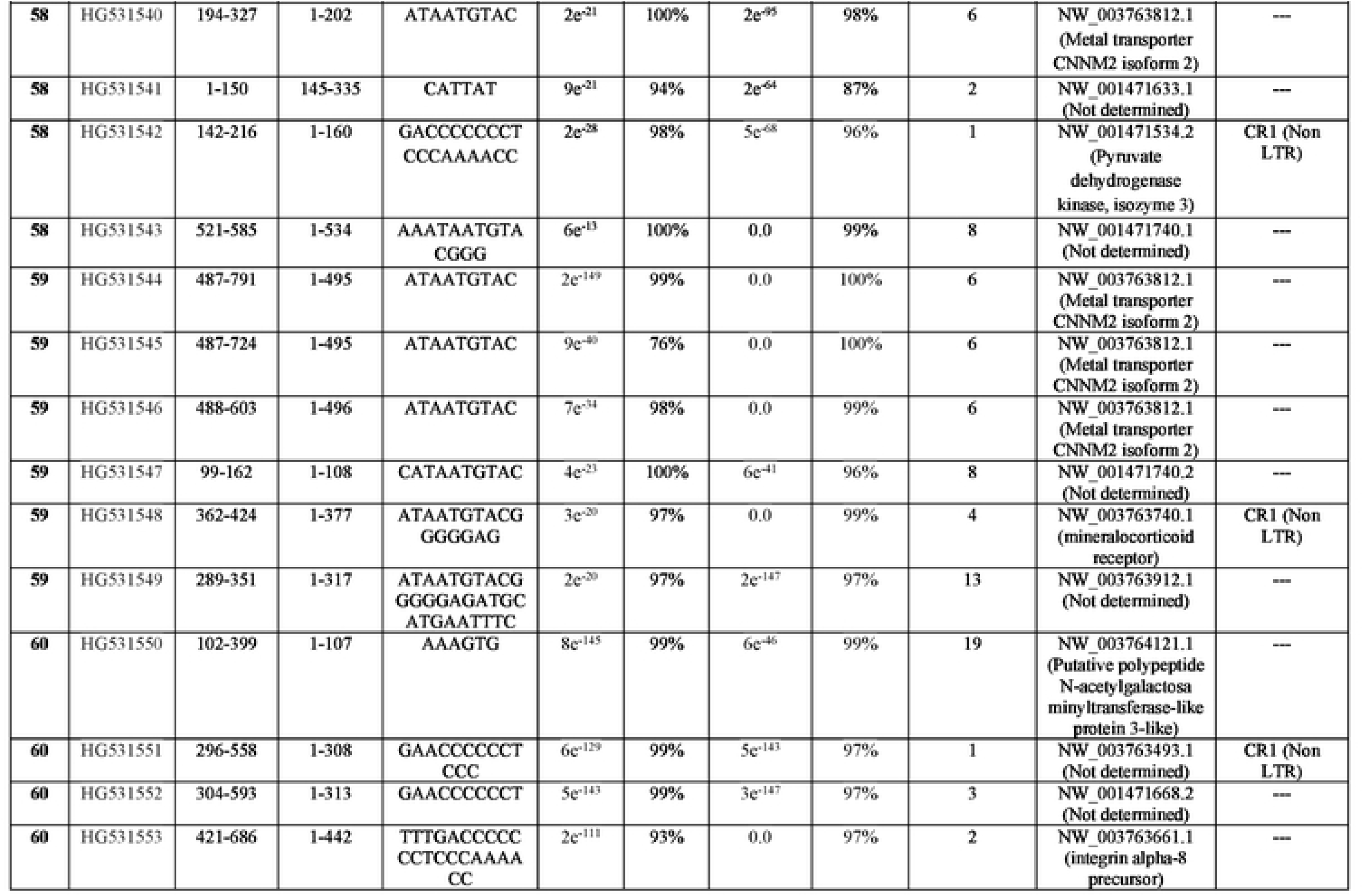
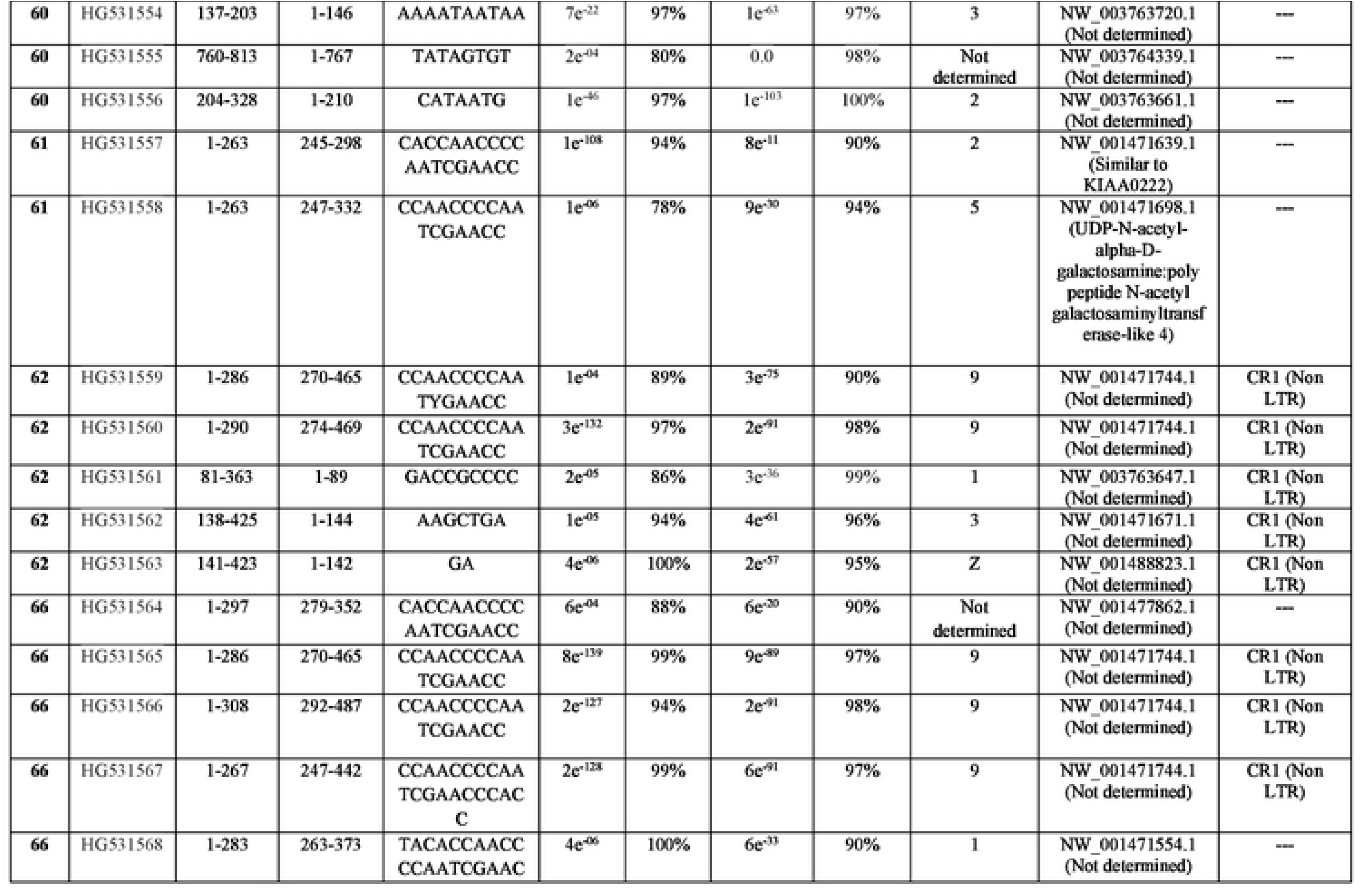
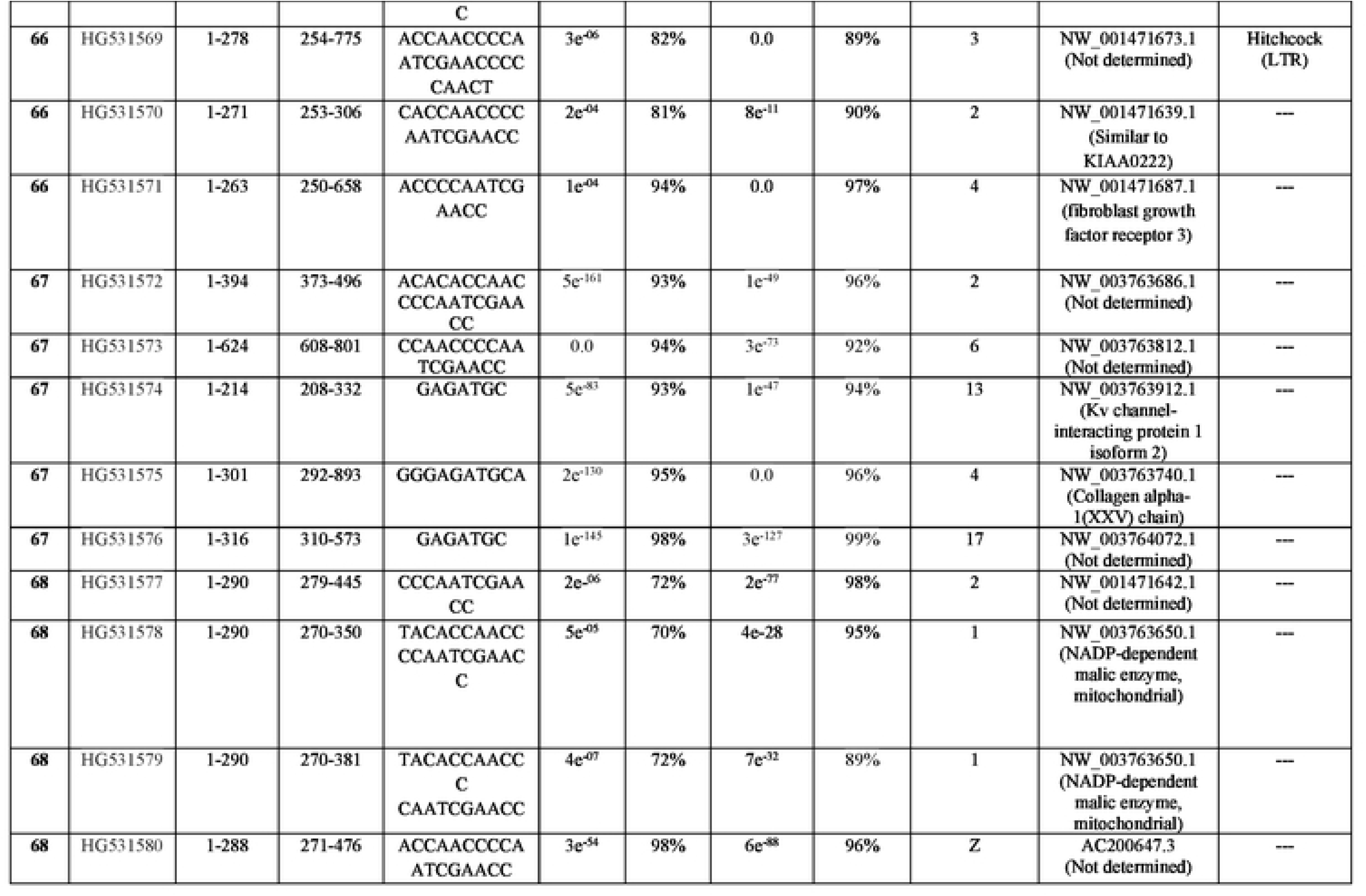
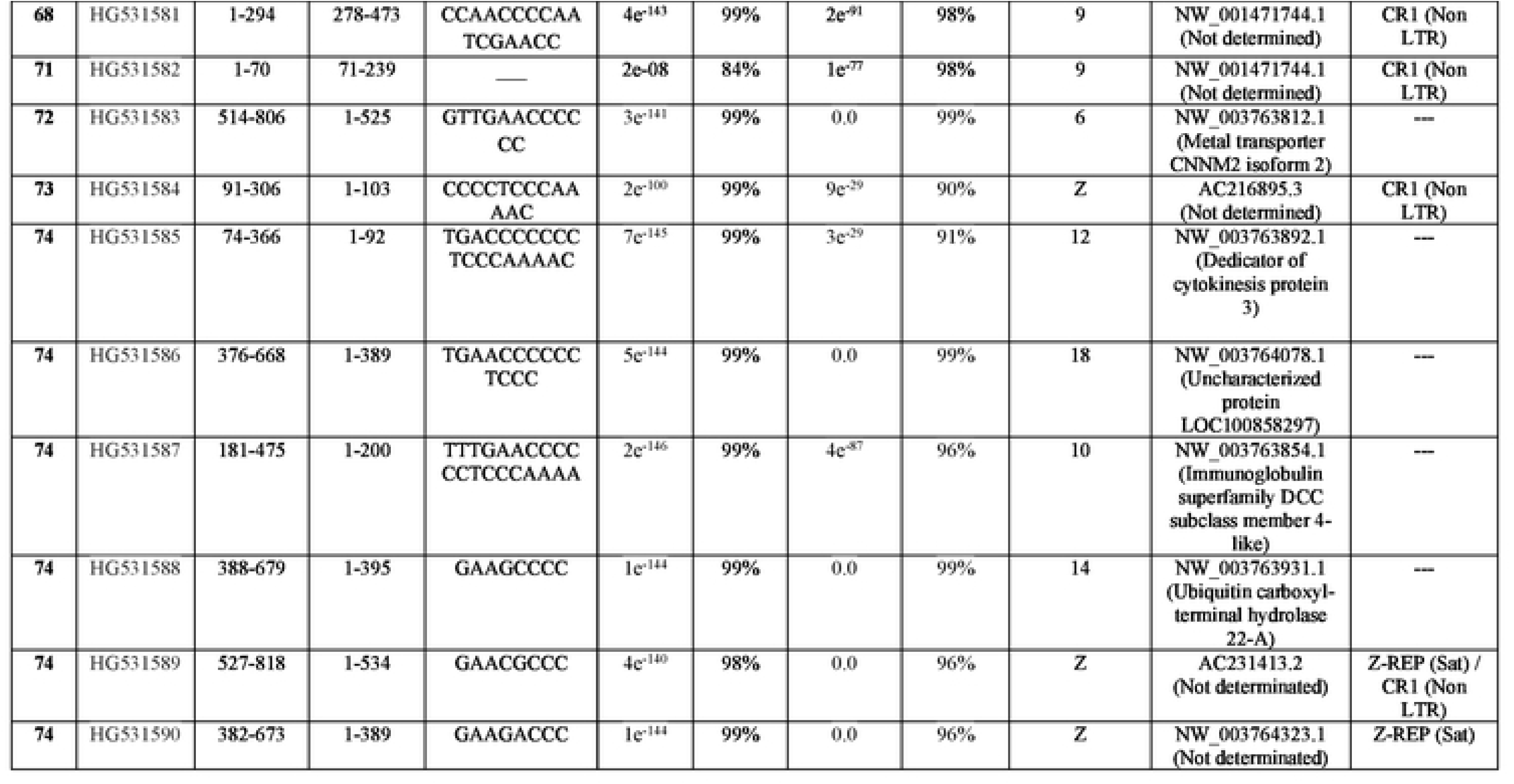
Lateral transfer of minicircle sequences of kDNA from *Trypanosoma cruzi* to the genome of *Gallus gallus* somatic cells.

**Table S5B.**
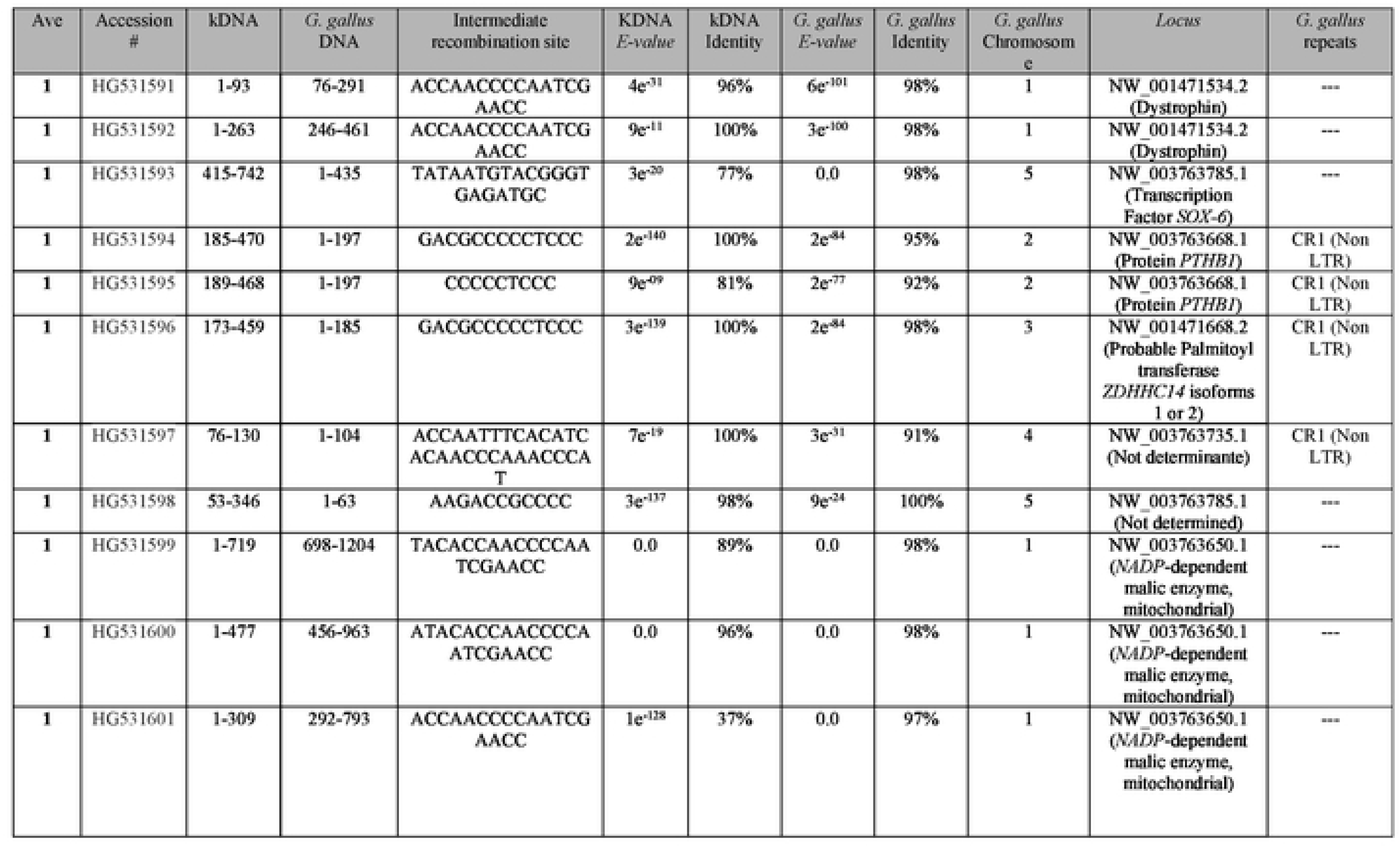
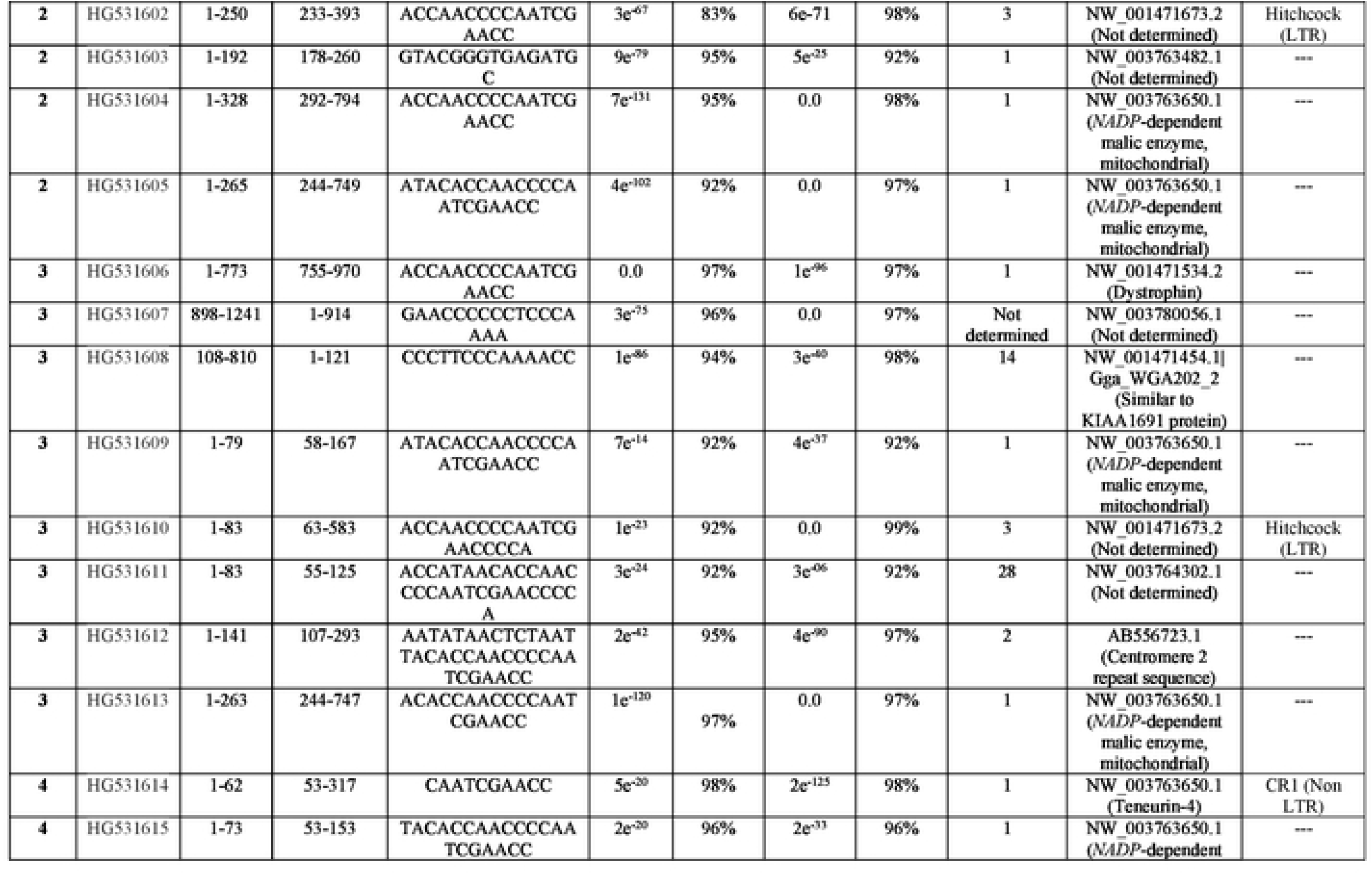
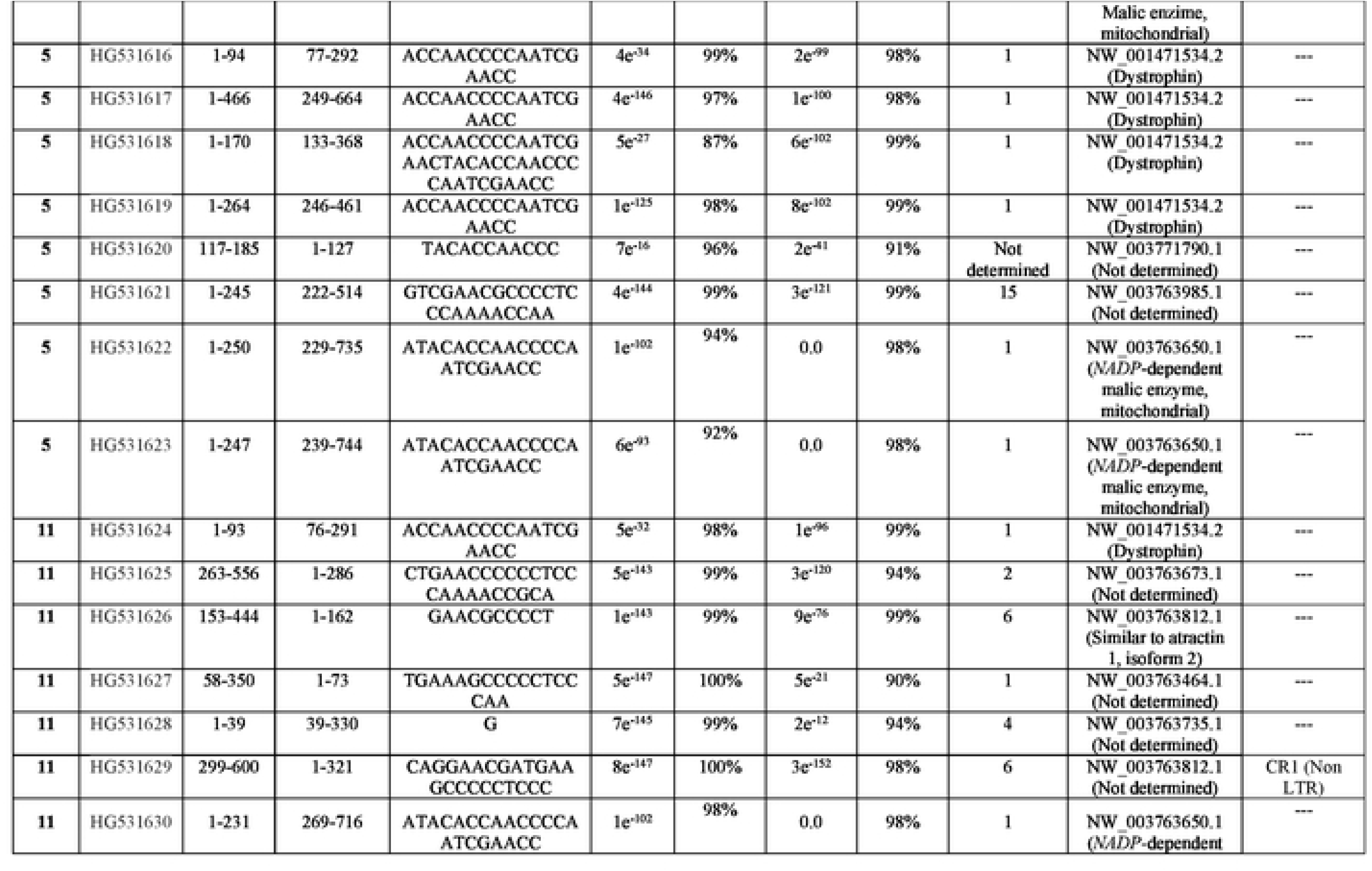
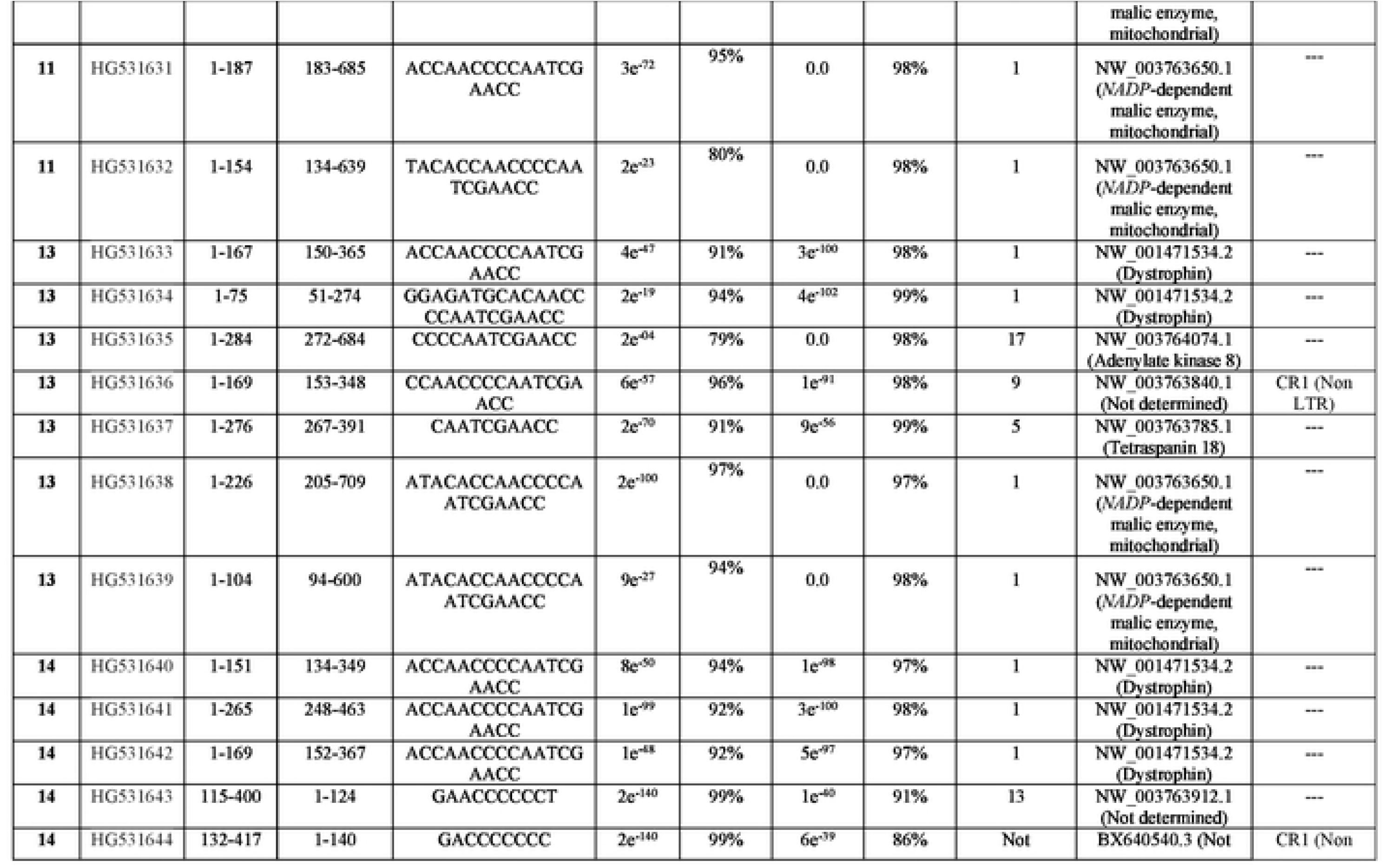
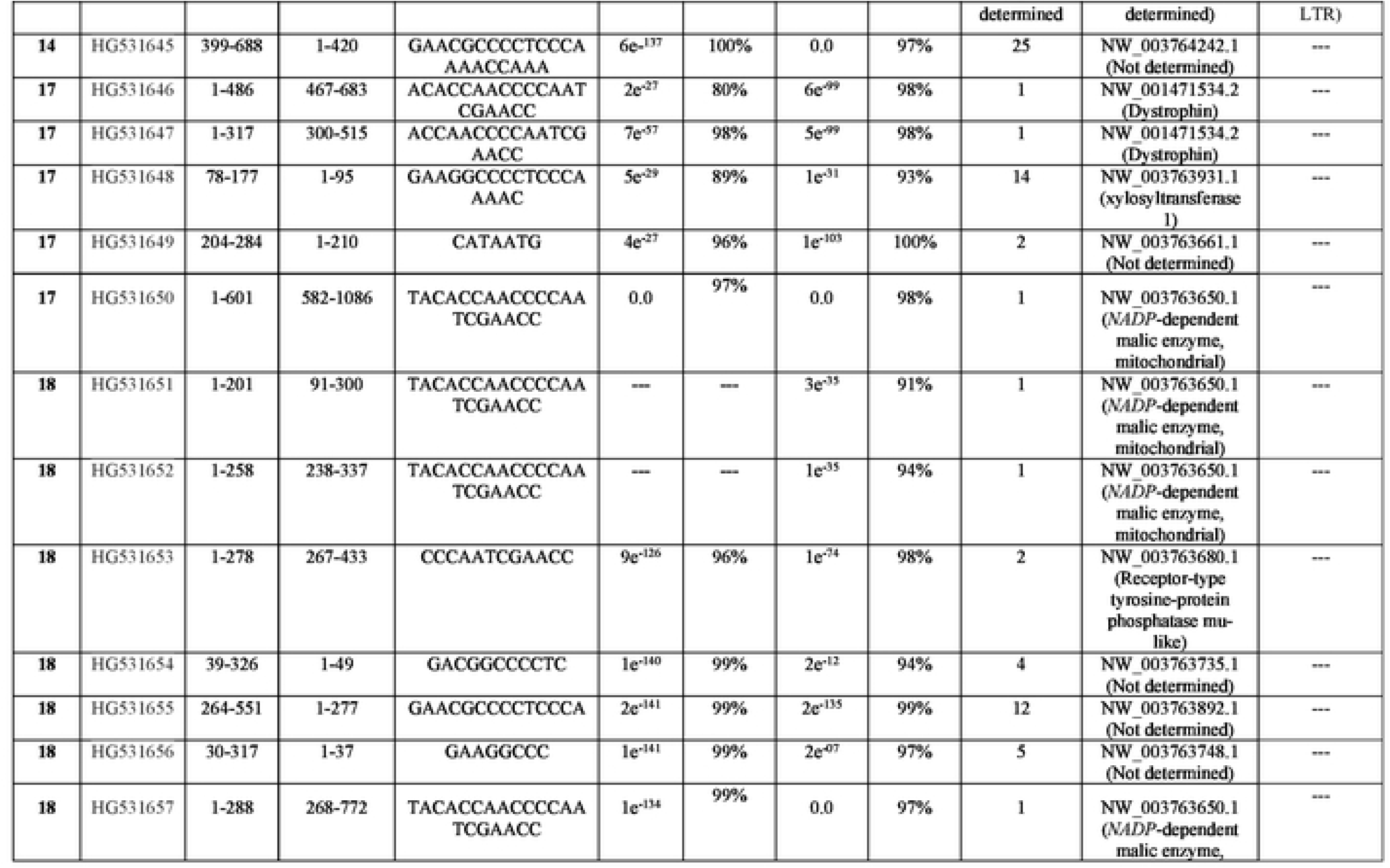
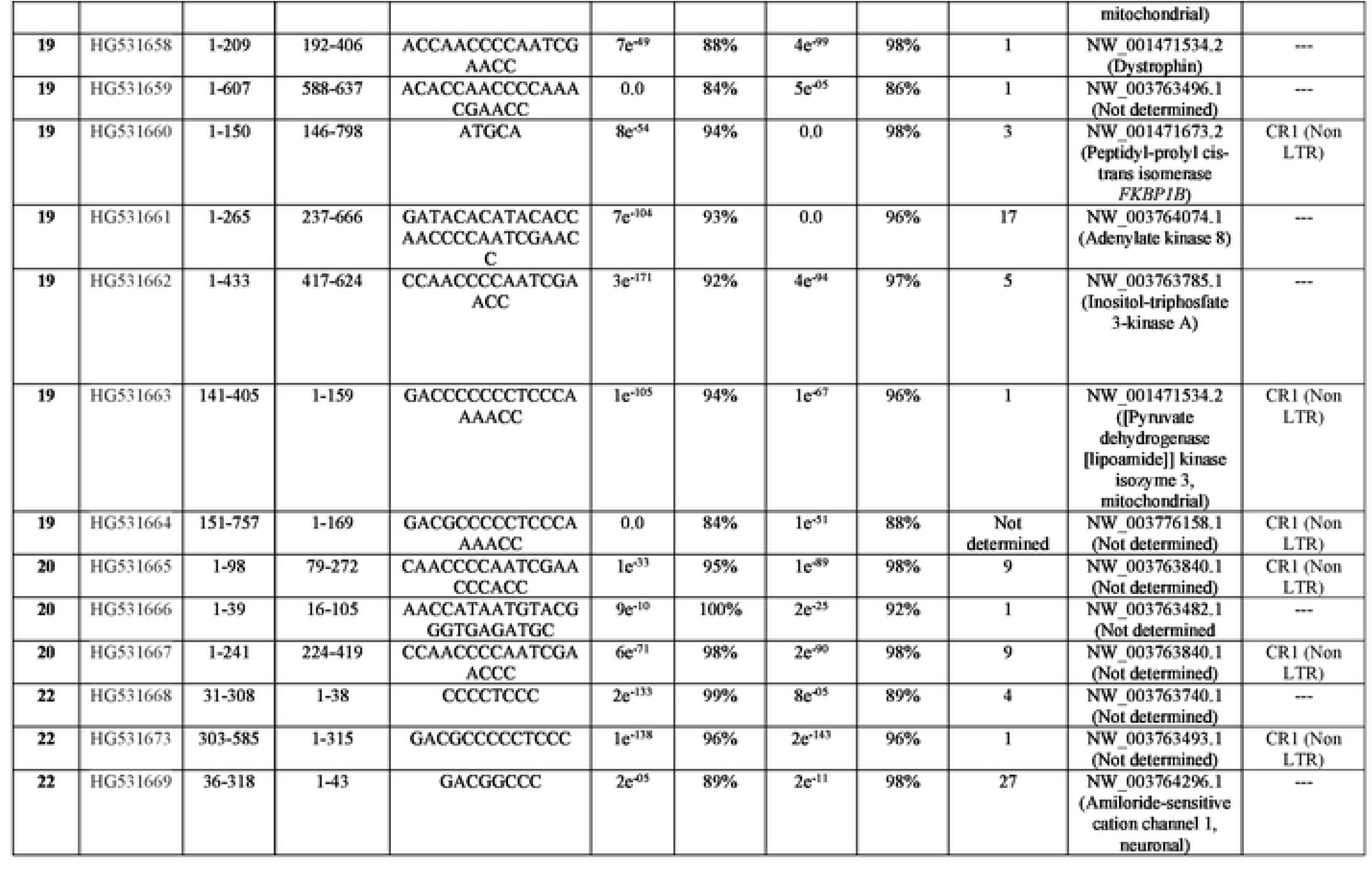
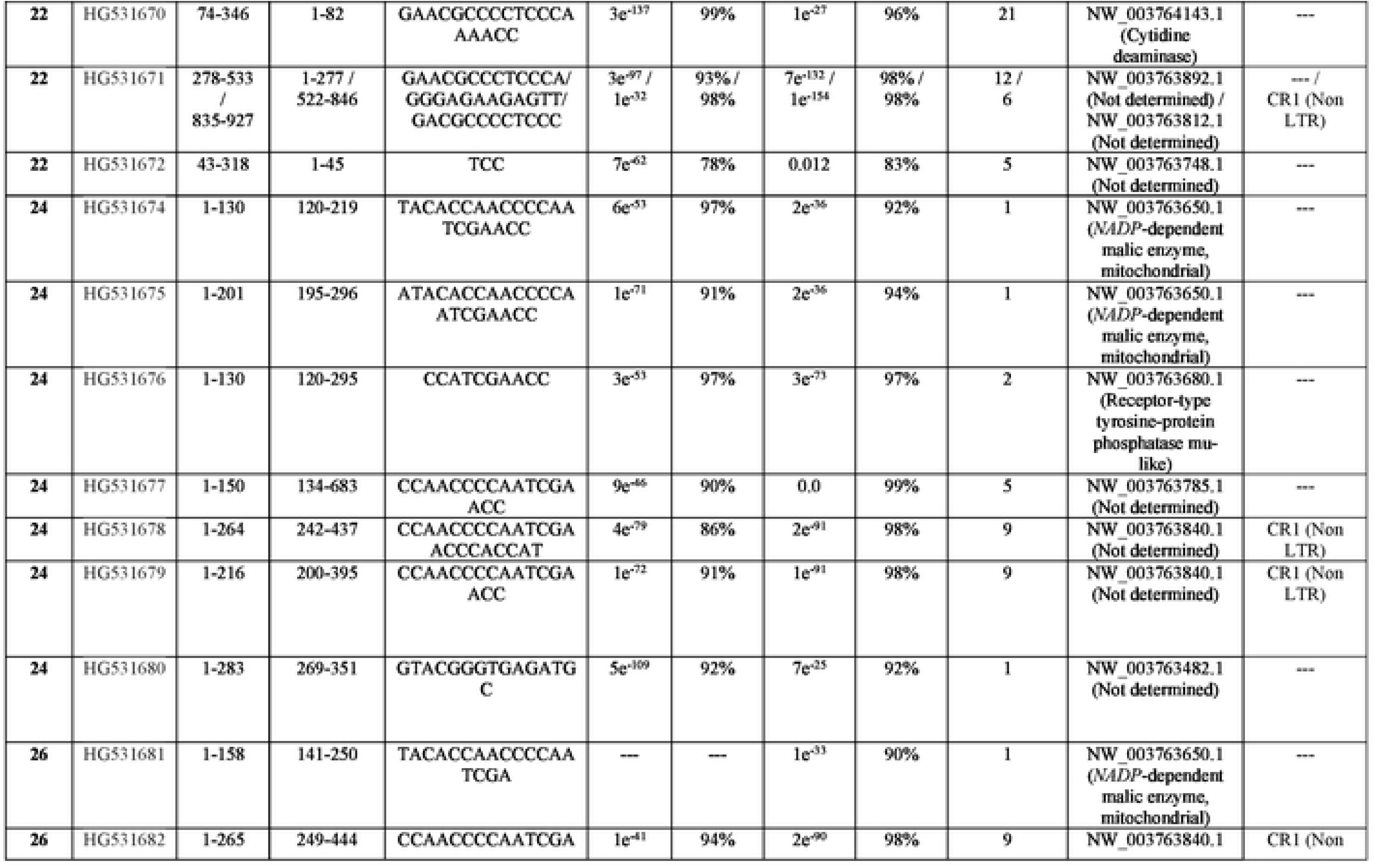
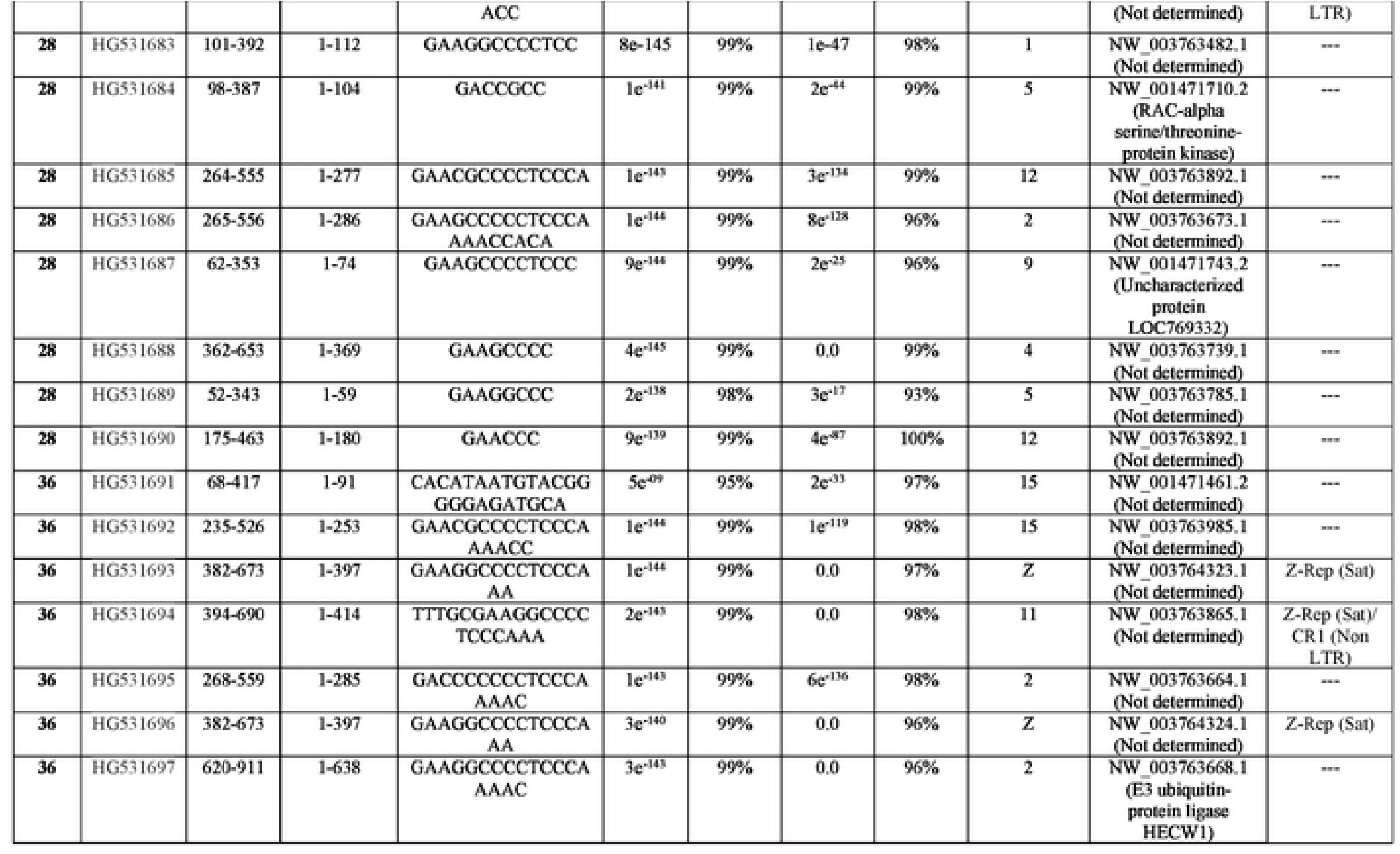
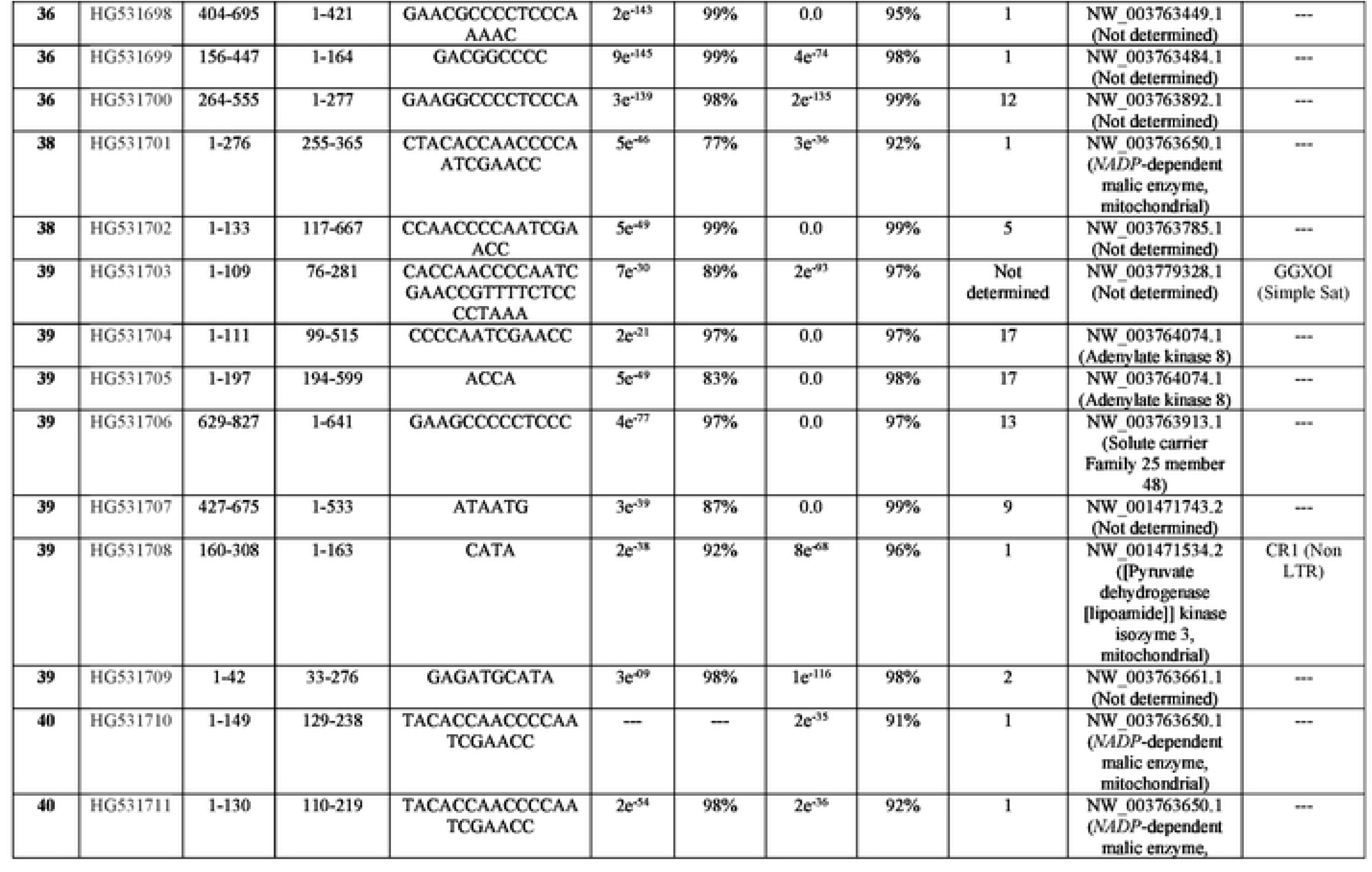
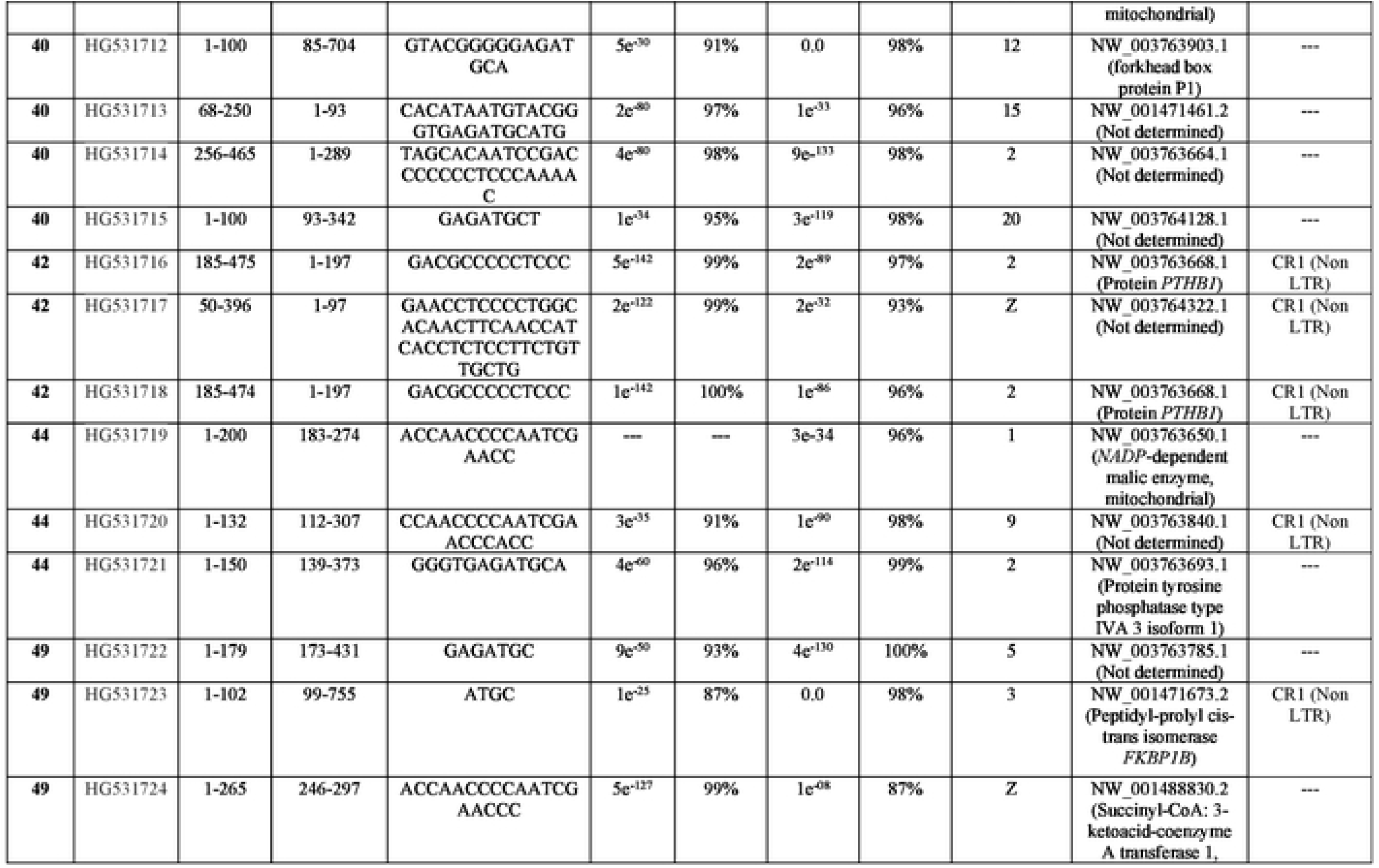
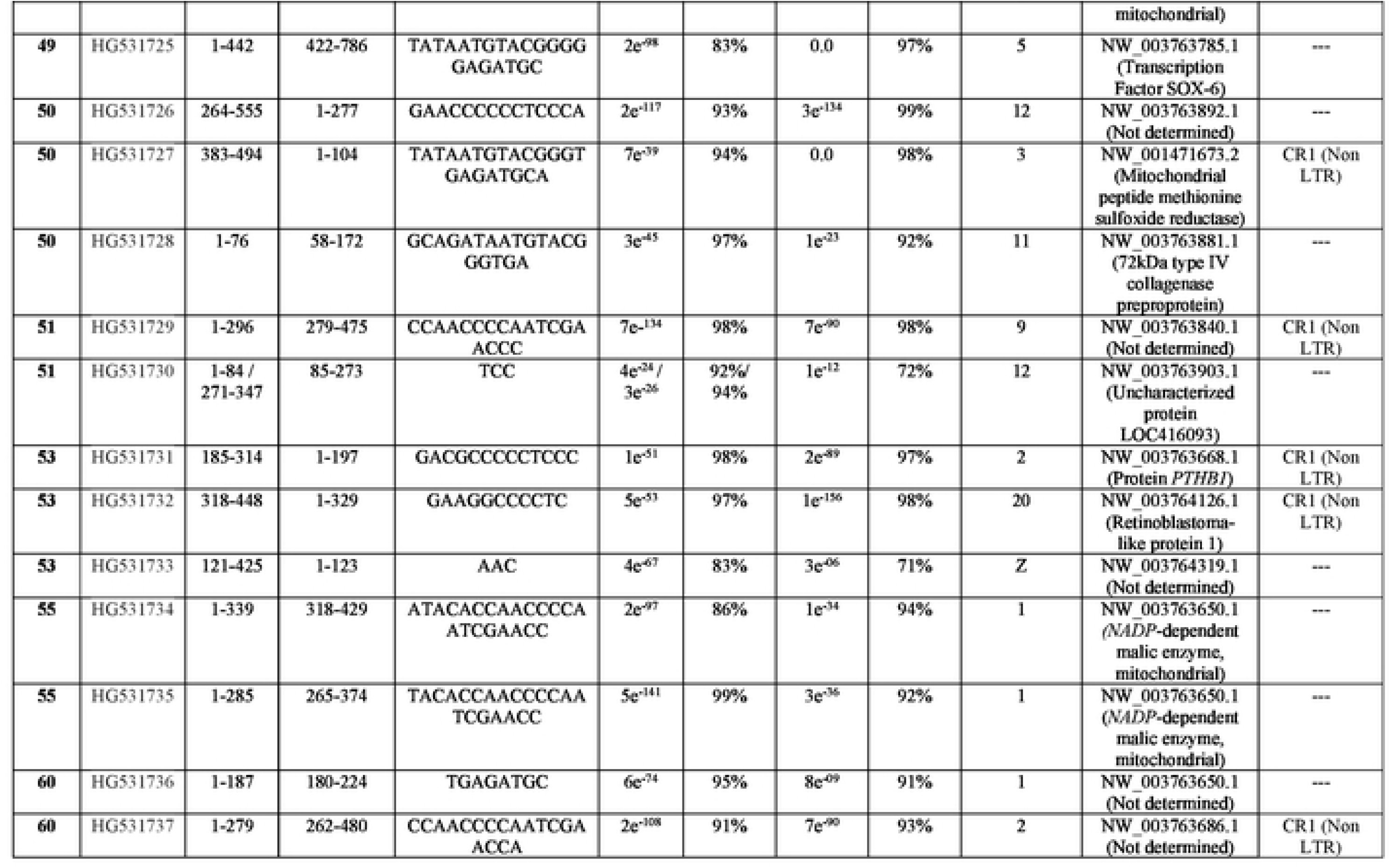
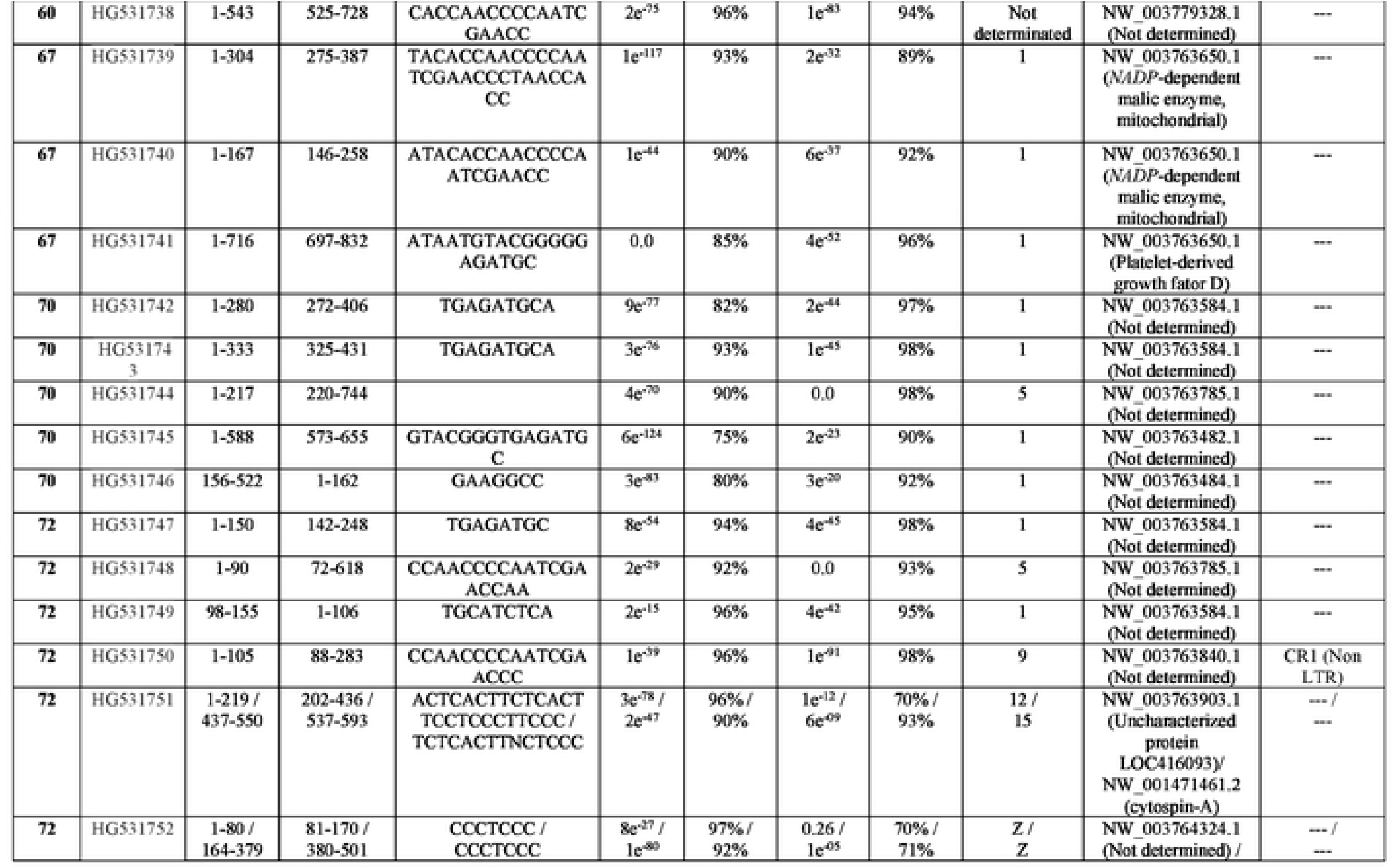
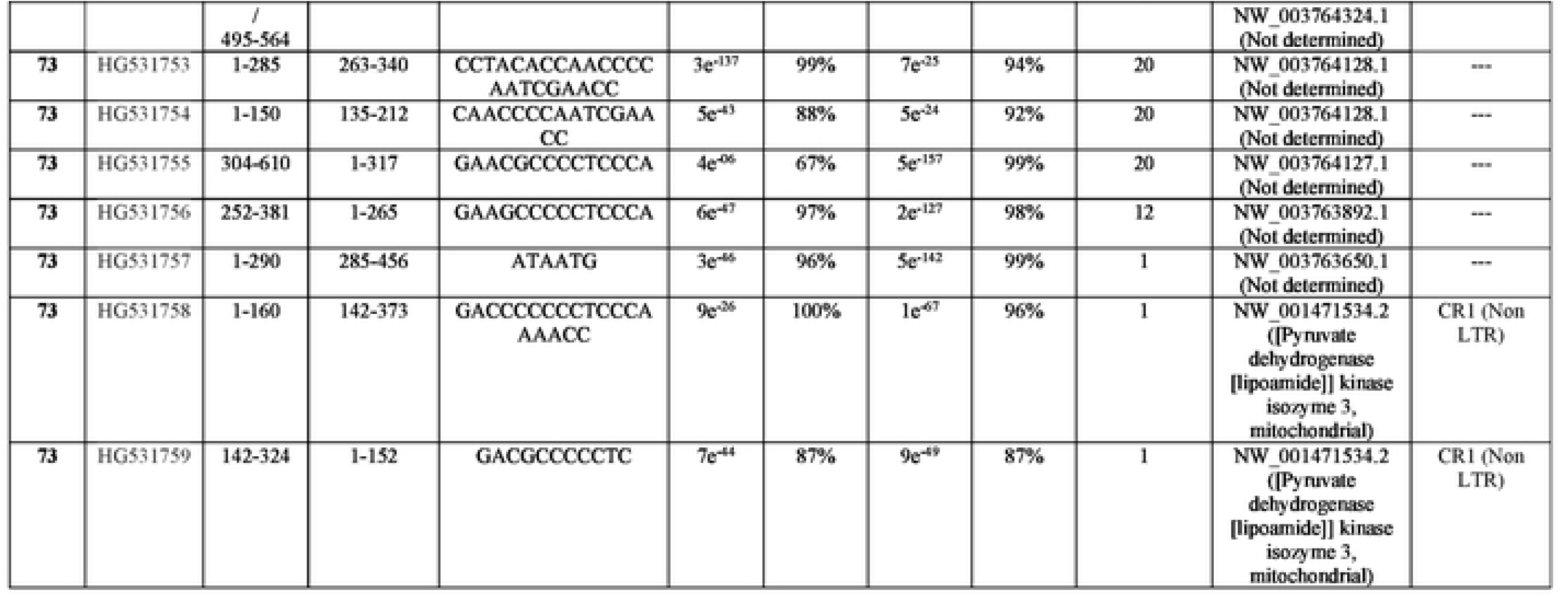
Veritcal transfer of minicircle sequences of kDNA from *Trypanosoma cruzi* into the *Gallus* gallus germ line cells.

**S6 Table.**
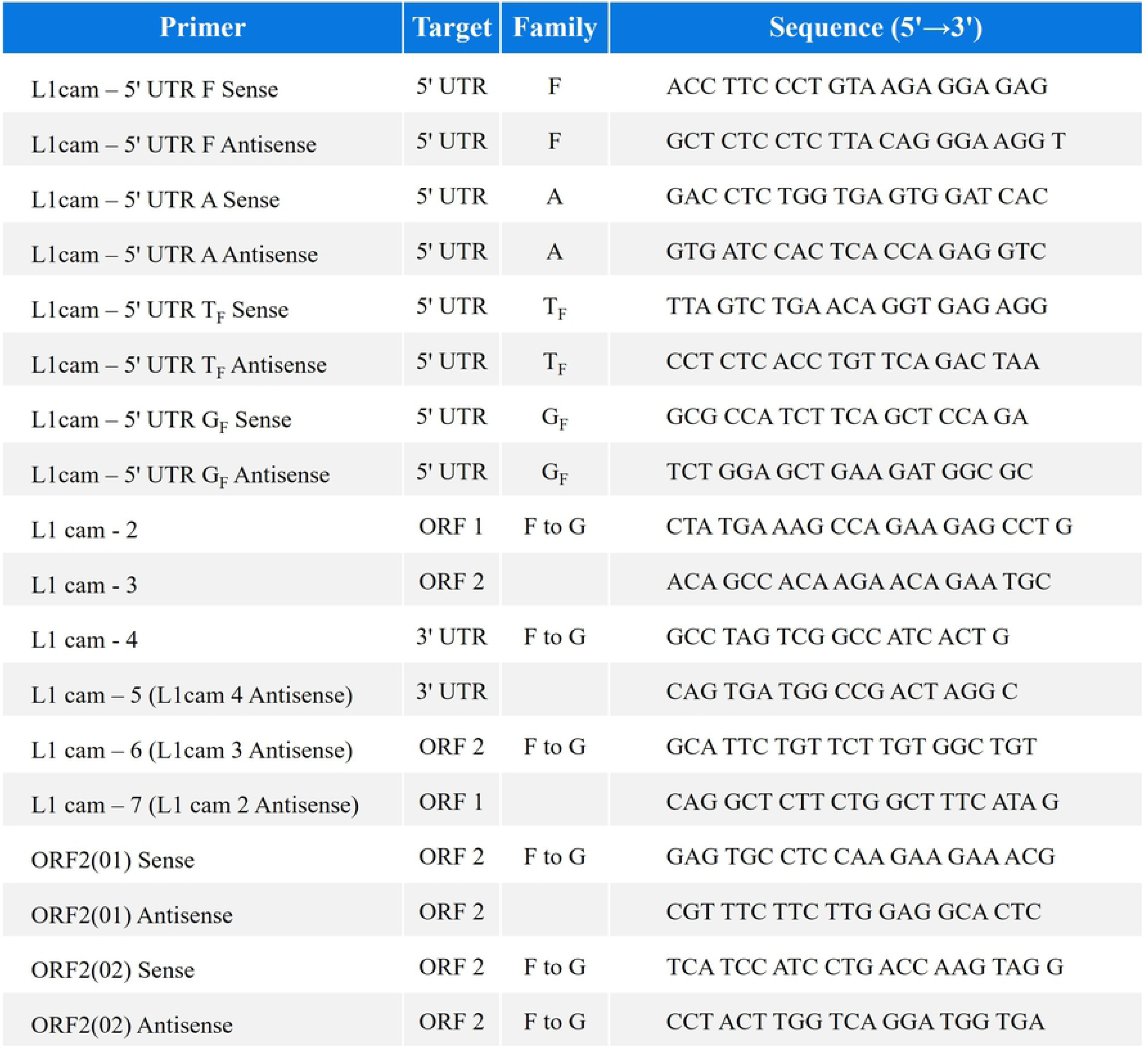
**The *Mus musculus* derived *specific* LINE-1 primers used in combination with the kDNA primers to unravel the lateral transfer of the kDNA into the mammalian genome.** Note the mouse family’s primer sets sense and antisense specific to the untranslated regions (UTR), 5’-3’ orientation to the ORF1 and ORF2, in combination with the kDNA specific primer sets, and Southern hybridization of the amplification products with the radio labeled kCR probe.

**S7 Table.**
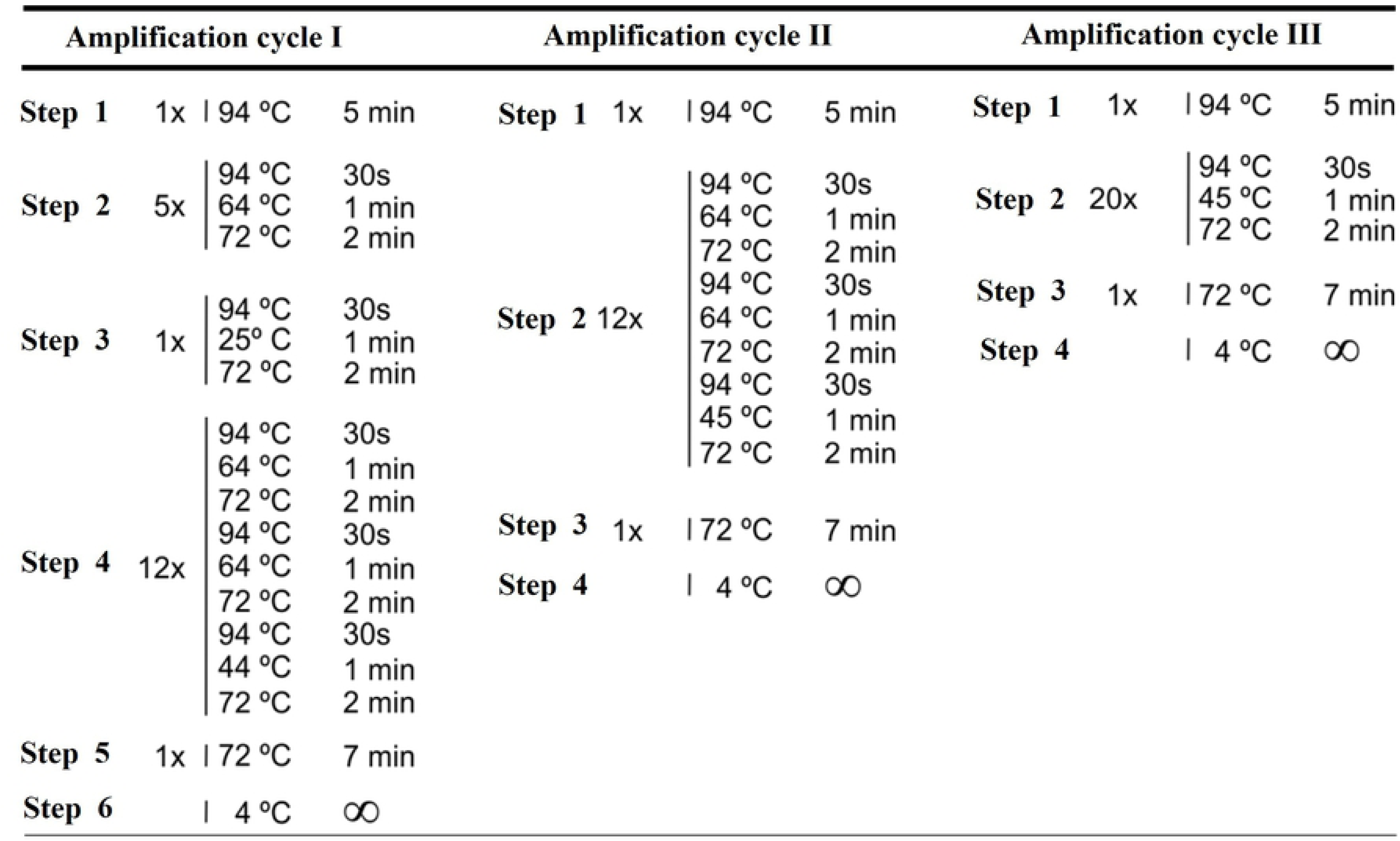
**The *tp*TAIL-PCR amplification cycles anneal temperature °C.** The sense and antisense mouse primer and the kDNA primer sets combination were observed at each cycle of the reaction under variation of the anneal temperature at several steps towards the end of the third cycle showed in the Figure 18. The amplification product showing the specific kCR radio labeled probe hybridization at the third cycle of amplification were clone and sequence.

**S8 Table.**
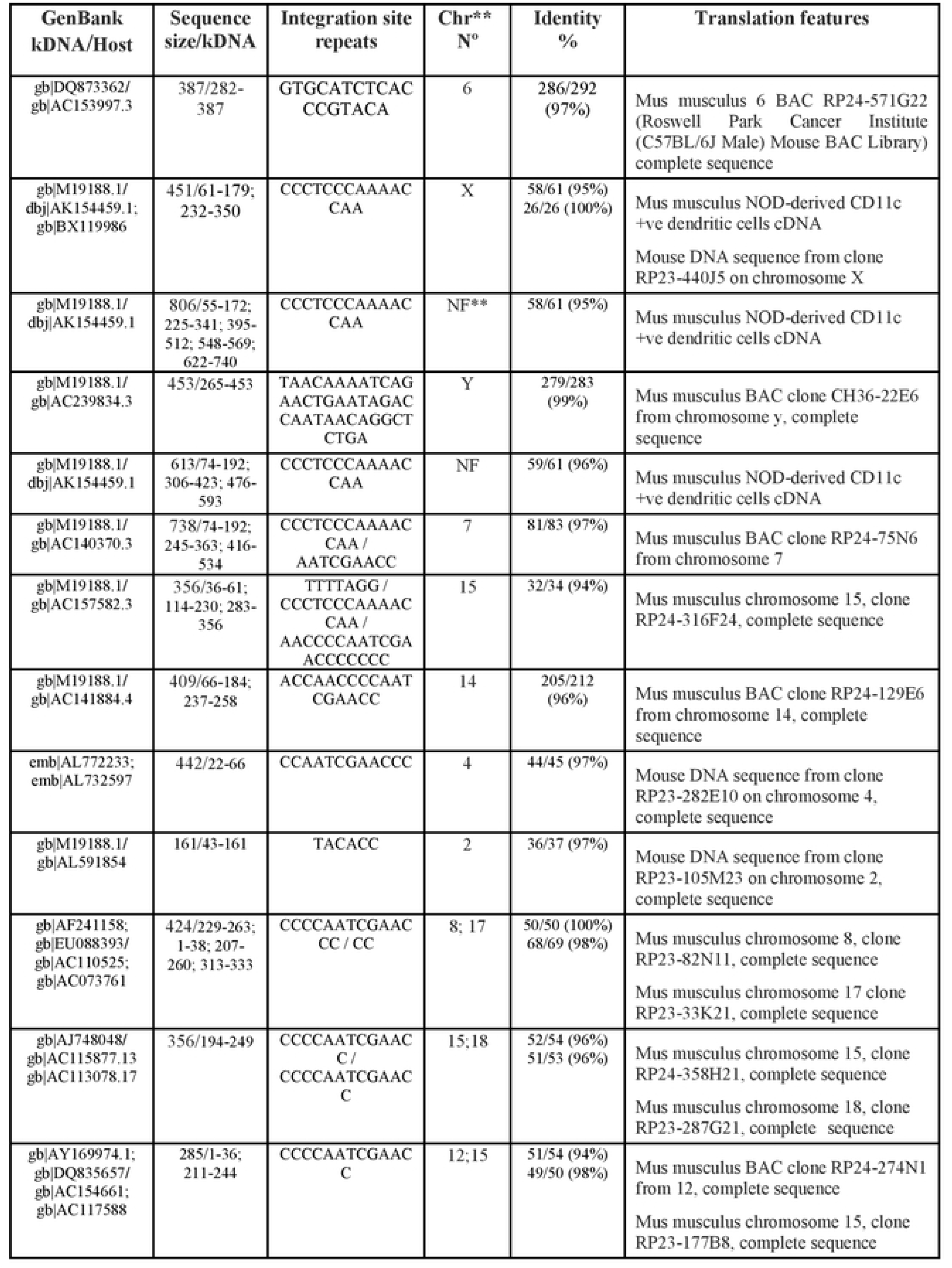
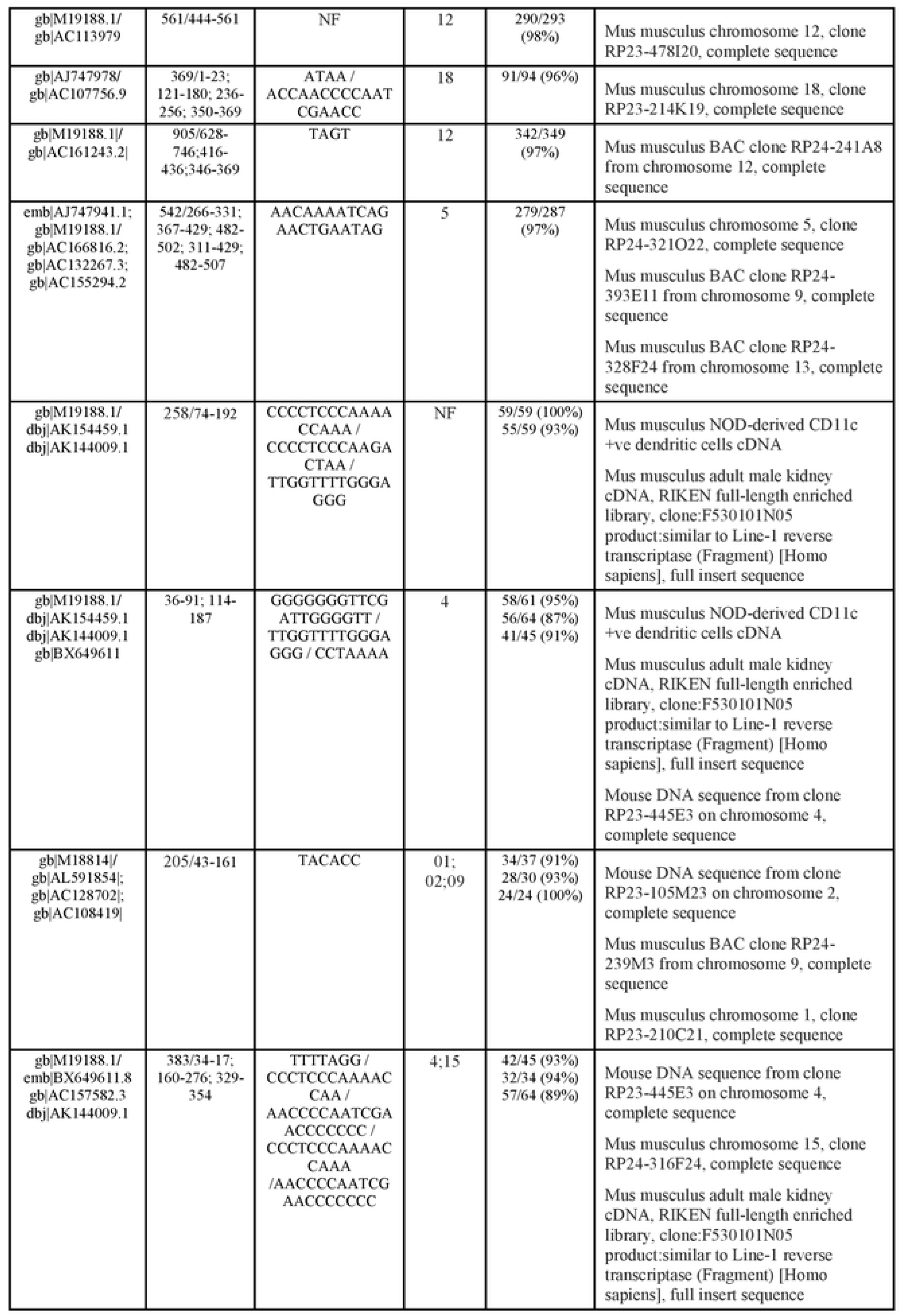
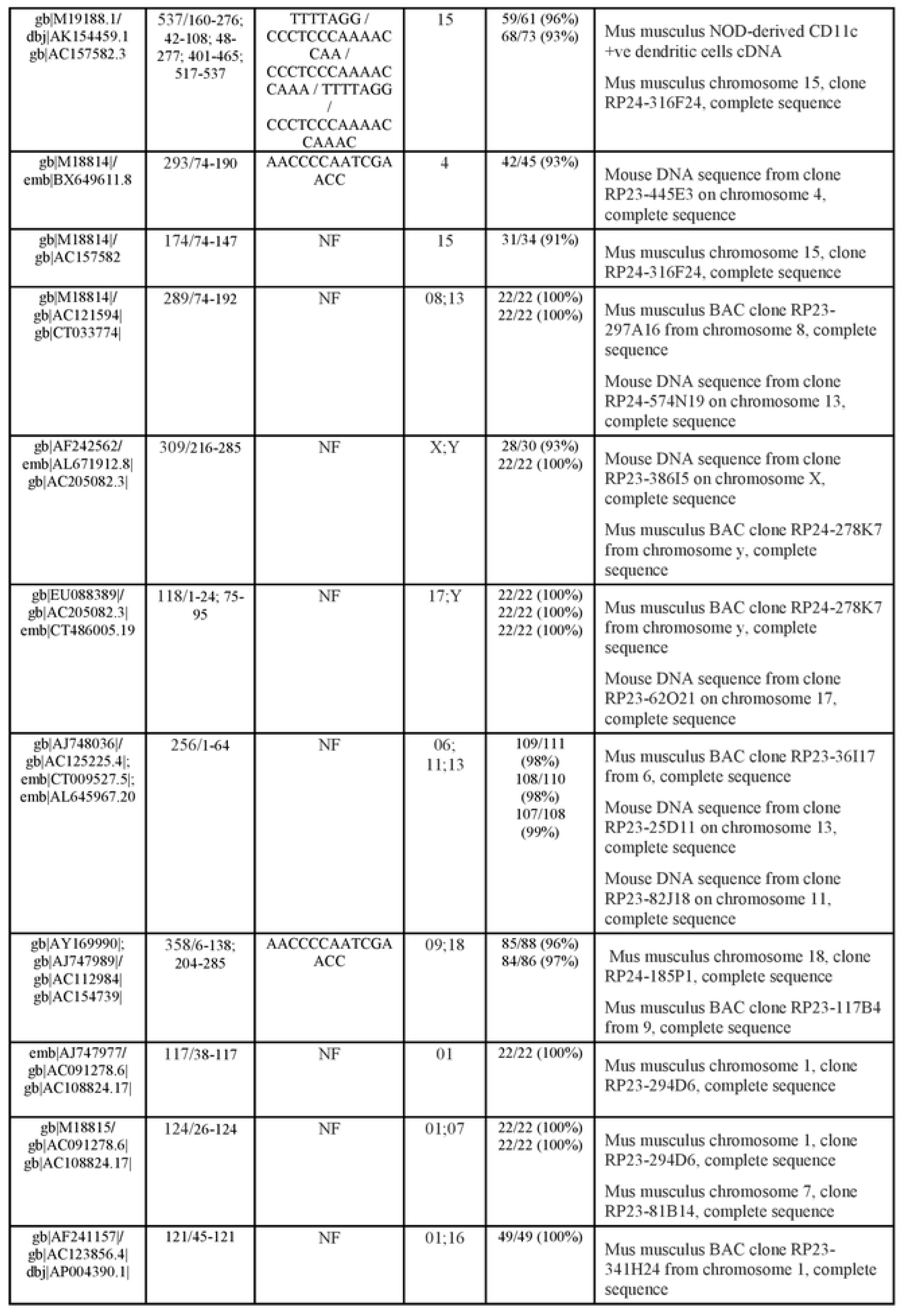
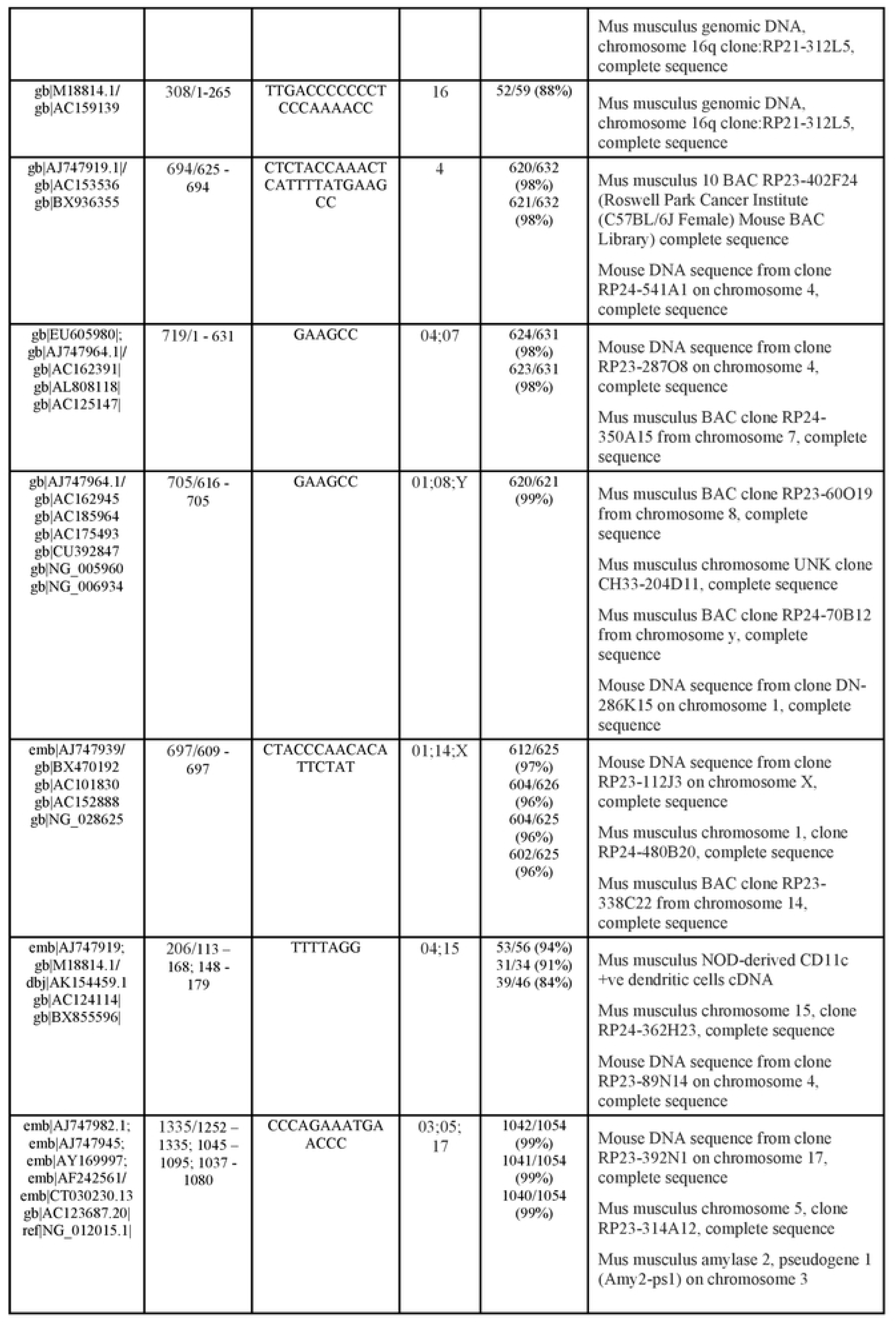
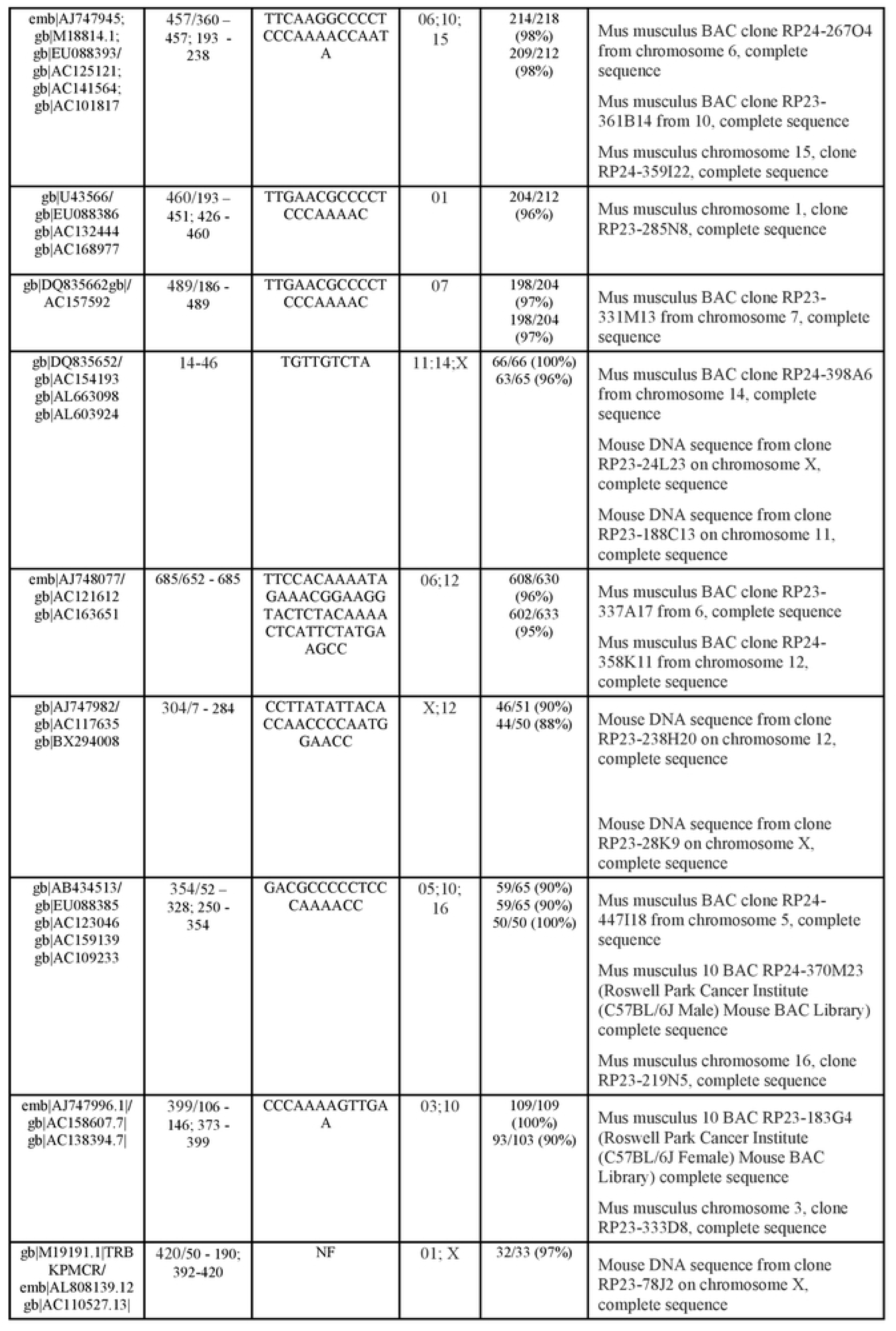
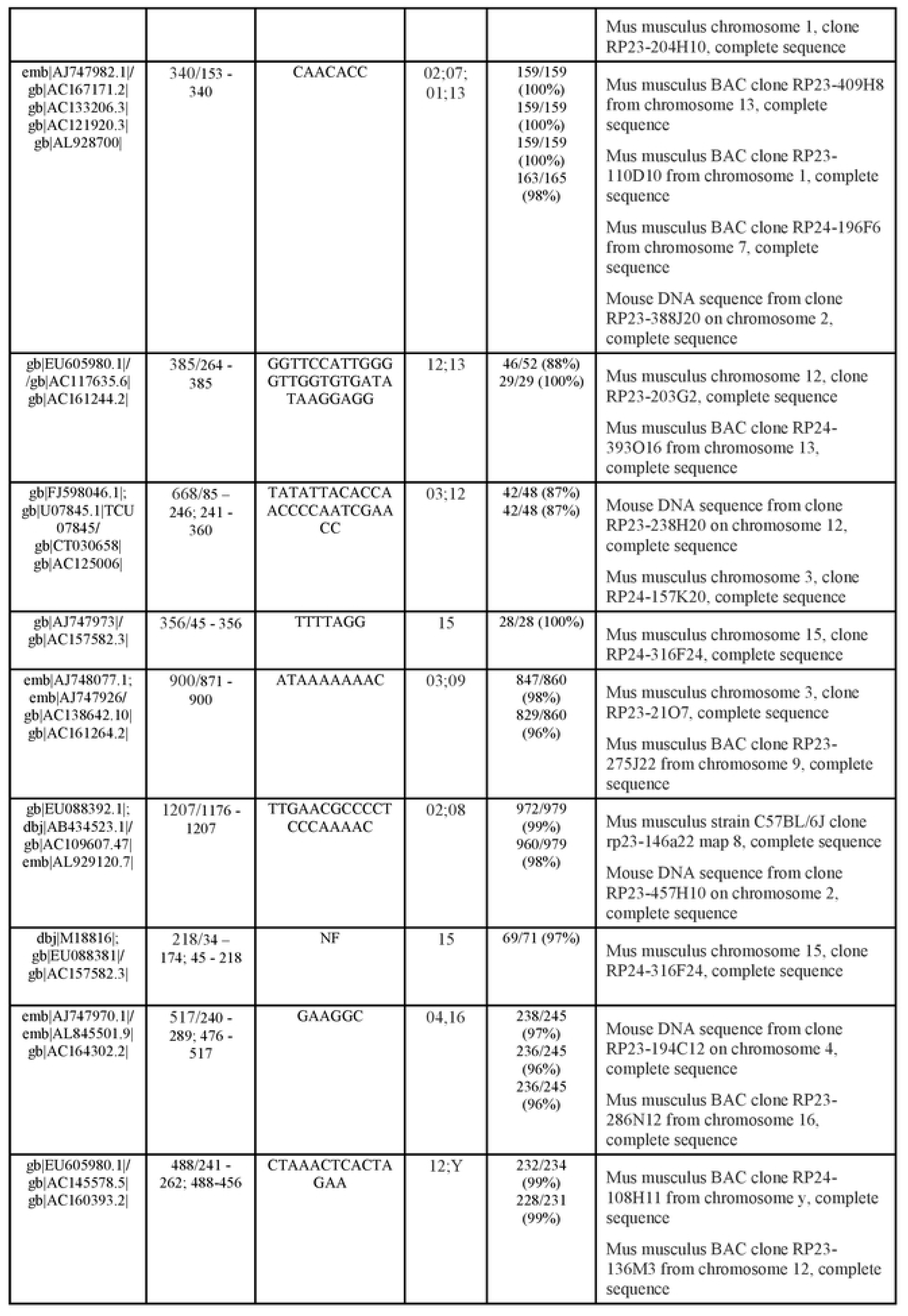
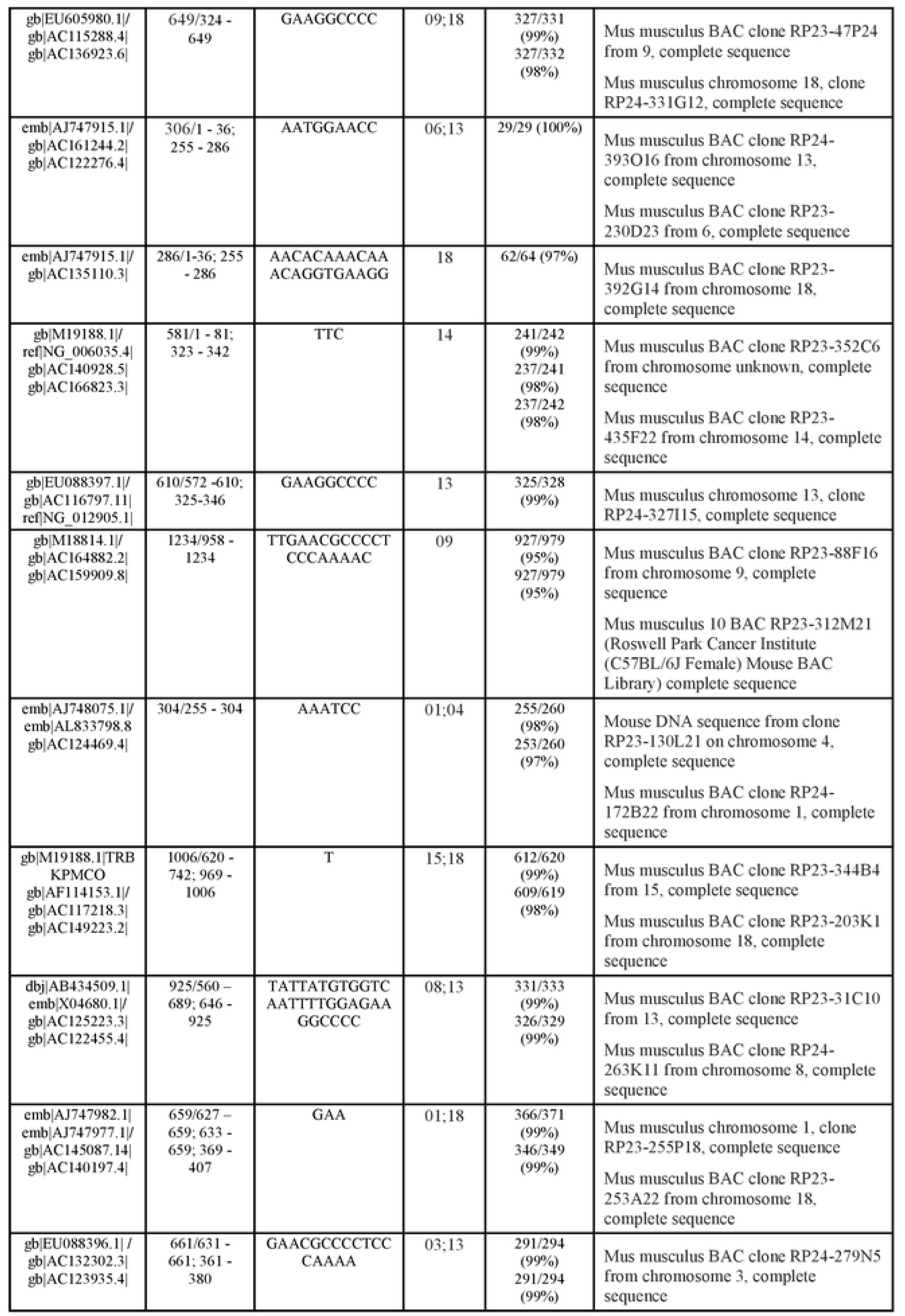
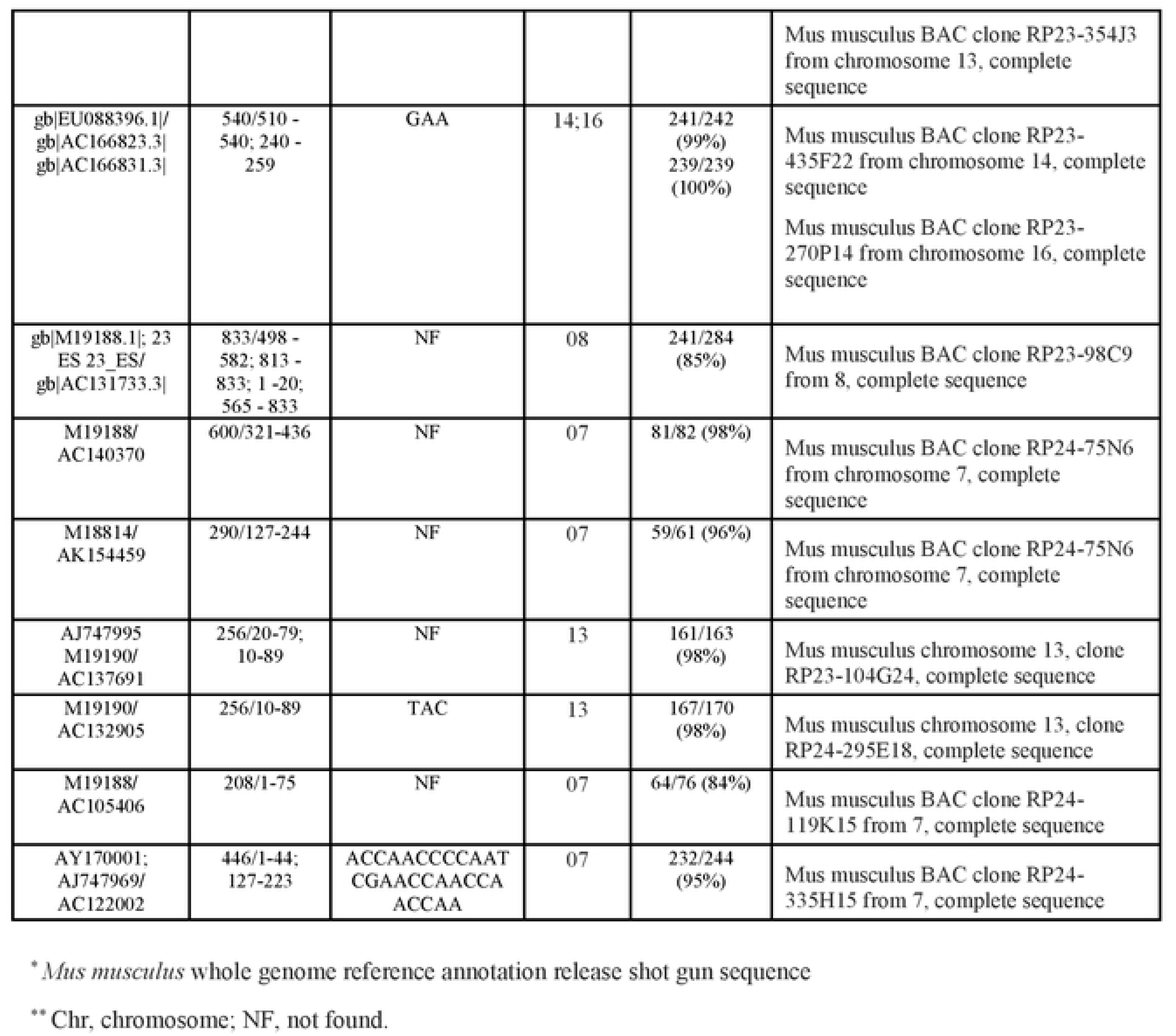
**Mapping of the sites of *Trypanosoma* kDNA integration into the chromosomes of the mouse.** The sequence analysis showed the AC-reach repeats (S2 Figure) intermediate the integration of the *T. cruzi* mitochondrion sequence into the mouse genome, and that 79% of the integrations took place in the family A retrotransposon in the mouse genome.

